# Establishment of local adaptation in partly self-fertilizing populations

**DOI:** 10.1101/2021.04.13.439676

**Authors:** Bogi Trickovic, Sylvain Glémin

## Abstract

Populations often inhabit multiple ecological patches and thus experience divergent selection, which can lead to local adaptation if migration is not strong enough to swamp locally adapted alleles. Conditions for the establishment of a locally advantageous allele have been studied in randomly mating populations. However, many species reproduce, at least partially, through self-fertilization, and how selfing affects local adaptation remains unclear and debated. Using a two-patch branching process formalism, we obtained a closed-form approximation under weak selection for the probability of establishment of a locally advantageous allele (*P*) for arbitrary selfing rate and dominance level, where selection is allowed to act on viability or fecundity, and migration can occur via seed or pollen dispersal. This solution is compared to diffusion approximation and used to investigate the consequences of a shift in a mating system on *P*, and the establishment of protected polymorphism. We find that selfing can either increase or decrease *P*, depending on the patterns of dominance in the two patches, and has conflicting effects on local adaptation. Globally, selfing favors local adaptation when locally advantageous alleles are (partially) recessive, when selection between patches is asymmetrical and when migration occurs through pollen rather than seed dispersal. These results establish a rigorous theoretical background to study heterogeneous selection and local adaptation in partially selfing species.

## 1 Introduction

Most advantageous alleles are lost from populations (Haldane, 1927), but those that escape extinction are ultimately destined to fix, providing that the fitness effect of an allele remains positive over space and time. However, fitness effects can vary across species’ range, so that an allele that is advantageous in one ecological patch might be deleterious in another. Two mutually exclusive outcomes are then possible: Locally advantageous allele is maintained by divergent selection as polymorphism, or the allele with the best overall performance across the patches becomes fixed. Since ranges of essentially all species are spread across heterogeneous environments, it is of some interest to understand the conditions under which these outcomes are materialized. Although previous works investigated these conditions for randomly mating populations in both a deterministic (Bulmer, 1972, Felsenstein, 1976, Levene, 1953, Maynard Smith, 1970) and a stochastic setting (Tomasini and Peischl, 2018, Yeaman and Otto, 2011), conditions for maintenance of local adaptation in partially self-fertilizing populations remain unexplored. Partial selfing is widespread among eukaryotes and well-described in plants (Igic and Kohn, 2006), animals (Jarne and Auld, 2006), algae (Hanschen *et al*., 2018), and fungi (Billiard *et al*., 2012), and considering its effect on spatially heterogeneous selection is especially relevant for at least two reasons. First, partial selfing is more frequently found among sessile or less mobile organisms, for which landscape heterogeneity directly translates into local selective pressures because of the absence of the homogenizing effect of movements across the environment. Second, partial selfing likely influences the conditions for local adaptation by altering the effective population size and effective migration rate, hence the genetic structure of a population, but also by reducing the role of heterozygotes’ fitness in the dynamics of allelic frequencies.

In a single population, selfing exerts two opposing effects on the fixation probability of an advantageous allele (Pollak (1987), Pollak and Sabran (1992),Charlesworth (1992), Caballero and Hill (1992)). Firstly, it increases the rate at which a rare advantageous allele spreads through the population. As the selfing rate increases, homozygotes appear more quickly so that the spread of an advantageous allele becomes increasingly decoupled from the fitness of the heterozygote. In other words, selfing increases the effective dominance coefficient: the rate of spread in a partially selfing population with dominance coefficient *h*, would be equal to the rate of spread in a randomly mating population with dominance coefficient 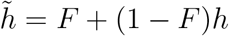 (where *F* is the equilibrium population inbreeding coefficient). Secondly, since alleles making up the progeny are not independently sampled, selfing also reduces the efficacy of selection by reducing the effective population size, 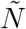, by a factor of 1 + *F* (Gale (1990)). These two opposing effects cancel out when an advantageous allele is codominant (*h* = 0.5), implying that the fixation probability of a codominant advantageous allele is invariant across populations with different selfing rates. If an advantageous allele is partially recessive (*h* < 0.5), then the effect of increased efficacy of selection through increased effective dominance outweighs the reduction in 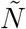 yielding an increase in the establishment probability. Conversely, partially dominant advantageous alleles (*h* > 0.5) are less likely to fix because the reduced efficiency due to lower effective population size trumps increased efficacy of selection due to increased homozygosity. These predictions are altered for male and fecundity selection. Under selfing, no selection occurs on the male function, but the strength of selection on the female function is doubled because two gametes are transmitted for one selected parent (Damgaard, 2000).

The evolutionary dynamic becomes further elaborated in structured populations with divergent selection. The simplest scenario involves two patches with an invading allele being advantageous in one but deleterious in the other patch. The dynamic of local adaptation then further depends on the effect that selfing has on gene flow. Partial selfing reduces the effective haploid migration rate due to the reduction in the number of female gametes that can be sired by immigrating male gametes, for individuals in the selfing population will have already fertilized themselves to a large extent. This reduction in gene flow has been proposed to prevent the spread of maladaptive alleles from neighboring populations and thus to promote local adaptation (Linhart and Grant, 1996), but two meta-analyses of reciprocal transplant experiments reported the absence of correlation between local adaptation and the mating system (Hereford, 2010, Leimu and Fischer, 2008). However, it is not immediately clear how the interplay between the mating system, migration, and selection modes affects the propensity for local adaptation. When local reproduction is panmictic, maintenance of local polymorphism roughly requires that local selection is stronger than migration. On the one hand, selfing reduces effective migration. On the other hand, it may either increase or decrease the efficacy of local selection, depending on the balance between the effect of increasing effective dominance and increasing genetic drift. In addition, in diploids, conditions where alleles are (partially) dominant when locally beneficial and (partially) recessive when locally deleterious are especially favorable to the maintenance of polymorphism (Yeaman and Otto, 2011). By unmasking recessive alleles, selfing is expected to reduce the range of applicability of those conditions. Overall, we still lack correct theoretical predictions, which makes it difficult to interpret the empirical results such as in Leimu and Fischer (2008) and Hereford (2010).

In the present work, we extend the results of previous theoretical models of local adaptation to partially self-fertilizing populations. We first derive a closed-form approximation for the probability that a locally advantageous allele escapes extinction. We then use this result to examine how previously described effects of selfing affect (1) the probability that allele escapes extinction, (2) conditions required for the establishment of protected polymorphism, and (3) the dependence of these properties on the mode of selection and migration.

## 2 Model

We work with a hermaphroditic population that is divided into two patches connected by bidirectional migration. Migration can occur during both the haploid and the diploid phase. In the following, we will use the example of a plant life cycle, but results can readily be transposed to other life cycles. Thus, only male migration, pollen dispersal, occurs during the haploid phase, and diploid migration corresponds to seed dispersal. Fitness is controlled by a single biallelic locus, with allele *A* favored in the first patch and allele *a* favored in the other patch. These patches are referred to as favored and disfavored patch, respectively. The life cycle of the modeled organism is composed of the following sequential stages: (1) male and female meiosis leading to to the production of male gametophyte (pollen containing sperm cell) and female gametophyte (embryo sac, containing egg cell), (2) pollen dispersal, (3) mating (including selfing) and seed formation, (4) seed dispersal, and (5) seed establishment and development, which yields adults of the next generation. To make the model general, we allow selection to operate on viability and on male and female fecundity. More precisely, adults in patch *i* with genotype *k* (*AA, Aa*, or *aa*) produce 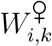 female and 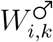 male gametophytes. We do not consider haploid selection, but this could readily be included in the model setting. Finally, each offspring with genotype *k* survives to maturity with probability *V_i,k_*. The population selfing rate, *S*, is defined as the proportion of individuals in the population that self-fertilize and is equal across patches. Following previous works (ex: Hartfield and Glémin, 2016, Hössjer and Tyvand 2016) we assume that the population reaches equilibrium for inbreeding coefficient *F* and genotypic composition on a much shorter time-scale than the change in allelic frequencies (separation of time scale argument). Within each patch, we can thus write genotype frequencies directly as a function of allelic frequencies and *F*. If selection is not too strong, we can also use the neutral expectation: *F* = *S*/(2 − *S*). This assumption is relaxed in simulations. Mathematical symbols and their meaning are outlined in Table 1.

**Table 1:**
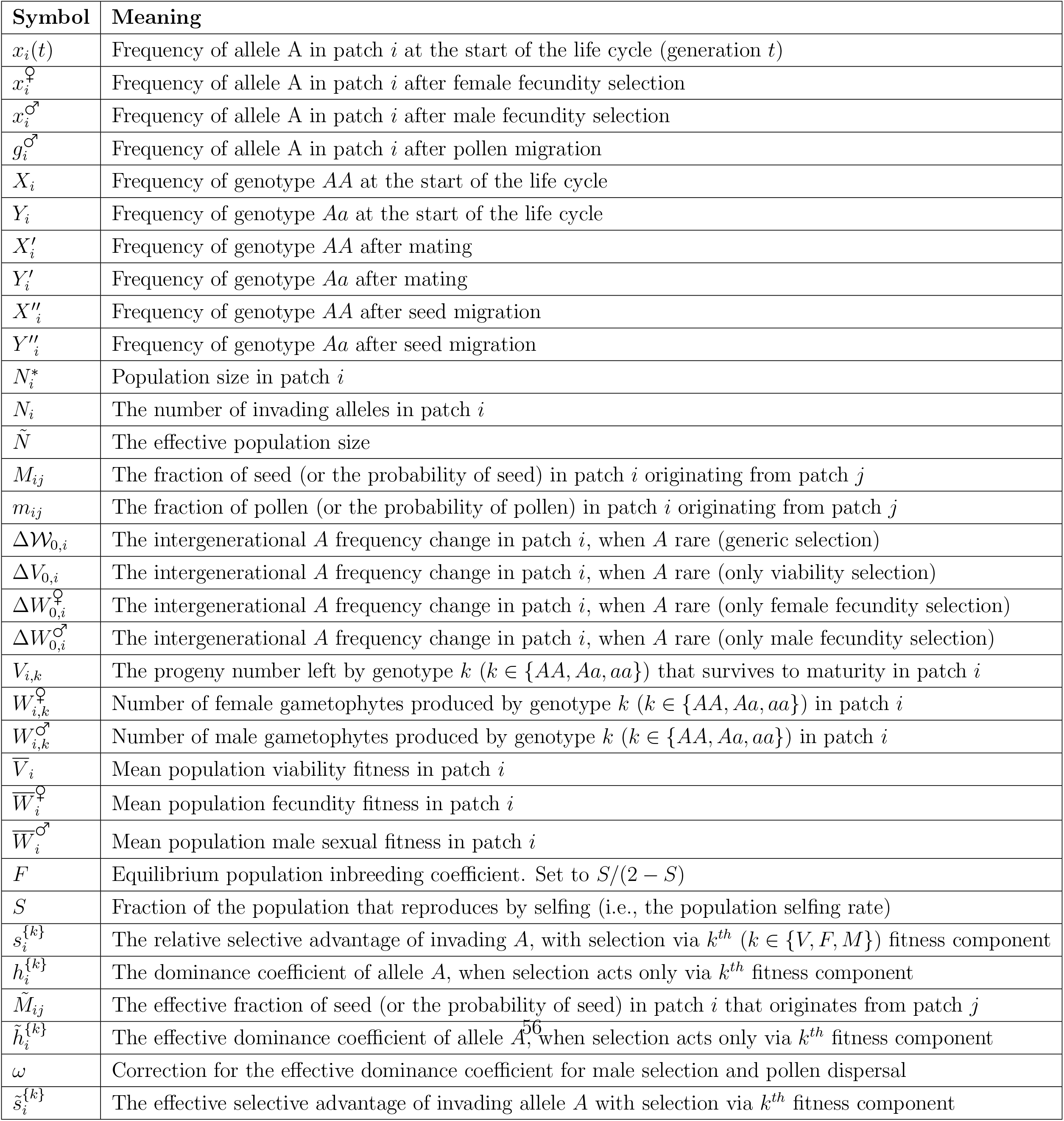
List of symbols used in the text and their meaning.

### 2.1 Deriving the difference equation for allele frequency

Using the preceding definitions, one can derive the difference equations for allele frequencies in each patch. Let *X_i_* and *Y_i_* be the frequencies of genotypes *AA* and *Aa*, and *x_i_* be the frequency of allele *A* in *i^th^* patch. Our system has two degrees of freedom, given that there are three genotype frequencies and they must add up to unity. Thus, we keep track of frequencies of genotypes *AA* and *Aa*, while the frequency of *aa* genotype is by definition 1 − *X_i_* − *Y_i_*. Adult genotype frequencies in generation *t* in *i^th^* patch are:

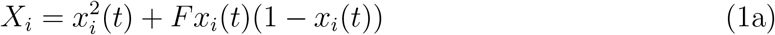

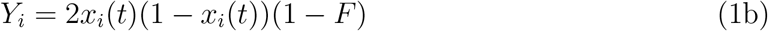

Adult genotypes may produce different number of male and female gametophytes. Let 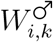 and 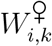 be the number of male and female gametophytes that an adult with genotype *k* (*AA, Aa* and *aa*) produces in patch *i*. Thus, the frequency of female and male gametophyte *A* in *i^th^* patch are, respectively:

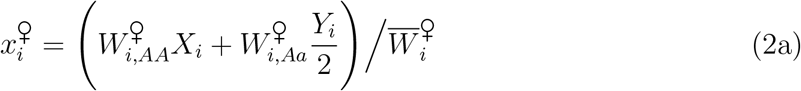

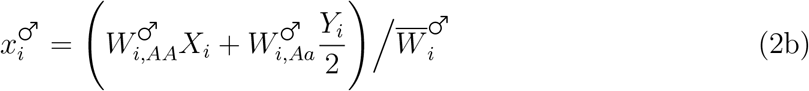

where 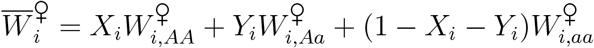. and 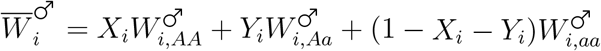. After pollen migration, the frequency of allele *A* in pollen in *i^th^* patch is:

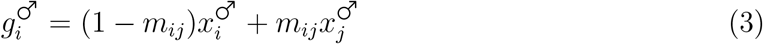

where *m_ij_* is the fraction of pollen in *i^th^* patch that come from *j^th^* patch. Pollen dispersal is followed by mating, which yields offspring genotypes:

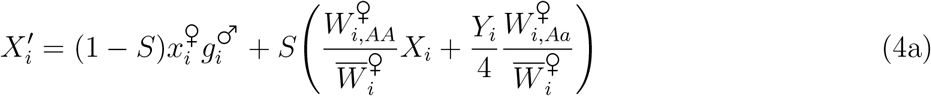

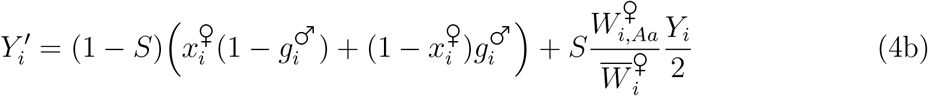

Seed migration changes the genotype frequency to:

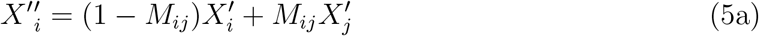

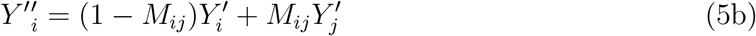

where *M_ij_* is the fraction of seed in patch *i* originating from patch *j*. Genotype *j* in patch *i* has probability *V_i,j_* to survive to maturity, so the frequency of allele *A* in the generation *t* + 1 is:

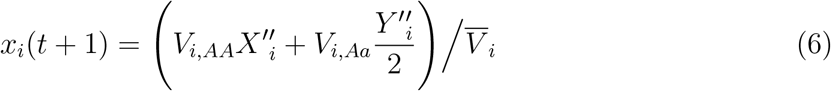

where 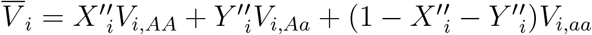.

Equation (6) is our difference equation that expresses the frequency of allele *A* in the next generation in terms of its frequency in the current generation.

### 2.2 Analysis of the deterministic model

We first analyze the deterministic model that gives the conditions for a protected polymorphism. This extends results of Bulmer (1972) to partial selfing and more general forms of selection. Given that investigation of the consequences of an arbitrary selection, regime is prohibitively complicated, the proceeding analysis is restricted to a few special cases. We are ultimately interested in three selection scenarios: (1) when selection operates on differential survival to maturity and when selection operates on (2) female and (3) male fecundity. For each of these scenarios, we first consider seed migration only, as in Bulmer (1972) initial model (also as in Yeaman and Otto, 2011). In addition to the comparison with previous models, it allows analyzing the effect of selfing on selection only, as selfing does not affect seed migration. Then, we consider pollen migration thus analyzing the joint effect of selfing on both selection and migration. In total, we examine six categories of scenarios.

We use the standard approach to obtain conditions for stable polymorphism by considering the conditions for which both monomorphic equilibria are unstable. Let both patches be fixed for allele *a* and allele *A* acts as an invader. Equivalent results can be obtained by considering patches fixed for allele *A* and *a* acting as the invader. The logic of derivation is the same in all cases, wherein we linearize equations (6) around 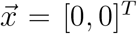 to obtain the system of linear difference equations, where **J** is the associated Jacobian. While **J** captures the invasion dynamics for an arbitrary mode of selection and migration, it is difficult to analyze, so we focus on special population genetic cases outlined above. Each of these scenarios will have an associated Jacobian **J**^{*k*}^:

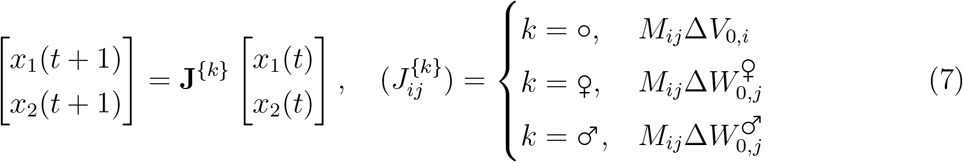

Terms Δ*V*_0,i_, 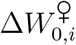, and 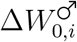 are the intergenerational change in frequency of *A* in *i^th^* patch due to selection on a particular fitness component, and *M_ij_* accounts for migration between patches, with *M_i,i_* = 1 − *M_i,j_*, with *j* ≠ *i*. In the case of viability selection, the frequency change of *A* in the patch 1 reflects either the contribution of alleles *A* that stay in patch 1 (1 − *M*_12_) and survives to adulthood (Δ*V*_0,1_), or the contribution of alleles *A* that migrate from patch 2 (*M*_12_) and then survive to adulthood (Δ*V*_0,1_). Similar logic holds for female and male fecundity selection but off-diagonal selection terms are exchanged because selection occurs prior to migration: note the *j* indices in Δ*W* terms instead of *i* in the Δ*V* term in equation (7). Selection terms for the three scenarios are obtained by keeping constant the two fitness components not involved in the scenarios in the general recursion equations. Thus, under seed migration the three selection terms are:

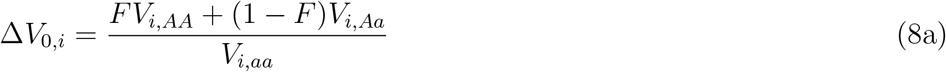

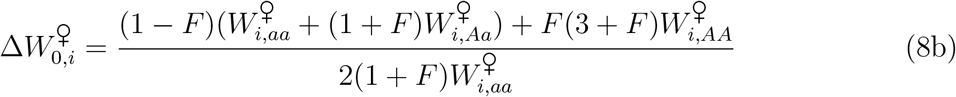

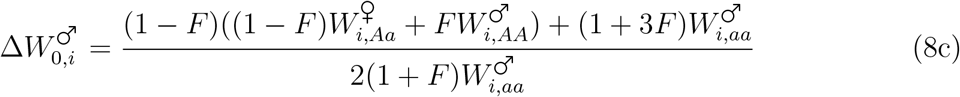

These expressions can be made more intuitive by parameterizing fitnesses relative to genotype *aa*, where 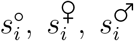 are the reproductive advantage of *AA* homozygote relative to *aa* homozygote when selection acts on viability, female, and male fecundity, respectively. Parameters 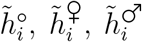 represent the effective dominance of the heterozygote in *i^th^* patch for the three respective selection modes. These are composite parameters that depend on *F* and the actual dominance and are introduced to capture the fact that selfing increases the effective dominance of invading heterozygote due to contribution of mutant homozygotes to the invasion process. For the three cases of viability, female, and male fecundity effective dominances are respectively:

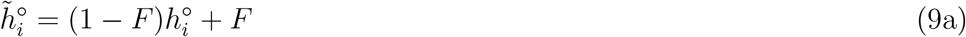

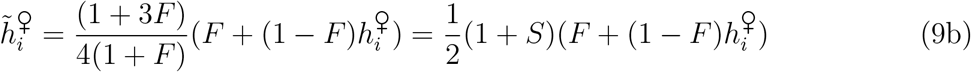

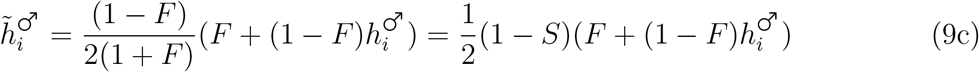

These forms capture the fact that selfing increases the effective dominance of invading mutants: the invader has dominance *h_i_* when it invades in a heterozygous form (that is, 1 − *F* fraction of time), and dominance of one when it invades in a homozygous form (that is, *F* fraction of time). Selfing also affects the global selection intensity on male and female fecundity. The 1/2 factor captures the fact that selection acts only on half of the gametes produced by each genotype, the 1 − *S* factor that selection on male fecundity only operates under outcrossing and the 1 + *S* factor that under selfing an individual contribute two alleles through seed production (female fitness). Taken together:

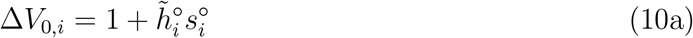

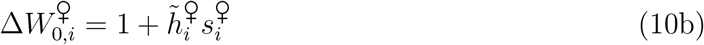

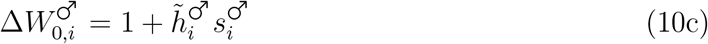

The fitness of the mutant always take the same form as in a haploid model with an effective selective advantage 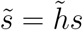.

The preceding approach can be readily extended to the selective scenarios where migration occurs solely via pollen. One can still use the Jacobian in (7), albeit with the modified migration rate to account for the facts that (a) pollen is haploid and thus do not contribute in the same way as diploid seed to the gene pool of the next generation, and (b) that in all three cases migration operates prior to selection rather than after:

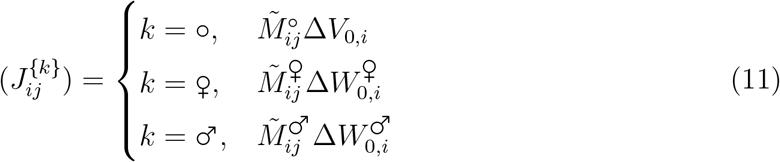

Selection terms are still parameterized according to equations (10a–10c), but effective migration is given by:

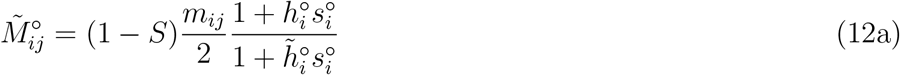

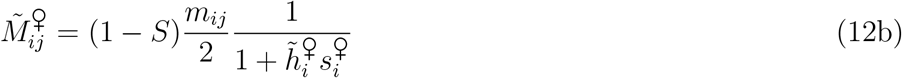

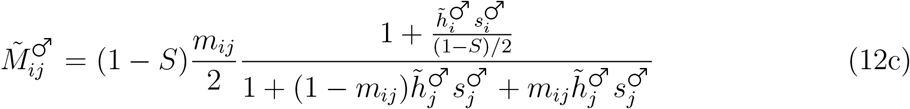

Viability effective migration rate (12a) can be intuitively understood by noting that 1 − *S* corresponds to the fraction of pollen that is exchanged between the patches, *m_ij_*/2 corrects for the fact that haploid phase contributes two-fold less alleles to the gene pool of the next generation relative to the diploid progeny dispersal, and the relative fitness ratio 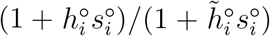 accounts for the fold increase in invader frequency in the patch of origin due to the inhibitory effect of selfing on gene flow. Similar interpretation holds for (12b) and male fecundity selection (12c), except that fitness ratio now corrects for selection acting prior to migration. Note that for weak selection, the three effective migration rates reduce to 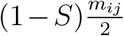. The only complication is that when selection acts on the male fitness component, one has to modify the effective dominance along with the effective migration rate. More precisely, 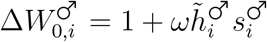, where *ω*:

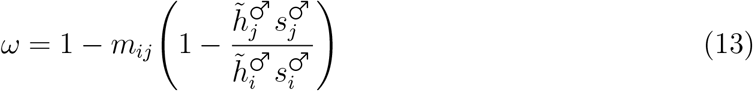

Overall, a simple parameterization procedure allows the exploration of a wide breadth of population genetic scenarios.

### 2.3 Branching process approximation

By analyzing the stability of system 7, one can only examine conditions necessary for the invading allele to escape extinction. However, we also want to know the probability of the escape. We pursue this by representing the change in the number of invading allele as a multi-type branching process. Selection is thus assumed to be stronger than drift (i.e., 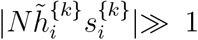). We work with two types, each tracking the number of mutant alleles in one of the two patches. The number of the mutant alleles is represented by the vector 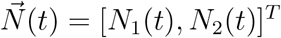, where *N*_1_(*t*) and *N*_2_(*t*) are the number of mutants in patch 1 and 2 in generation t, respectively. Let *f_i_* be the probability generating function (PGF) of offspring number of mutant alleles of type *i*. The ultimate probability of extinction of type *i* (1 − *P_i_*) is given by the smallest positive root of the following system of equations (Haccou *et al.* (2005)):

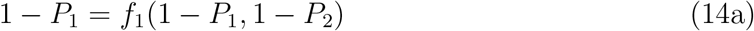

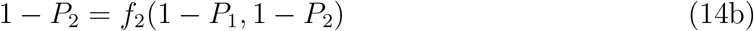

As we can assume that the number of offspring that types *i* and *j* leave are uncorrelated, then *f_k_*(1 − *P*_1_, 1 − *P*_2_) = *f*_*k*,1_(1 − *P*_1_)*f*_*k*,2_(1 − *P*_2_), with *k* = {1, 2}. Intuitively, the probability that mutant goes extinct conditioning on appearance in patch 1 is the probability that all of its offspring left in patch 1 ultimately go extinct (*f*_1,1_(1 − *P*_1_)), and the probability that all offspring left in patch 2 via migration (*f*_1,2_(1 − *P*_2_)) are ultimately lost. A similar explanation holds when mutant originates in patch 2. To solve the system, one needs to obtain the *f_i,j_*(*z*) PGFs.

In the Wright-Fisher model with random mating, the number of mutant offspring is approximately Poisson-distributed with mean 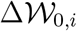, where 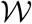 stands for the different form of selection as described above. Partial self-fertilization affects the mutant offspring distribution in two ways. First, it increases the mean number of offspring by inflating the effective dominance coefficient (see above). Second, selfing increases the variance in offspring number by the factor 1 + *F* relative to the randomly mating population (Caballero and Hill, 1992, Pollak, 1987). Therefore, offspring number should follow a distribution with mean 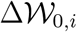 and variance 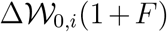. To our knowledge, the full distribution has never been obtained, even in single populations, probably because only the two first moments are needed for branching process and diffusion approximations. In appendix, we derive the exact probability generating function (PGF) of the distribution in different cases. In a single population we have:

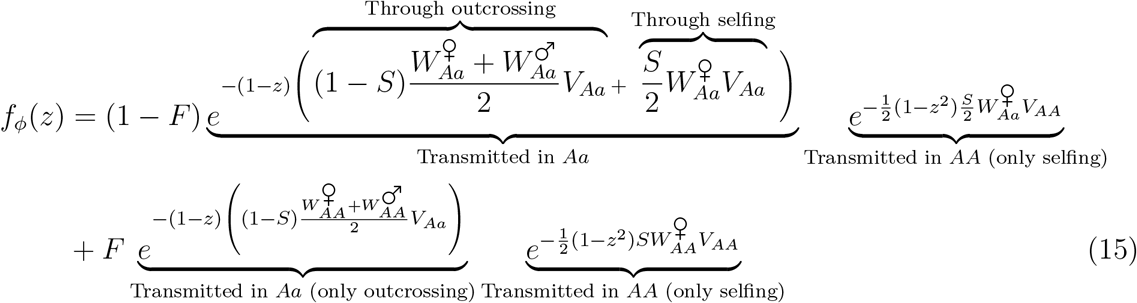

The first term in equation (15) corresponds to a *A* parental allele coming from an *Aa* individual (probability 1 − *F*) and the second term to an *AA* individual (probability *F*). Offspring can be transmitted either in *AA* offspring, which occurs only through selfing (because we consider a rare mutant) or in *Aa* offspring, which can occur either through outcrossing or through selfing when the parent is *Aa* (so with a probability *S*/2). Note that fecundity selection depends on parent genotypes whereas viability selection depends on offspring genotype. Using the properties of PGFs we easily retrieve that the mean of the distribution is 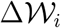, obtained in the deterministic analysis and the variance is 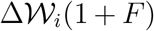. For *F* = 0, equation (15) boils down to the PGF of a Poisson distribution. Hereafter, as for the deterministic analysis, we consider the different forms of selection separately by setting constant the other fitness components.

For the two-patch model we need to include migration in the *f_i,j_*(*z*) PGFs. For seed migration, we show that:

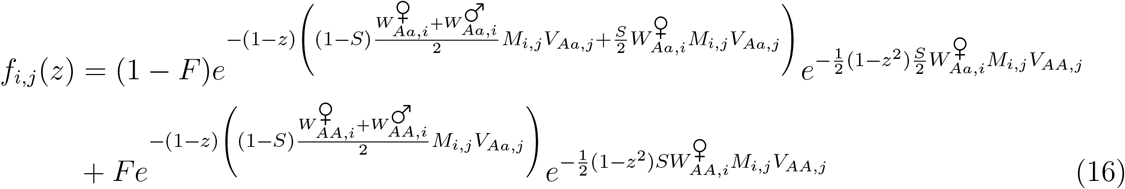

recalling that *M_i,i_* = 1 − *M_i,j_* with *j* ≠ *i*. Note that fecundity selection occurs in patch *i*, before seed migration, whereas viability selection occurs in patch *j*, after seed migration. However, the different orderings of migration and selection yield the same results (see *Mathematica* notebook). We also retrieve that the mean of the four distributions are given by the Jacobian terms obtained above, 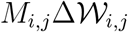 and the variance is also inflated by 1 + *F*. For pollen migration the form is slightly different because migrants can only contribute to the next generation through outcrossing and male gametes (whereas selfed seeds can migrate). The form of the PGF is thus different for the resident and migrant contribution. We have:

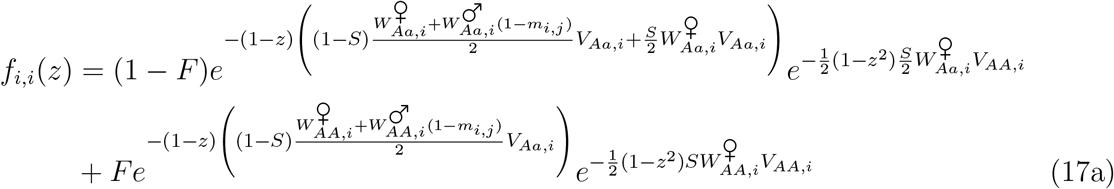

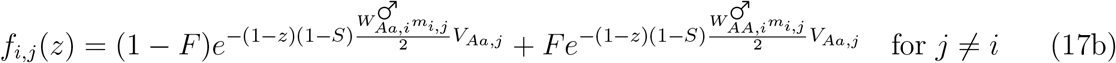

As expected, the mean of the distributions are still given by the Jacobian terms with the appropriate effective migration rates, 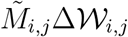 (see above). However, the variances are not inflated by 1 + *F*. For migrant alleles, equation (17b) boils down to a Poisson distribution because the contribution is only through outcrossing. In contrast for resident allele, the variance is inflated by more than 1 + *F* (see appendix for details). The reason is that for a given selfing rate, only alleles produced by outcrossing can be exported to the other patch, which increase the proportion of resident offspring contributed by selfing, hence the variance. Note, however, that the difference in PGFs between the two migration modes only affects high-order terms of selection and migration so has no effect on branching process approximation results, which can all be expressed with the same form using appropriate effective parameters as defined in previous section (see *Mathematica* notebook).

Previous section also showed that 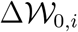 always has the form of 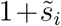, where 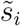 is the effective advantage of an invading mutant in patch *i*, while *M_ij_* represents the effective migration rate between patches *i* and *j*. Thus, one can derive *P_i_* for this generic case and then parameterize migration and selection to reflect the specific population genetic case. Since we cannot obtain the exact solution for extinction probabilities, we approximate to weak selection using the approach of Tomasini and Peischl (2018), which is based on the approximation for slightly supercritical multitype branching processes (Haccou *et al*., 2005). All model parameters are rescaled by 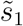 such that 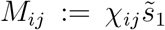 and 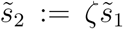. By linearizing around 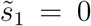 and neglecting higher-order terms, we obtain a closed-form solution for the probability of establishment conditioning on the patch in which locally advantageous allele appeared. All analytical solutions are compared to simulated data using the method described in Appendix D. Simulations were conducted with the help of GNU Parallel (Tange, 2011).

### 2.4 Diffusion process approximation

An alternative way of incorporating stochastic transmission of alleles across generations is by approximating frequency change to diffusion. This approach has been originally developed by Sakamoto and Innan (2019) in the case of wholly outcrossing population. Here, we recapitulate their derivation and extend it to the case of partial selfing. The establishment probability satisfies the Kolmogorov backward equation (KBE):

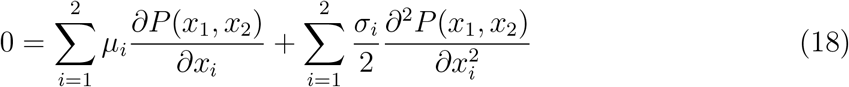

where *μ_i_* and *σ_i_* are drift and diffusion coefficients for patch *i*, and are given by:

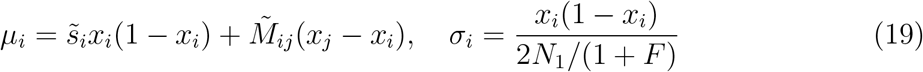

Note that 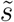 corresponds to effective selective advantage of mutant, and 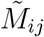 is the effective migration rate; These were derived in section 2.2. Diffusion coefficient is inflated by the factor of 1 + *F* as described previously. Unfortunately, we could not find neither the explicit nor approximate solution to this partial differential equation. Following treatment in Barton (1987), the problem is simplified by focusing on the first phase of fixation when mutant allele is rare. Thus, one can approximate coefficients as:

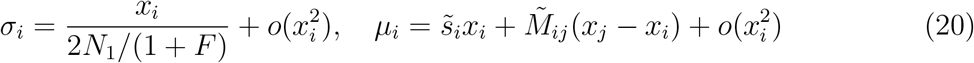

which leads to the simplified KBE:

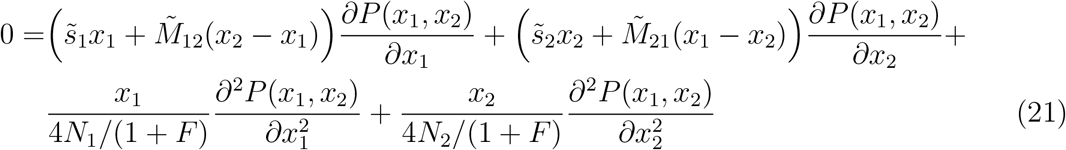

Because the invading allele is rare, the mutants in patch *i* go extinct independently of one another with probability *e^−l_i_^*, which means that the establishment probability takes the form:

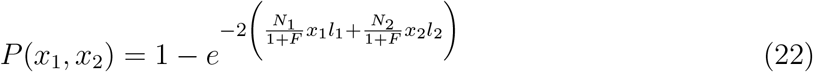

Substituting eqn. 22 into 21 results in an algebraic equation which captures the establishment probability for an arbitrary number of initial mutants across two patches. We are interested in two special cases, when mutant either starts in favored (*x*_1_ = 1/(2*N*_1_) and *x*_2_ = 0) or disfavored patch (*x*_1_ = 0 and *x*_2_ = 1/(2*N*_2_)). With this parameterization, we obtain the system of two non-linear equations:

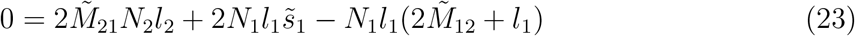

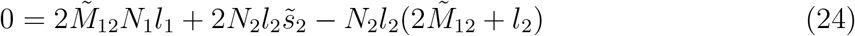

By solving these for *l*_1_ and *l*_2_, taking the real root, and substituting into eqn. 22, one retrieves a closed-form solution for the establishment probability (see *Mathematica* notebook). With all of the simplifying assumptions above, the diffusion approximation works in the same regime when *N_e_s* ≫ 1, as is the case with branching approximation. As will be shown later in the text, the comparison of two solutions reveals that diffusion approximation performs identically when the allele starts in the favored patch and slightly better when it originates in the disfavored one. Because branching process approximation is easier to work with (e.g., series expansion readily yields intuitive special cases), we use this solution throughout our analysis and note that the diffusion approach yields marginally better precision.

## 3 Results

### 3.1 Probability of establishment of a mutant

#### 3.1.1 General equations

The probability of establishment of a new allele is non-zero if the branching process is super-critical, which means that the leading eigenvalue of the mean reproductive matrix must be larger than unity (Haccou *et al*., 2005). For the invasion of allele *A* under viability selection, it corresponds to:

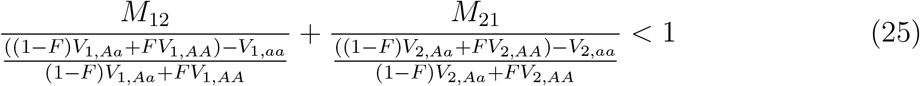

The denominator is the relative reproductive advantage of invading mutant against the resident allele. The invading population is a combination of mutant heterozygotes (when the two gene copies of an individual are not identical by descent, 1 − *F* fraction of time) and homozygotes (when the two gene copies are identical by descent, *F* fraction of time). So the whole term ((1 − *F*)*V_i,Aa_* + *FV_i,AA_*) gives the fitness of the invading mutant weighted by the contribution of invading genotypes. In an outcrossing population (*F* = 0), condition (25) reduces to *M*_12_/(1 − *V*_1,*aa*_/*V*_1,*Aa*_) + *M*_21_/(1 − *V*_2,*aa*_/*V*_2,*Aa*_) < 1, which is identical to Bulmer’s inequality. The same inequality with re-parameterized selection holds for fecundity and male sexual selection (see Appendix C). The full equations for the establishment probabilities are too long and not very informative and are thus reported in the appendix (see equations (A16) and (A17)). Without loss of generality, consider that allele *A* is advantageous in patch 1 but deleterious in patch 2 (*s*_1_ > 0 > *s*_2_); in the case of symmetrical migration (*M*_12_ = *M*_21_ = *M*/2) and migration prior to selection the general probability of establishment of allele *A* in either patch 1 or 2 is given by:

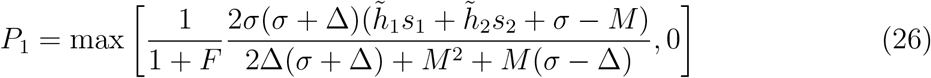

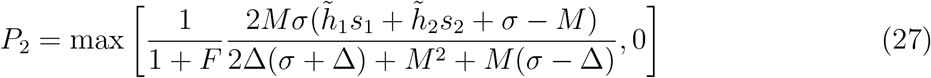

where 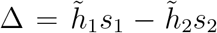 and 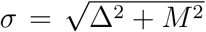 with 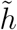 as defined above for viability, female and male fecundity selection. These resemble – but are not equal – to equations (4) and (5) in (Tomasini and Peischl, 2018), who obtained an inexact result because of a typo in their application of Haccou et al.’s theorem. Initially the typo came from Aeschbacher and Bürger (2014) and was independently detected by Pontz and Bürger (2021) (see their Appendix A). Interestingly, the correct solutions are more complicated and less elegant than the ones of Tomasini and Peischl (2018).

#### 3.1.2 Limiting cases and comparison with previous results

Despite the complexity of the general equations, useful insights can be obtained from simple limit conditions. In the limit of no migration (*M*_12_ = *M*_21_ = 0), we get back to the single-patch scenario where selection favors allele *A* in patch 1: 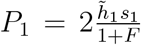. This is equivalent to Charlesworth (1992) and Caballero and Hill (1992) for viability selection. For male and female fecundity selection, the result is similar to the one intuited in Damgaard (2000), except that we also accounted for the reduction in *P* due to increased variance of offspring number. Conversely, in the limit of full migration we obtain 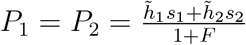, which is equivalent to Nagylaki (1980), with the appropriate rescaling for dominance and partial selfing. The probability of establishment is simply the average over the two patches. Interestingly, the fact that the effects of selfing on viability selection cancel out for codominant alleles (*h*_1_ = *h*_2_ = 1/2) is no longer true with intermediate migration, even for seed migration that is not affected by selfing. In the limit of weak symmetrical migration, we obtain 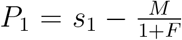 and 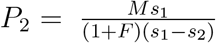 and in the limit of strong migration (i.e. *M* >> *s_i_*): 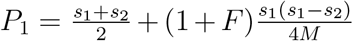 and 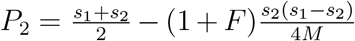. Thus, unlike in a single population a locally advantageous allele that is codominant in both patches is more likely to escape extinction in selfers than in outcrossers. The relation between *F* and *P*_2_ is more complicated, and the allele can either be more likely, for high migration rates, or less likely, for low migration rates, to escape extinction. The effects of selfing under codominance only vanish either for no migration or full migration.

#### 3.1.3 Comparison with simulations

In general, our approximation for viability shows good correspondence with simulations across different selfing rates (Figure 1). This also holds true across different dominance coefficients, and when selection or migration are asymmetrical (Figure 2). Correspondence is better in cases where the invading allele initially appears in the favored rather than in the disfavored patch. Diffusion approximation gives identical results when allele starts in the favored, and slightly better results when in the case of the disfavored patch. The improvement only occurs when the migration rate is low, and this pattern persists across dominances and modes of selection. A possible reason for the slightly worst accuracy of the branching process approach in the disadvantageous patch with low migration is that the rescaling of the model assumes that selection and migration parameters are of the order of 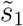. For low migration, we can have 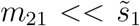, so that migration could not be sufficient to introduce the allele in the favored patch before it goes extinct in the disfavored one.

**Figure 1:**
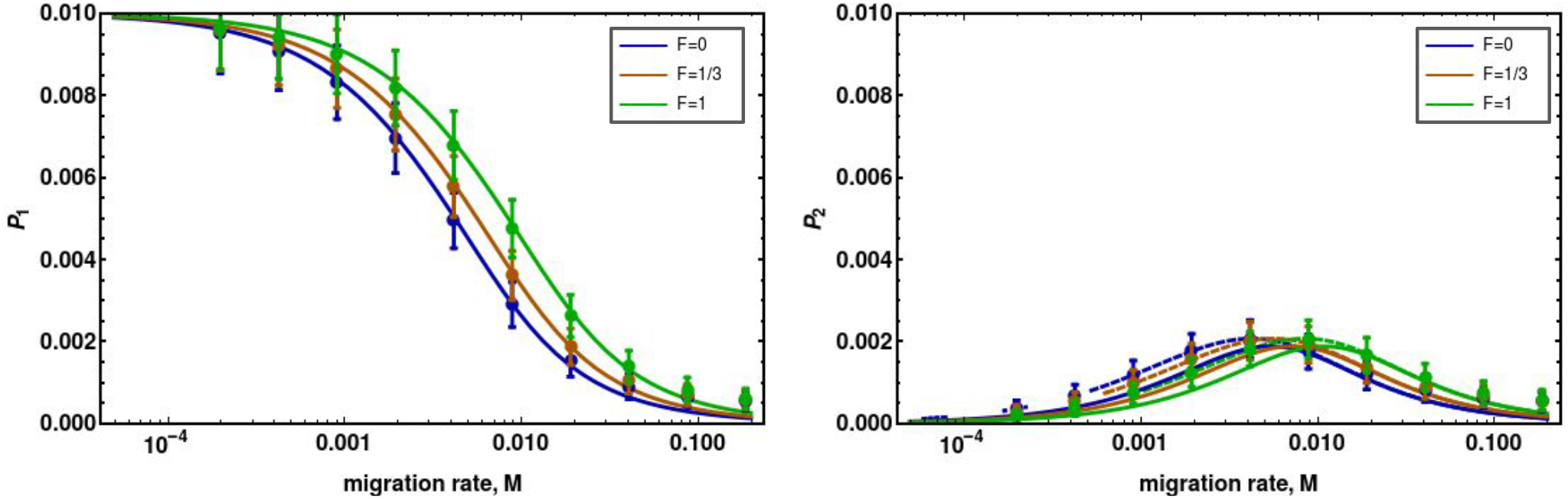
Comparison of analytical solution to simulated data across different selfing rates (corresponding to different equilibrium *F*). Dashed lines are diffusion approximation. Left panel: the probability of establishment when the allele emerges in the favored patch; Right panel: the probability of establishment when the allele emerges in the disfavored patch. Parameters: 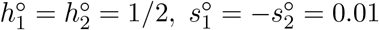, *M*_12_ = *M*_21_ = *M*.

**Figure 2:**
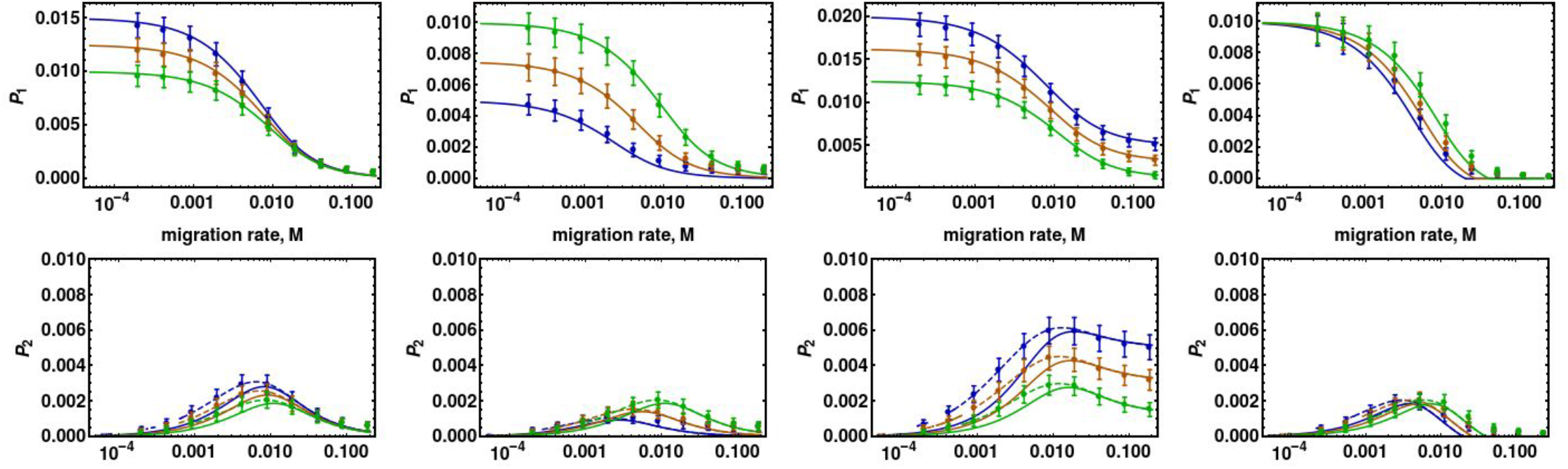
Simulations of different selection regimes. Going from left to right, the examined scenarios are as follows. Dashed lines denote diffusion approximation. First column: dominant case (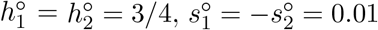, *M*_12_ = *M*_21_ = *M*); Second column: recessive case (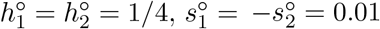, *M*_12_ = *M*_21_ = *M*); Third column: Asymmetric selection case (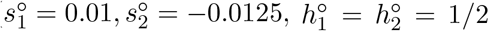, *M*_12_ = *M*_21_ = *M*); Fourth column: asymmetric migration (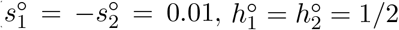, *M*_12_ = *M*, and *M*_21_ = 1.25*M*).

*P*_1_ is a monotonically decreasing function in *M*, because strong migration causes the spreading of the locally advantageous allele to become swamped by the resident from the disfavored patch. *P*_2_ is non-monotonic function in *M*. Increasing migration initially increases the probability that an invading allele escapes from the disfavored patch before it goes extinct. However, at large migration rates, the spreading allele escapes the disfavored patch but is swamped due to a large influx of deleterious residents from the disfavored patch. For male fecundity selection, the analytical solutions for *P*_1_ and *P*_2_ are slightly worse against the simulations (Figure A11) than those provided for viability selection. Note, however that as the selfing rate tends towards 1, the effective selection coefficient on male function vanishes so that the strong selection condition for branching process approximation is not met anymore. Slight discrepancies are also observed when selection or migration are asymmetrical (Figures A12, and A13).

We also varied *F* continuously to obtain a fine-grained view of the analytic performance. Three insights emerge. First, there is a good agreement between simulated data and analytical solution, although deviations increase as the migration rate increases. Second, the branching process poorly captures the dynamics of the invasion when mutants are partially recessive. For example, in the limit of complete recesivity (*h* = 0), the mutant does not have any reproductive advantage (i.e., equations 10a–10c reduce to unity). We clearly see this in Figure 3, where the analytical solution underestimates *P* when the allele is partially recessive (first column). However, this discrepancy disappears as *F* increases, as homozygotes increasingly contribute to the invading process. Third, diffusion approximation yields a better fit when mutant starts in the favored patch, but this improvement generally works only when *M* is low.

**Figure 3:**
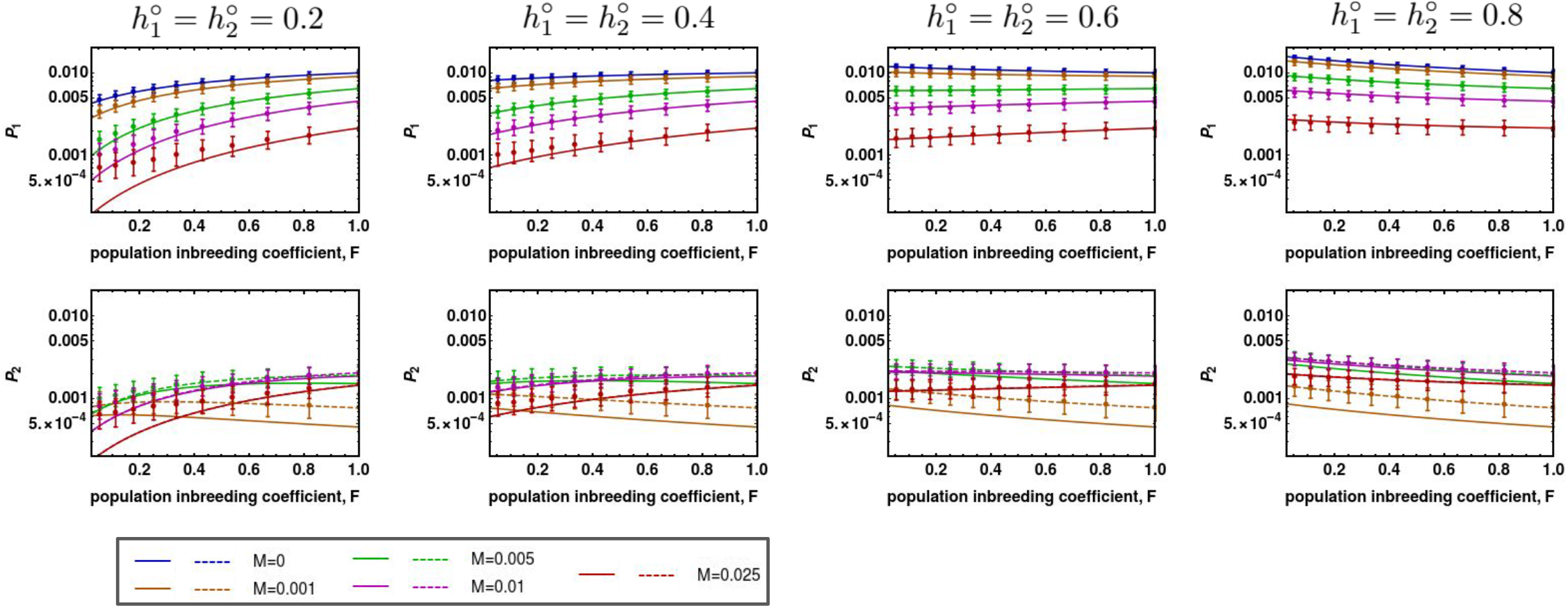
Simulations of different selection regimes. Going from left to right, the examined scenarios are as follows. Dashed lines denote diffusion approximation. Each column denotes the establishment probability for a set of dominance coefficients noted above the panel. Lines correspond to different migration rates (see legend box below). Other parameters: 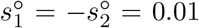, *M*_12_ = *M*_21_ = *M*). The axes are on logarithmic scale and error bars are excluded.

### 3.2 Decomposing the effects of selfing on the establishment probability

Selfing impedes the spread of an invading beneficial allele by reducing the effective population size and by increasing the effective dominance in the disfavored patch. On the other hand, selfing also facilitates the invasion process by increasing the effective dominance in the favored patch and reducing the gene flow when dispersal occurs through pollen. In this section, we ask: are the impeding effects generally more important than facilitating effects? When does the introduction of selfing increase *P*? To this end, we introduce an indicator *β_i_*(*y*) which denotes the effect that selfing has on *P_i_*, solely via parametery *y*. For example, 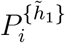 corresponds to the establishment probability in patch *i*, if selfing only exerted its effect by inflating favored dominance. More generally:

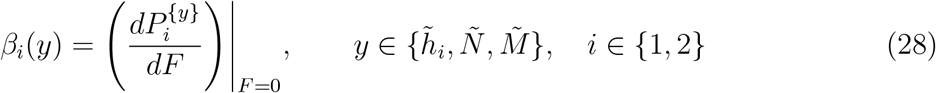

Consider decomposing the effects of selfing on the establishment of an allele affecting viability. For example, the indicator 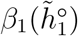 tells us whether a shift to selfing increases *P*_1_, if selfing solely acted through an increase in the effective dominance in the favored patch. This allows us to examine the effect of each factor separately and assess when one outweighs the other. The indicator is obtained by taking equation A16 and sequentially setting: (1) *F* = 0 which eliminates selfing’s effect on 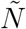, then (2) parameterizing 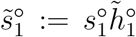 which introduces the effect of selfing on favored dominance, and finally, (3) setting 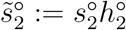 which excludes the effect of selfing on the effective dominance in the disfavored patch. The outlined method is also applicable to other selection and migration scenarios, and the general procedure is given in Appendix F.

#### 3.2.1 Emergence in the favored patch

We focus on the establishment conditioning on allele emerging in the favored patch. Consider viability selection first (see Figure 4). Given our interest in knowing whether selfing increases or decreases the establishment probability, we only focus on the region of parameter space where allele can become established in fully outcrossing population (right of the solid line in Figure 4). When the invading allele is partially recessive in favored and partially dominant in the disfavored patch (white region), the criterion for escaping extinction is not satisfied, and a shift to selfing can only have a promoting effect on the establishment in this region of parameter space.

**Figure 4:**
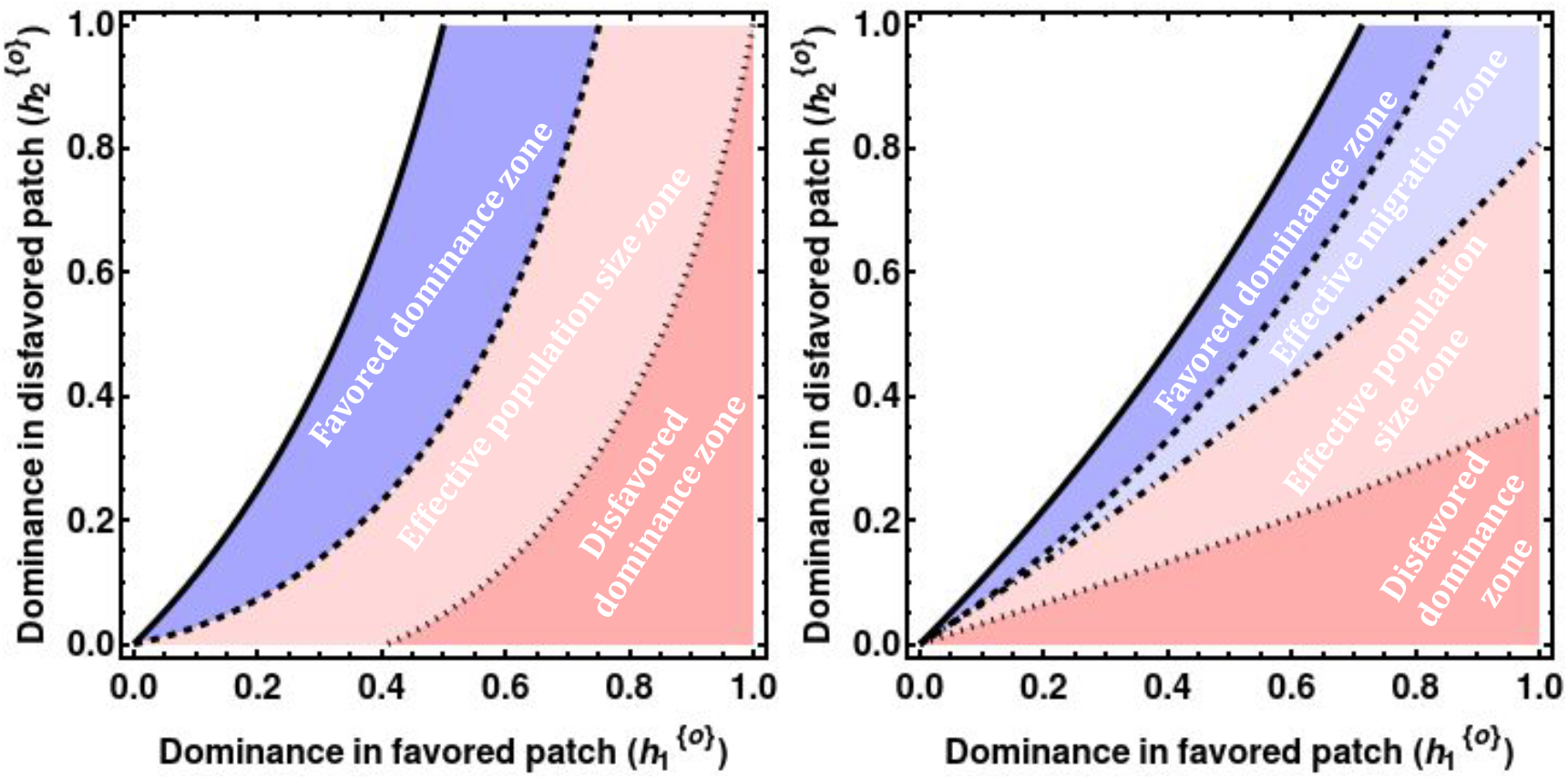
Four zones of selfing’s effects on the establishment probability when mutant originates in the favored patch. White region denotes parameter space where the mutant cannot establish under outcrossing. Selection act only on viability, and migration occurs only via pollen (left), or only via seed (right). Blue and red regions correspond to parameter space where selfing facilitates and impedes *P*_1_, respectively. Lines in the left panel are as follows. Solid black line: *P*_1_ = 0; Dashed black line: 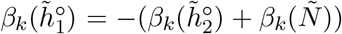; Dotted black line: 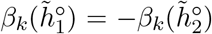. Lines in the right panel are as follows. Solid black line: *P*_1_ = 0; Dashed black line: 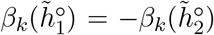; Dashdotted line: 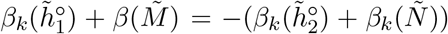; Dotted line: 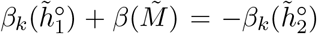. Parameters for the left panel: 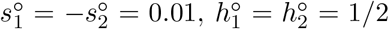, *M*_12_ = *M*_21_ = 0.01. Parameters for the right panel: 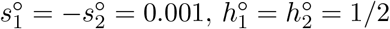, *M*_12_ = *M*_21_ = 0.005.

Under seed dispersal, selfing only affects selection via effective dominance and the effective population size. One can recognize three regions of selfing effects (Figure 4). Firstly, when the invading mutant is partially recessive in the favored and partially dominant in the disfavored patch, selfing increases *P*_1_ via the inflation of 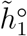 (blue region). More formally, this will be the outcome when 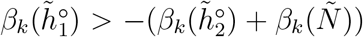 (dark blue region). This is because the mutant is already dominant in the disfavored patch so selfing does not significantly increase the rate of purging, but it does increase the rate of spread owing to the mutant being partially recessive in the favored patch. Thus, the net effect is increased *P*_1_. Secondly, when the mutant is partially dominant in the favored and partially recessive in the disfavored patch, selfing decreases *P*_1_ via inflation of 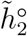 (darker red region). This happens when 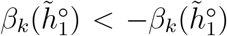 (dark red region). The effect is opposite to the first regime: selfing has a larger effect on the increase in purging from the disfavored patch (since the dominance in the disfavored patch is low) than on the increase in spreading in the favored patch (because dominance in the favored patch is already high). The net effect is a decrease in *P*_1_. The region of parameter space that lies between these two zones corresponds to cases when the facilitating effect of selfing (via dominance in the favored patch) roughly cancels out the impeding effect (via dominance in the disfavored patch) so that the net effect on *P*_1_ is determined by the reduction in effective population size and the net effect of selfing is to decrease *P*_1_.

Under pollen dispersal, we also found the same three regions but with different boundaries (Figure 4). In addition, selfing also affects pollen dispersal by reducing the effective number of migrants, so in the intermediate zone the effect of selfing on *P*_1_ depends both on the reduction in effective population size and effective migration rate, which creates a fourth region – the region for which all parameter combinations where 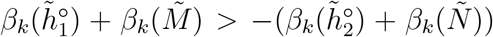 (light blue region). It corresponds to conditions where the reduction in effective migration rate overwhelms the impeding effects of selfing. Overall, selfing increases the establishment of new alleles under broader conditions under pollen than under seed dispersal. Note that selection coefficients are an order of magnitude lower in the right panel of Figure 4, and the region where selfing has a net-positive effect is comparable (if not larger) to the blue region in the left panel.

Although qualitatively similar results holds for two other modes of selection (Figure 5), one notices that the region where the selfing exerts its effect solely via effective dominance is significantly expanded. This is because an increase in the selfing rate diminishes the male component’s relevance and increases the importance of the female component. The relative sizes of these regions vary based on parameter values, so more general quantitative statements are hard to make.

**Figure 5:**
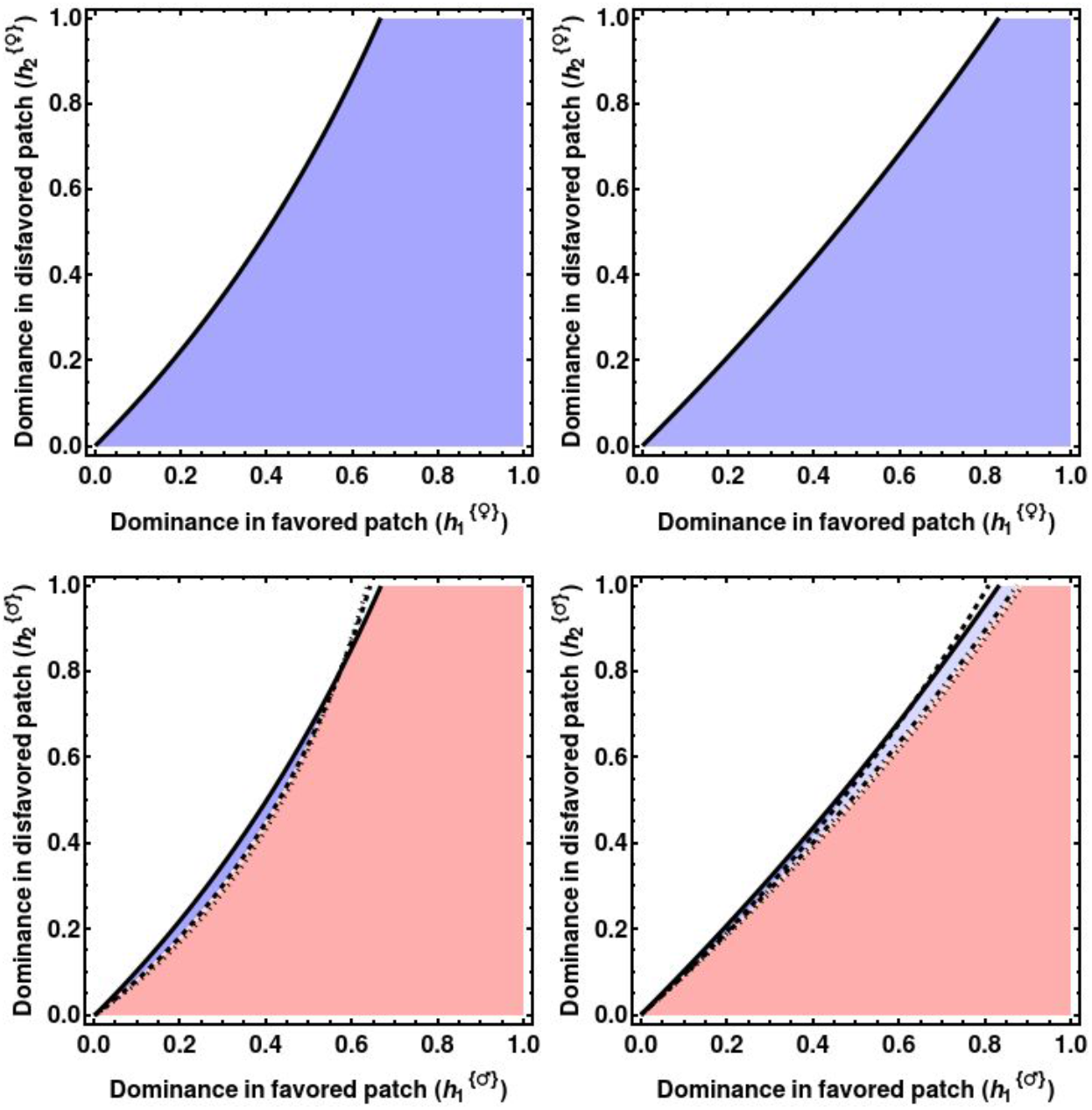
Zones of selfing’s effect on *P*_1_ under female fecundity (upper row), and on male fecundity (lower row). Color code as in Figure 4. Migration via seed is depicted in the left column, and via pollen in the right column. Black lines as in the previous figure. Depending on selection and migration scenario, parameters are either: 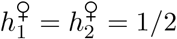 or 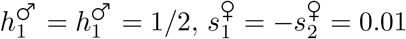 or 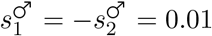, and *M*_12_ = *M*_21_ = 0.01 or *m*_12_ = *m*_21_ = 0.01.

#### 3.2.2 Emergence in the disfavored patch

Similar conclusions are reached if one considers emergence of mutant in the disfavored patch (Figure A16). However, under pollen dispersal, selfing reduces the effective migration rate, which can have either positive or negative effect on *P*_2_ depending on the parameters of the model. This phenomenon arises due to *P*_2_ being a non-monotonic function of *m* (see Figure 2). More specifically, if migration is much stronger than selection, then a mutant can escape to favored patch where it can spread, but the influx of maladapted allele is also high, thus having a net impeding effect on invading allele. This means that introduction of selfing will reduce the effective migration rate, and thus increase *P*_2_. Put differently, increasing *F* has the same kind of effect on *P*_2_ as decreasing *m*. However, if the migration rate is already low enough, the introduction of selfing further depresses *m* and, thus, prevents invader from successfully escaping into the favored patch.

### 3.3 Consequence of selfing on the establishment of protected poly-morphism

Once the allele *A* escapes extinction, it can either fix across both patches or segregate at intermediate frequencies for a finite but large number of generations due to divergent selection. Because both patches are of finite size, one of the two alleles will ultimately fix but this quasi-stationary behavior corresponds to protected polymorphism in deterministic models. We wish to delineate the effects of selfing on these two outcomes. Given that an invading allele can appear in any of the two patches where the favored patch accounts for *z* fraction of the total population across both patches, the global probability of alleles *A* and a becoming established is:

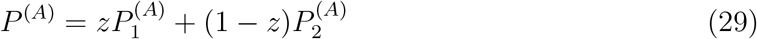

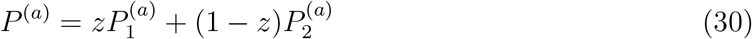

where 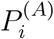 and 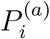 are probabilities that *A* and *a* become established, conditioning on initially appearing in patch *i*. For the sake of simplicity, we will further assume that patches are of equal size (*z* = 1/2), and migration is symmetrical. Probability that allele *A* is established is computed by parameterizing equations (26) and (27) where selection coefficients were determined from Jacobian associated with boundary 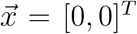, while the establishment probability of *a* was calculated with elements of Jacobian associated with boundary 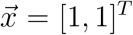. For example, if selective advantage of invading heterozygote *Aa* relative to homozygote *aa* in patch *i* is *s_i_h_i_*, then the advantage of invading *Aa* relative to homozygote *AA* is − *s_i_*(1 − *h_i_*)/(1 + *s_i_*).

#### 3.3.1 Conditions for protected polymorphism and critical migration rates

Polymorphism is protected if both *P*^(*A*)^ > 0 and *P*^(*a*)^ > 0, that is for the conditions under which the branching processes for the invasion of *A* and for the invasion of *a* are supercritical. This boils down to the extension of Bulmer’s inequality given by equation (25) directly obtained from the Jacobian matrix, which must also be satisfied with genotypes *AA* and *aa* shifted (corresponding to the supercriticality for the invasion of allele *a*). From this, we can define critical migration rates above which polymorphism is lost and the allele with the highest marginal fitness invades as in Yeaman and Otto (2011). Without loss of generality we can assume that *s*_1_ > −*s*_2_ > 0. For seed dispersal and viability selection *M_c_* is then given by:

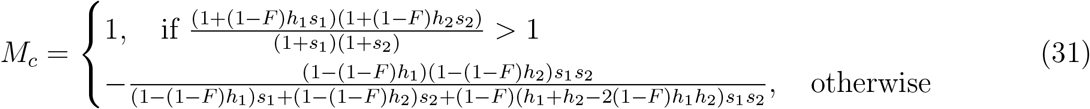

For *F* = 0, (31) reduces to equation (5) in Yeaman and Otto (2011) and for *F* = 1 to 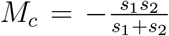. The first condition corresponds to a form of dominance reversal where the allele is dominant when advantageous and recessive when deleterious. Under such a condition, polymorphism can be maintained even under full migration.

Depending on dominance, selfing thus has two opposite effects. On the one hand, by unmasking recessive alleles, selfing makes selection stronger relative to migration and so enlarges the conditions for the maintenance of protected polymorphism. On the other hand, selfing makes dominance reversal conditions less likely as selection mainly operates on homozygotes, and so restricts the conditions for protected polymorphism. Figure 6 shows the dominance conditions under which selfing increases or decreases the critical migration rate. We determined these conditions in two ways: either when *M_c_* is higher under full selfing than under full outcrossing (i.e., *M_c_*|_*F*=0_< *M_c_*|_*F*=1_) or whether introduction of selfing in an otherwise outcrossing population increases *M_c_*, that is 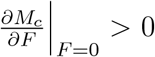. For symmetrical selection (*s*_2_ = −*s*_1_), selfing increases migration rate if *h*_2_ > *h*_1_. Under the opposite condition, the critical migration is 1 and local adaptation is always maintained under outcrossing. As selection becomes more asymmetrical, selfing favors local adaptation for a broader range of dominance coefficients. Note that the effect of selfing on the migration rate is not monotonic for a narrow range of conditions (light blue region in Figure 6), where little selfing disfavors local adaption 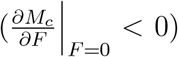 but high selfing increases it (*M_c_*|_*F*=0_< *M_c_*|_*F*=1_). Finally, it is worth noting that when selfing favors local adaptation, the effect is rather weak (blue curves on Figure 6). On the contrary, under dominance reversal conditions, above a given threshold, selfing dramatically reduces the migration rate from 1 to 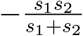 under full selfing. Similar results are obtained for fecundity selection but with selfing favoring local adaption under broader conditions for female fecundity and for narrower conditions for male fecundity (see Appendix).

**Figure 6:**
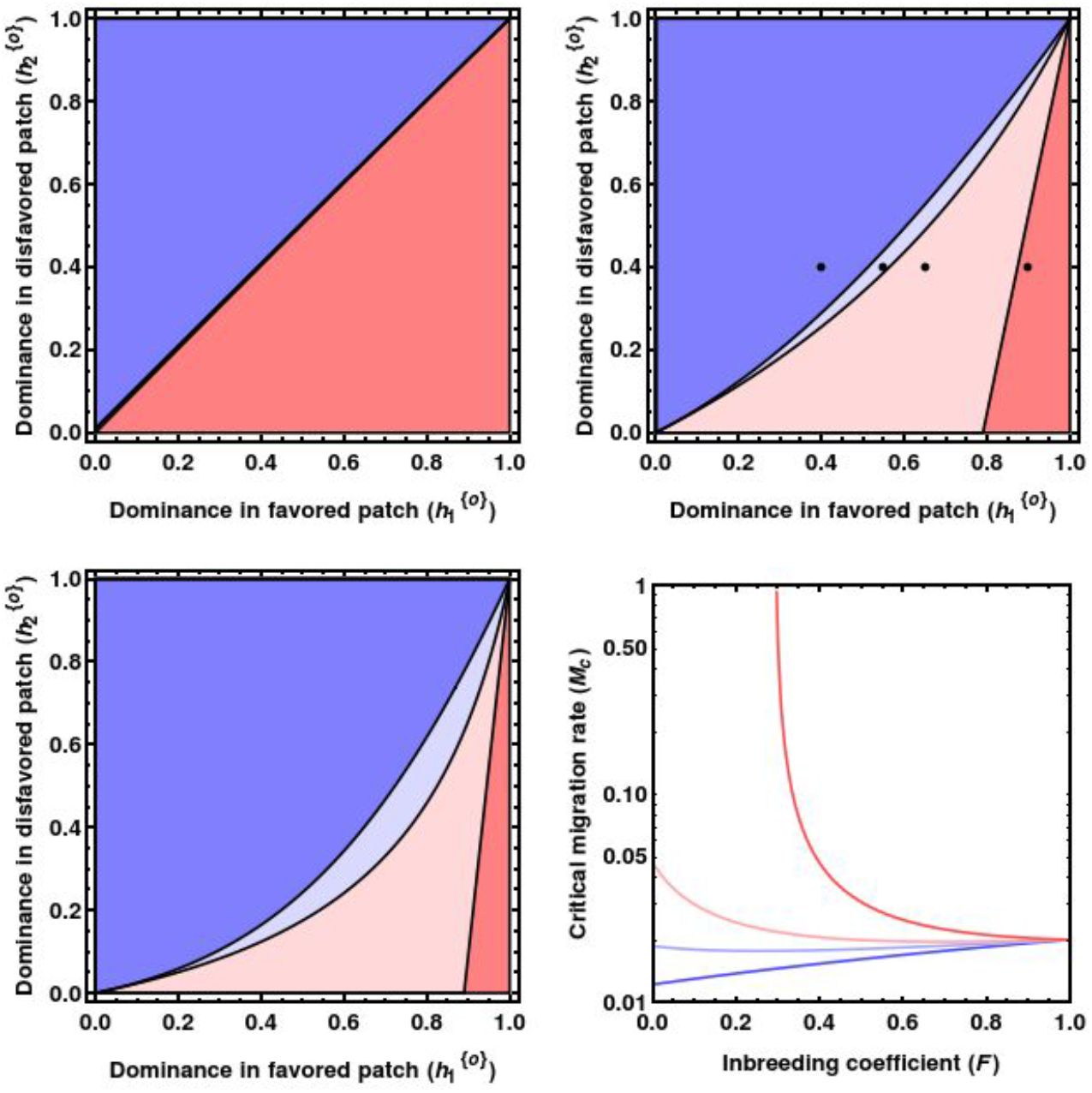
Effect of selfing on the critical migration rate. The three first panels show dominance conditions under which selfing increases (blue regions) or decreases (red regions) the critical migration rate, *M_c_*. Selection is symmetrical in the first panel (*s*_1_ = −*s*_2_ = 0.01), and asymmetrical in the second (*s*_1_ = 0.02 and *s*_2_ = −0.01) and third panels (*s*_1_ = 0.05 and *s*_2_ = −0.01). In the light blue region, *M_c_* is not monotonic with *F*: introduction of selfing in an outcrossing population decreases 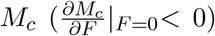 but above a given selfing rate, *M_c_* becomes higher than in an outcrossing population and we always have *M_c_*|_*F*=0_< *M_c_*|_*F*=1_. In the dark blue region, selfing always increases *M_c_*(*M_c_*|_*F*=0_< *M_c_*|_*F*=1_). In the dark red region, *M_c_* = 1 in outcrossing populations and *M_c_* < 1 in the light red region. The last panel illustrates how *M_c_* varies with *F* in the four regions, corresponding to the black dots of the second panel. For all curves *s*_1_ = 0.02 and *s*_2_ = −0.01. Dark blue: *h*_1_ = 0.4 and *h*_2_ = 0.4; light blue: *h*_1_ = 0.55 and *h*_2_ = 0.4; light red: *h*_1_ = 0.65 and *h*_2_ = 0.4; Dark red: *h*_1_ = 0.9 and *h*_2_ = 0.4. Note that *M_c_* is in log-scale.

The picture is different under pollen dispersal. Under full selfing, effective migration is zero so selection can act independently in each patch and polymorphism is always maintained. As a consequence, there is always a threshold selfing rate above which the critical migration rate is higher than in an outcrossing population. Here, we thus concentrate on the effect of the introduction of selfing in an outcrossing population, so the conditions for which 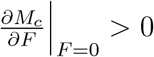. Under symmetrical selection, the conditions are the same as for seed migration. However, for asymmetrical selection, selfing increases *M_c_* for a much broader range of parameters.

#### 3.3.2 Establishment and maintenance of protected polymorphism

The above analysis showed how selfing affects the critical migration rate, hence the conditions for local adaptation. However, when local adaptation is possible, selfing also affects the probability of establishment and maintenance of polymorphism. This can be analyzed by quantifying how selfing affects *P*^(*A*)^ and *P*^(*a*)^, which is given by:

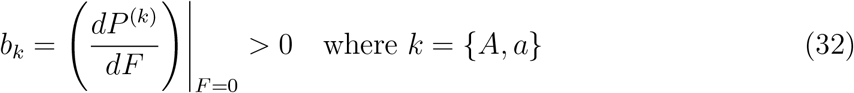

Four outcomes are possible upon the introduction of selfing. First, *P*^(*A*)^ increases, and *P*^(*a*)^ decreases, meaning that allele *A* is more likely to fix across both patches in selfers than in outcrossers. Second and conversely, *P*^(*A*)^ decreases, but *P*^(*a*)^ increases, implying that selfers are less likely to fix *A* compared to an outcrossing population. Third, both *P*^(*A*)^ and *P*^(*a*)^ increase upon shift to selfing, meaning that selfing increases the probability that protected polymorphism is established. Fourth and final, both *P_A_* and *P_a_* decrease, making the protected polymorphism less likely to become established in the selfing population. Some examples are given for different migration rates with symmetrical or asymmetrical selection (Figure 8).

One can intuitively understand these outcomes by noting that dominance reverses when the alternative allele invades. So, if invader *A* is dominant, then invader *a* will be recessive. The dominant allele *A* is likely to escape extinction when invading but unlikely to result in protected polymorphism because allele *a* is recessive and thus likely to go extinct. This appears clearly in the limit of low migration, where probabilities of emergence tend toward single population predictions, with four quadrants corresponding to the four dominance conditions (Figure 8). Importantly, these four regions vary with the amount of migration, and they do not directly align with the conditions given by the critical migration rates. The reason is that the sign of *P*^(*A*)^ and *P*^(*a*)^ only depends on the effect of selfing through the effective dominance, whereas the absolute value also depends on the reduction in effective size. Here, we only consider the two-fold reduction in *N_e_* caused by selfing, but the linked selection can further reduce the effective size of the local population under selfing (Roze, 2016). This will not affect the critical migration rate but can strongly reduce probabilities of emergence hence the establishment of local adaptation. Following this rationale, Yeaman and Otto (2011) considered a critical migration rate above which the probability of emergence of both alleles is higher than 1/(4*N_e_*). High local drift under selfing can thus strongly reduce the conditions for the establishment of local adaptation.

## 4 Discussion

We investigated the effect of selfing on the establishment of an allele in a population subdivided into two patches, where only one locus determines the fitness of an organism. By representing the spread of an invading allele as a branching process, we obtained a closed-form analytical solution for the probability of establishment of the locally advantageous allele for an arbitrary population selfing rate. This extends the work of Tomasini and Peischl, 2018 to include diploidy, dominance, and partial selfing. We also corrected for a typo in the derivation of their equations but which surprisingly lead to less elegant results. Below, we discuss the implication for adaptation in partially selfing species.

A well-known result is that selfing favors the establishment (and fixation) of recessive alleles but disfavors the establishment of dominant ones, with no effect under exact codominance (*h* = 1/2) (Caballero and Hill, 1992, Charlesworth, 1992). The pattern is more complex in a subdivided population with heterogeneous selection. First, the codominant allele has a higher probability of becoming established in the selfing population than in the outcrossing population if it emerges in the favored patch and generally smaller probability if it emerges in the disfavored patch. Therefore, the probability of establishment of a codominant allele is not independent of the mating system, as was the case for a single partly selfing population (Charlesworth (1992)). Second, assuming that migration and selection are symmetrical, selfing will increase the establishment probability when the invading mutant is recessive in the favored patch (resulting in a maximal increase in 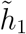) and dominant in the disfavored patch (yielding a minimal increase in 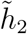). If the two dominances are roughly equal, the opposed effects of selfing on effective dominances cancel each other out, and the net effect is set by the interplay between the positive effect of reduced effective migration and the negative effect of the reduction in *N_e_*.

Once the invading allele escapes extinction, it can either fix in both populations, thus one population is fixed for a maladaptive allele, or segregate at intermediate frequencies, thus contributing to protected polymorphism, and both populations can be considered as locally adapted. It is usually thought that selfers are better locally adapted because of the reduced gene flow between patches with different selection demands (Linhart and Grant (1996)). However, selfing affects in a complex way three key parameters determining local adaptation - selection, drift, and gene flow. By separately considering seed and pollen migration, we were able to distinguish the straightforward effect of reducing gene flow from the more subtle effects of altering genotypic frequencies and drift induced by selfing.

Under seed dispersal only, selfing does not affect gene flow but still alters the propensity for local adaptation. The conditions under which selfing favors local adaptation strongly depend on the dominance of alleles in the two patches. Under outcrossing, the most favorable condition for local adaptation is under dominance reversal, that is, when the allele is dominant when locally beneficial and recessive when locally deleterious. If so, protected polymorphism can be maintained even under full migration (*M_c_* = 1), which corresponds to conditions for polymorphism in Levene (1953)’s model. Under this condition, selfing destabilizes polymorphism, which cannot be maintained under full dispersal: above a given selfing rate, the critical migration rate strongly decreases (Figure 6, and see also Glémin, 2021, for the Levene’s model with selfing). Conversely, under most other conditions, especially when selection is asymmetrical, selfing tends to increase the critical migration rate, hence favors local adaptation. What dominance patterns across heterogeneous habitat are the most frequent in natural populations is unknown. However, dominance reversal is maybe not as unlikely as it may seem because it can naturally arise in simple fitness landscape models (Connallon and Chenoweth, 2019). If we now consider pollen dispersal, selfing enhances local adaptation by reducing gene flow. Under dominance reversal conditions, this leads to non-monotonic behaviors where local adaptation can be the most easily maintained, either under outcrossing or under high selfing (Figure 7). Characterizing the pollination and seed dispersal modes and their quantitative impacts on gene flow appears thus crucial to make proper predictions on the effect of selfing. Finally, if we also consider the effect of linked selection that reduces local effective size beyond the two-fold level in selfing populations (Roze, 2016), selfing reduces the possibility of local adaptation, which requires stronger selection as exemplified by Hodgins and Yeaman (2019) who simulated local adaptation in selfing populations with and without background selection. Overall, the complex and contradictory effects of selfing on local adaptation may explain why no general empirical pattern has emerged so far (Hereford, 2010, Leimu and Fischer, 2008).

**Figure 7:**
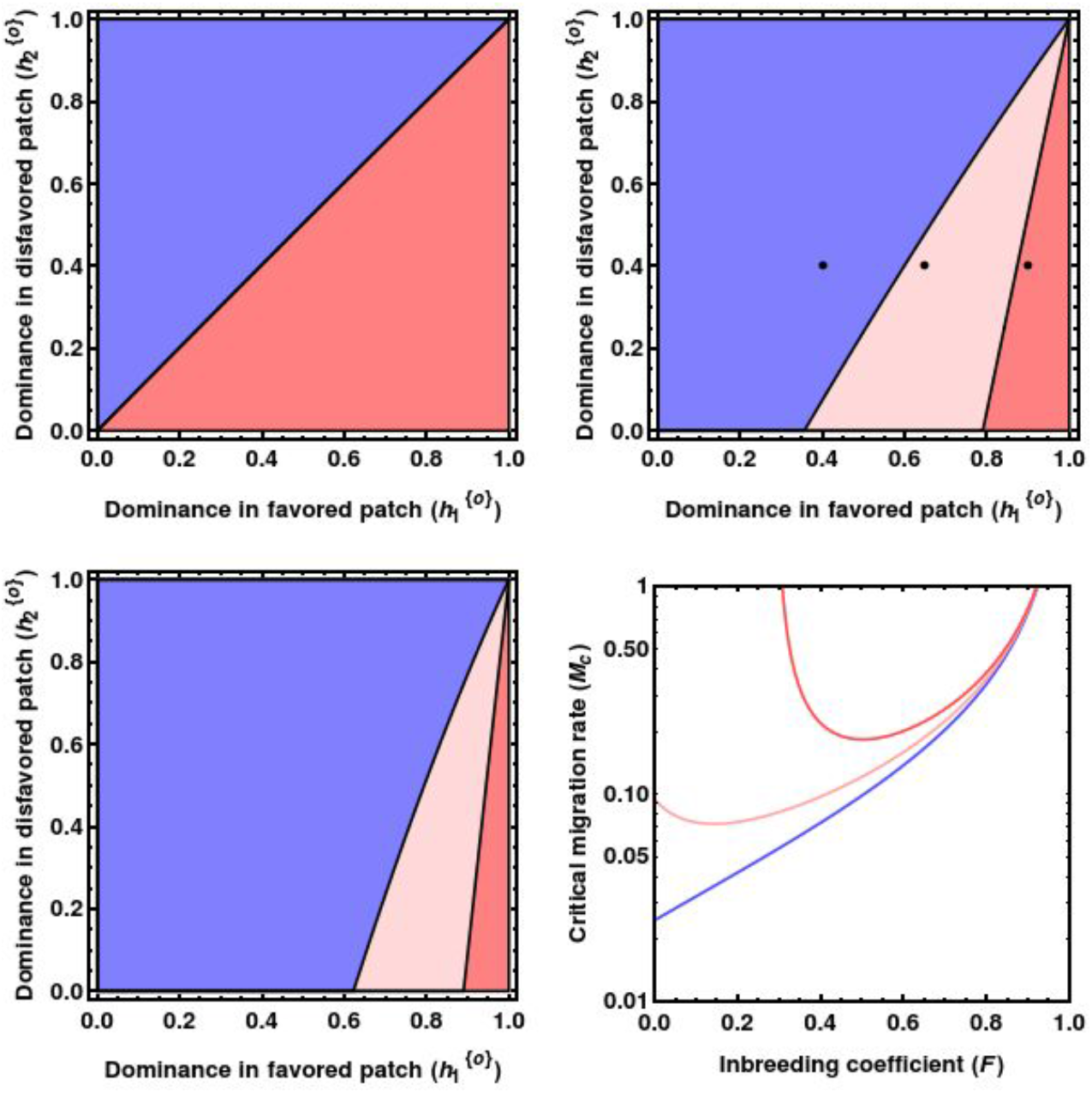
Same legend and same parameters as in Figure 6 but with pollen dispersal instead of seed dispersal.

**Figure 8:**
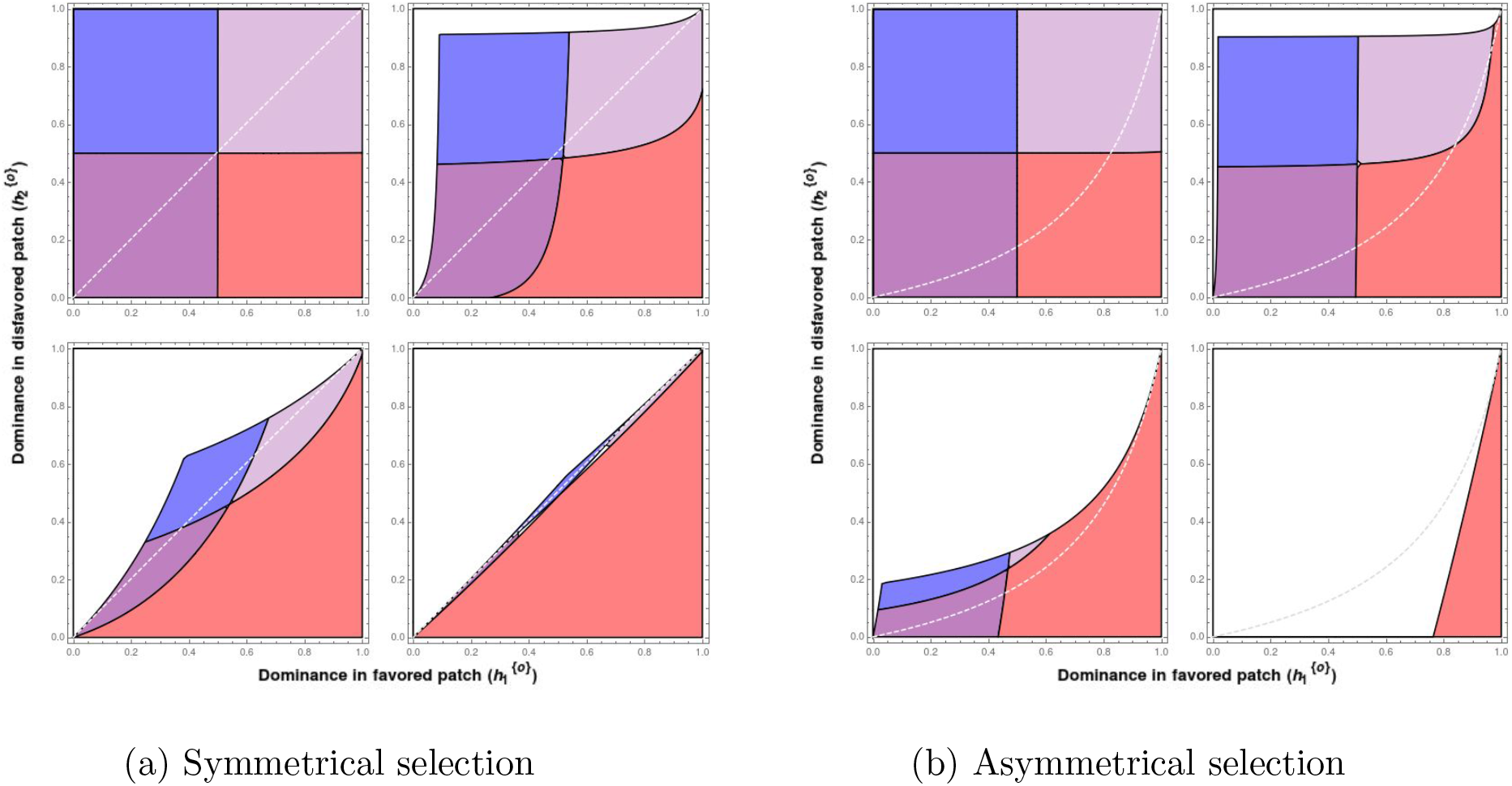
Effect of selfing on the establishment of protected polymorphism under symmetrical (a) and asymmetrical selection (b). Five regions can be distinguished: blue: *b_A_* > 0 and *b_a_* > 0, red: *b_A_* < 0 and *b_a_* < 0, dark purple *b_A_* > 0 and *b_a_* < 0, light purple *b_A_* < 0 and *b_a_* > 0, and white: polymorphism cannot be maintained under outcrossing. The dotted lines correspond to the limiting conditions for which selfing increases the critical migration rate (above the line) as in Figure 6. (a) *s*_1_ = 0.01 and *s*_2_ = −0.01; *m* = 0.0001, 0.001, 0.01, 0.1 from top left to bottom right. (b) *s*_1_ = 0.05 and *s*_2_ = −0.01; *m* = as in (a).

Despite the large uncertainties about the global effect of selfing on local adaptation, the analysis of the model yields some predictions about the genetic architecture of local adaptation in selfing versus outcrossing species. In outcrossing species, the global adaptation from de novo mutations is predicted to be biased towards dominant mutations, the so-called Hadane’s sieve (Haldane, 1927, Ronfort and Glémin, 2013), and local adaptation to beneficial dominant/deleterious recessive mutations (Yeaman and Otto, 2011). On the contrary, no sieve related to dominance is expected under high selfing. As for other forms of selection, we also predict that local adaptation should be more prominent on female traits because selfing reinforces selection on female fecundity components at the cost of male fecundity components. A similar conclusion was reached by Olito *et al*. (2018) from a model including male/female antagonism and heterogeneous habitat with full dispersal. Finally, beyond the current work, simulations showed that local adaptation genes tend to aggregate into clusters in outcrossers (Yeaman and Whitlock, 2011) whereas the genetic architecture tends to be more diffuse in selfers (Hodgins and Yeaman, 2019, Le Thierry d’Ennequin *et al*., 1999). Thus, a natural extension of the model would be to consider local adaptation at two loci to dissect the additional effect of selfing on genetic linkage. However, using the same formalism would require following a multi-type branching process in a higher dimension (at least six), which remains highly challenging.

## Supporting information

Contains simulator source code and Mathematica notebook that can be used to reproduce all theoretical derivations and figures in the manuscript.

Scripts for running the simulator and processing the raw data, together with the complete output necessary for plotting using Mathematica notebook.

## 5 Data availability

Supplemental files are available at https://github.com/BogiTrick/local_adaptation_single_locus and at Figshare (https://figshare.com/s/683d607a35f45a4978b1). These entries contain: *Mathematica* notebook required for reproduction of all reported theoretical results and figures; C++ source code of the custom simulator used for generation of simulated data for comparison with analytics; Shell scripts for the bulk run of the simulator; and R scripts used to process the raw simulated data for plotting in the above-mentioned notebook.

## 6 Acknowledgements

We thank Matthew Hartfield for the critical reading of the manuscript. B. T. was supported in the initial phase of this project by Erasmus Mundus Joint Masters Degree Scholarship awarded by Erasmus+ programme. Author contributions: B.T. and S.G. developed model. B.T. wrote the code and run the simulations. B.T. and S.G. wrote the manuscript.

## Appendices

### A Deriving the PGF of the distribution of allele offspring number

#### A.1 Single population

We first derive the full probability generating function (PGF) of the number of mutant alleles under partial selfing in a single population. When the *A* mutant is rare, it can be found either in a *Aa* individual with probability 1 − *F* and in a *AA* individual with probability *F*. The PGF can thus be written as (1 − *F*)*f_Aa_*(*z*) + *Ff_AA_*(*z*) where *f_Aa_* and *f_AA_* are the PGF for the parent allele being in each genotype, respectively. We can then decompose the total number of alleles, *G_i_*, contributed by one individual of genotype *i* as the sum of alleles transmitted through outcrossed ovule, *N_i,o_*, through exported pollen, *N_i,p_*, and through selfed ovule, *N_i,s_*. Because the mutant is rare, it can only be transmitted in heterozygote after outcrossing, so in a single copy. After selfing, two copies are always transmitted if the parent is *AA* and either two or one copies are transmitted if the parent is *Aa*. In this last case we note 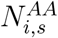 and 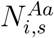 are the number of homozygote and heterozygote seeds produced under selfing. The total number of offspring alleles transmitted at the next generation is thus:

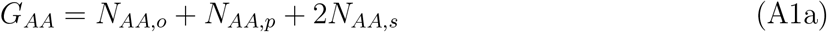

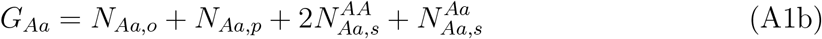

We assume that the number of exported pollen and the different numbers of seeds follow independent Poisson distributions. The different means depends on the fecundity of the parent, the viability of the seed produced and the proportion of each category:

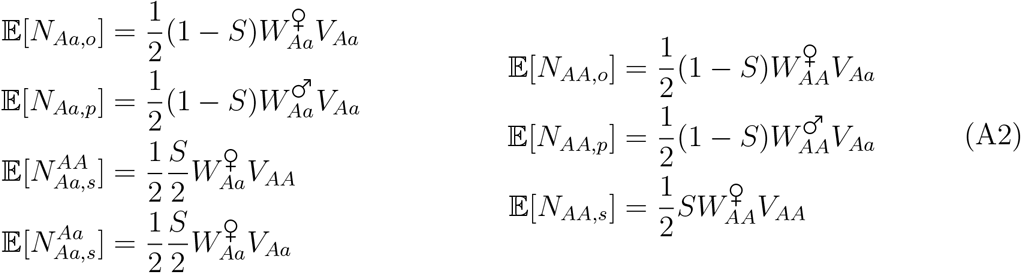

The factor 1/2 corresponds to the fact that the focal *A* is chosen with probability 1/2. Put another way, on average each individual transmits two alleles so the contribution for a single allele must be halved. Finally, we use the property that the PGF of 2*X* is *f*(*z*^2^) where *f*(*z*) is the PGF of *X* and that the PGF of the sum of independent variables is the product of the PGFs. Noting *g*(λ, *z*) = *e*^−(1−*z*)λ^, the PGF of a Poisson distribution with mean λ, we have:

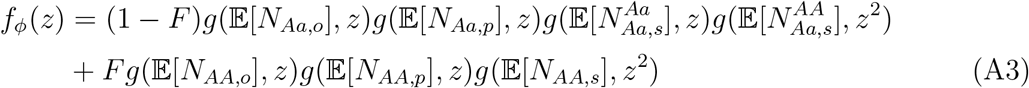

which yields equation (15) of the main text.

From the properties of PGFs we can easily obtain the moments of the distribution for the different forms of selection (see *Mathematica* notebook):

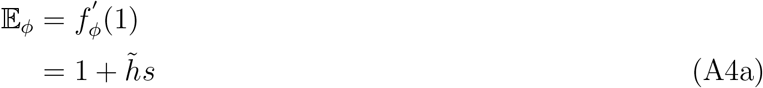

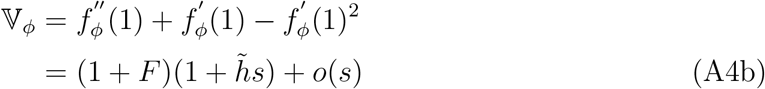

So we retrieved the well known result that selfing increases variance in allele offspring number by 1 + *F*. Interestingly, the full distribution presents a peculiar non-monotonic behavior for high selfing, with an excess (resp. a deficit) in even (resp. odd) numbers. Note also that the full distributions are not exactly the same under the different modes of selection.

#### A.2 Two-patch model

We need to derive the distribution of the number of mutant alleles issued from one patch and staying in the same patch, with PGF *f_i,i_*(*z*), and the distribution of those establishing in the other patch, with PGF *f_i,j_*(*z*). Assuming that the distribution of resident and migrant alleles are independent, the total number of alleles produced by patch *i* has PGF: *f_i_*(*z*) = *f_i,i_*(*z*)*f_i,j_*(*z*). The *f_i,j_*(*z*) can be obtained using the same equations as for a single population by paying attention to the order of events to correctly set indices: male fecundity selection, pollen migration, female fecundity selection, reproduction, seed migration and viability selection. Under seed migration, fecundity selection occurs in patch *i* but viability selection in patch *j*. The means given in equation (A2) become:

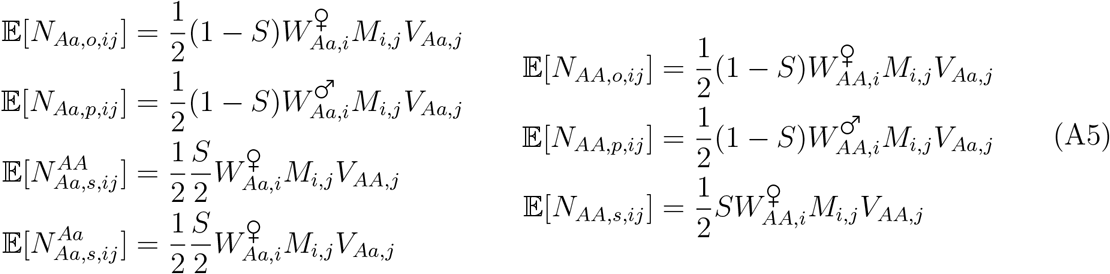

with *M_i,j_* = 1 − *M_i,i_*. Plugging (A5) into (A3) yields equation (16) in the main text. From the corresponding PGFs we then retrieve the same mean as obtained by the deterministic analysis (see below), and variances inflated by 1 + *F* (see *Mathematica* notebook):

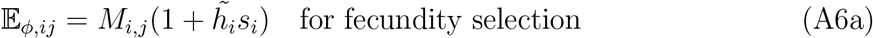

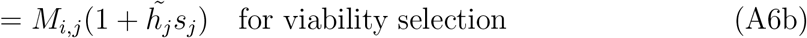

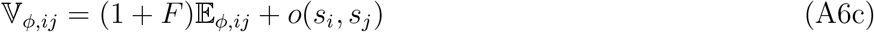

Note that in subsequent analyses the order of migration and selection terms yields the same results. Under pollen migration, the PGF for resident and migrant contribution have different forms because an allele can contribute offspring to the other patch only through outcrossing and through the male pathway. For offspring contributing to the resident patch we have:

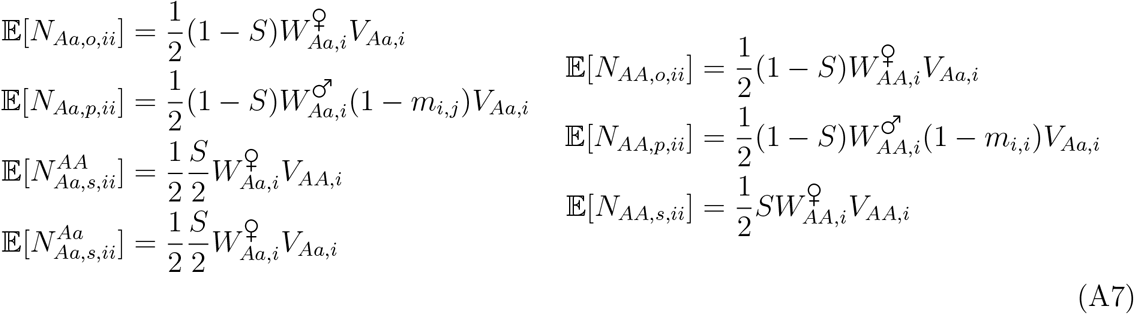

whereas for offspring contributing to the other patch:

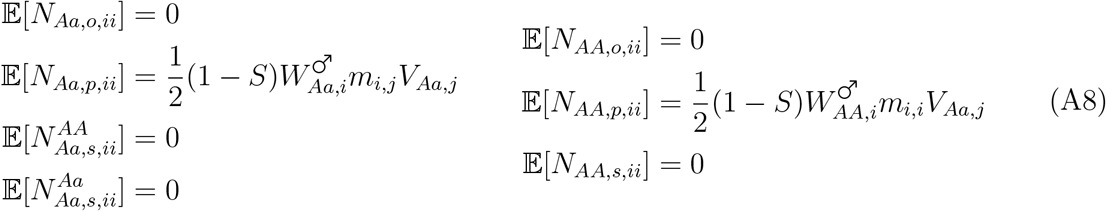

Plugging (A7) and (A8) into (A3) yields equations (17a) and (17b) in the main text. The mean and variance for resident offspring are:

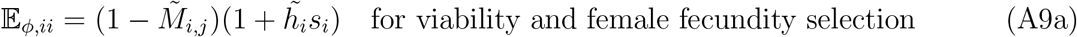

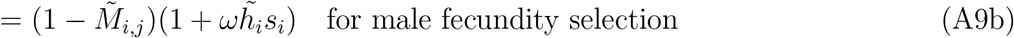

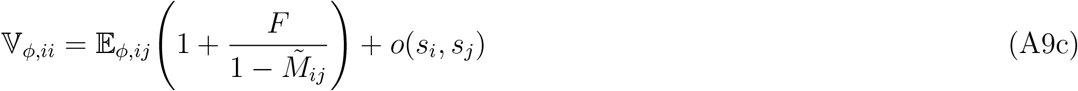

and for migrant offspring:

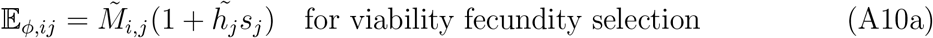

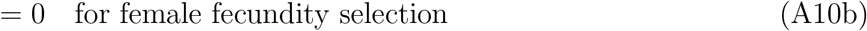

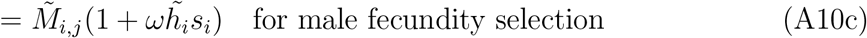

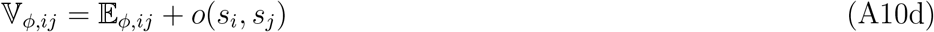

where 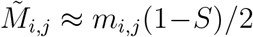 are effective migration rates as defined in the main text (equations 12a to 12c) and *ω* is a correcting factor defined in equation (13). Compared to seed migration, the variance is not uniformly increased by 1 + *F*. As migrant offspring can only be produced through outcrossing, the distribution is simply Poisson and the variance equal to the mean. On the contrary, because the proportion of outcrossed offspring contributing to the resident patch is reduced due to pollen migration, the variance is inflated by more than 1 + *F*. However, the difference between the two modes of migration does not affect the following approximations for weak selection.

### B The establishment probability approximated to weak selection

Following the procedure outlined in Haccou *et al*. (2005) (Section 5.6.2) and previously adapted for a similar problem by Tomasini and Peischl (2018), we seek to approximate *P* by working with slightly supercritical process. Let *ρ* be the leading eigenvalue and **u** = [*u*_1_, *u*_2_]^*T*^ and **v** = [*v*_1_, *v*_2_]^*T*^ normed left and right eigenvectors of of the mean reproductive matrix **M**. Eigenvectors are normalized such that:

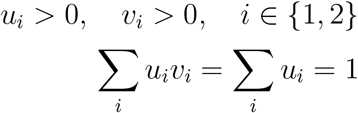

We choose a parameter *ϵ* in the model, such that the process is slightly supercritical when *ϵ* is small. All other parameters in the model are rescaled by *ϵ*. More formally:

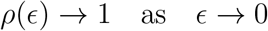

Eigenvalues and eigenvectors of **M** are dependent on *ϵ*. Probability of establishment starting from a single copy of allele in patch *i* is given by:

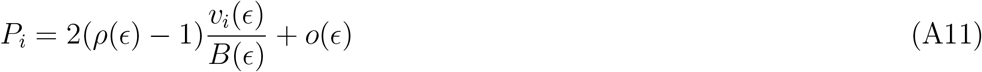

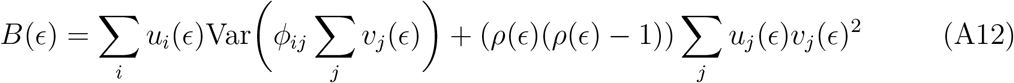

As *ϵ* → 0, *ρ* approaches unity and the second term in B (equation (A12)) can be neglected. As per equation 5.85 in Haccou *et al*. (2005), we take that *B*(*ϵ*) → *B*(0) and *v_i_*(*ϵ*) → *v_i_*(0):

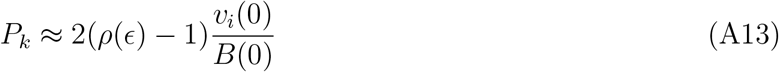

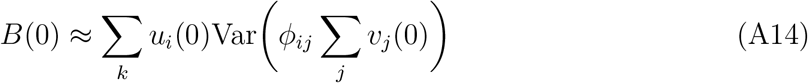

Recall that 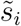 is the advantage of an allele in patch *i* accounting for the effect of self-fertilization. Assuming weak selective advantage, 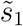 is taken as *ϵ*. All other parameters are expressed in terms of 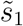:

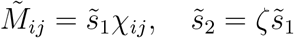

The leading eigenvalue of **M** can be written as 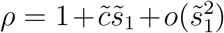. Dropping the higher-order terms in 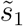, we retrieve the expression for 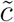:

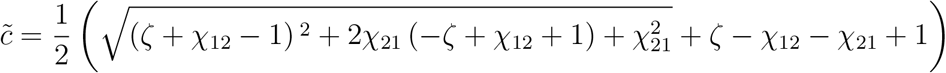

By taking only the zeroth term of *v_i_* and *B* in Taylor expansion about 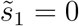, we obtain the approximation of the probability of establishment conditioning on a single mutant appearing in patch *i*:

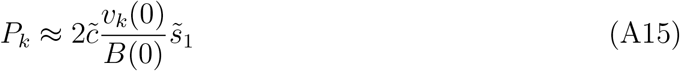

Transforming back to original variables 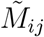 and 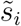 and letting

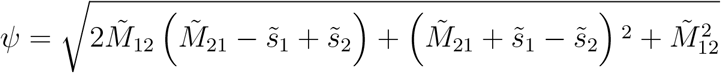

we have:

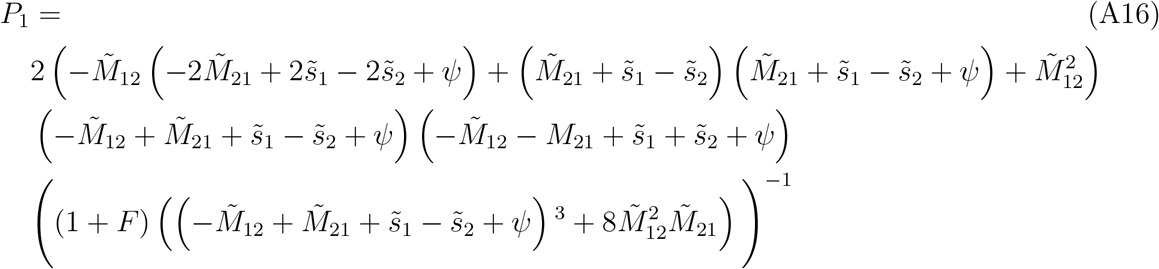

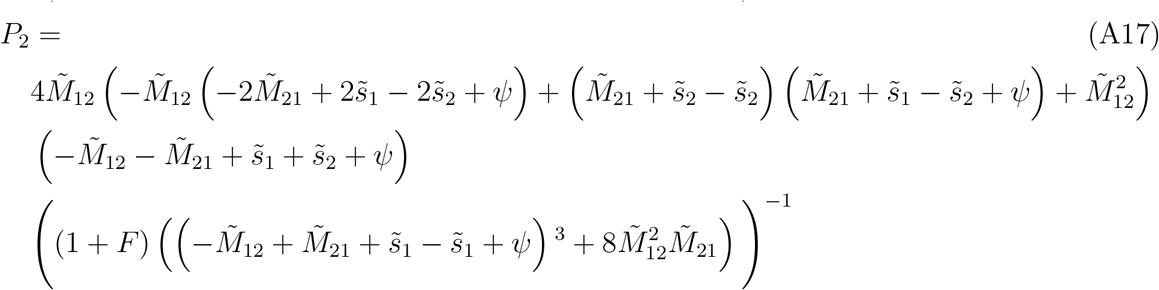

When migration is symmetrical 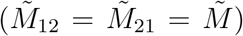, *ψ* reduces to 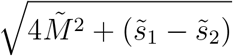, previously defined as the scaled measure of the heterogeneity in selection and migration. Setting *F* = 0, the equations (A16) and (A17) behave similarly to those derived previously (Tomasini and Peischl (2018)). Comparing the analytical solution against simulations, one can see that the solution reported here fits simulations slightly better than the Tomasini-Peischl result (Figure A9). The discrepancy is probably caused by the latter’s use of B term from Aeschbacher and Bürger (2014) (see their equation S22), which neglects to square elements of the eigenvectors after factoring them out of the variance. We, on the other hand, computed B directly from Haccou *et al*. (2005), which does not suffer from this error. Curiously, the Tomasini-Peischl approximation is much more elegant as the denominator can be interpreted as the measure of the heterogeneity of migration and selection. Relative to the result reported in Sakamoto and Innan (2019), our solution had identical performance in the favored patch, and worse performance in the disfavored patch (pink dashed lines). This discrepancy occurs only when the migration rate is low.

**Figure A9:**
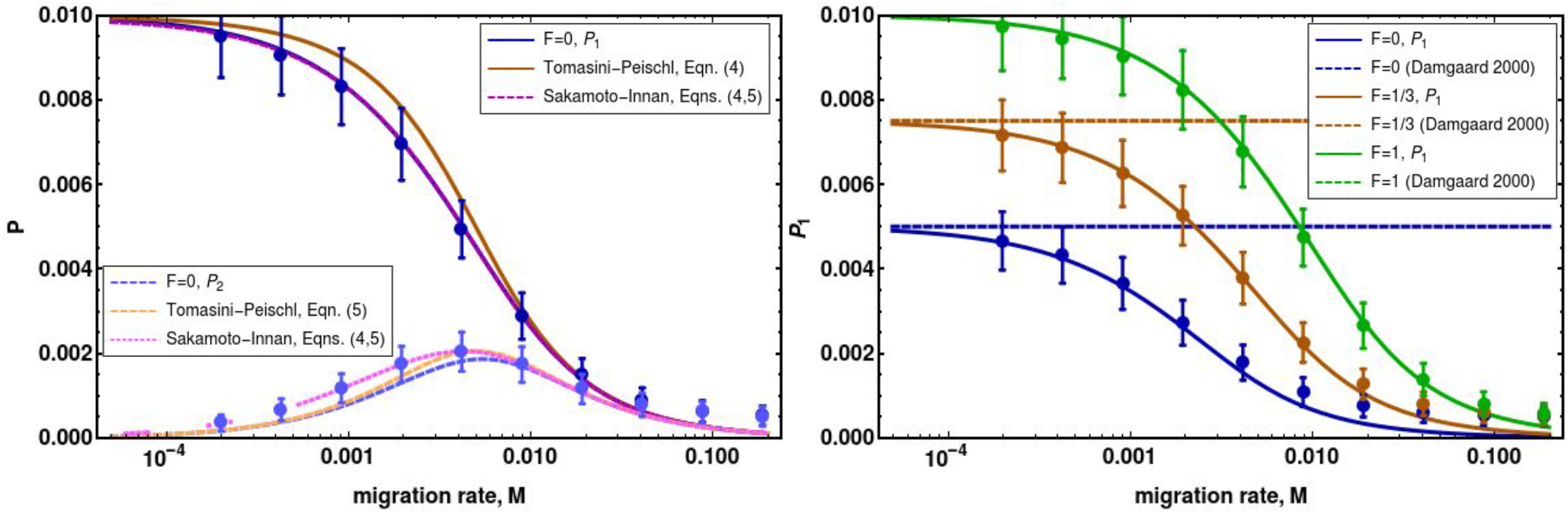
Comparison of analytical solution to simulated data and previous results. Comparison to approximation Tomasini and Peischl (2018) (left), and comparison to the single-patch heuristic under fecundity selection Damgaard (2000) (right). All equations parameterized with 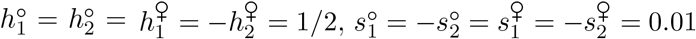, *M*_12_ = *M*_21_ = *M*.

### C The criterion for escaping extinction

#### C.1 Seed dispersal

If the selective disadvantage in the disfavored patch is too large or migration is too strong, the spreading locally advantageous allele can be swamped by its deleterious counterpart. The range of parameters that are necessary but not sufficient for a successful invasion are obtained by linearizing the system about 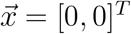 and investigating the conditions required for this equilibrium to be locally unstable. If migration occurs prior to selection – which is the case when seed disperses and selection acts on viability, then Jacobian **J** is:

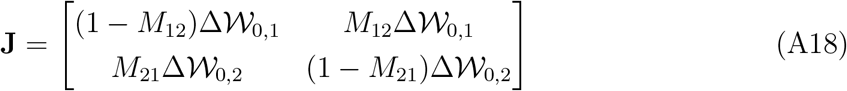

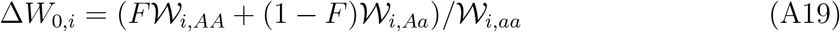

Symbol 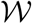 is a place-holder for any of the three selection modes (*V*, *W*^♀^, or *W*^♂^). If selection occurs prior to migration – which happens when seed disperses and selection acts on sexual components– then Jacobian is:

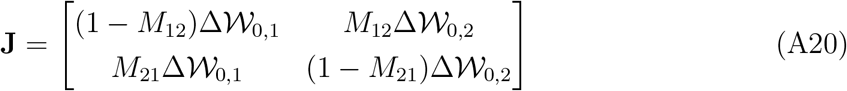

However, the eigenvalues of these two matrices are identical, so we here use (A18). The equilibrium is locally unstable whenever the leading eigenvalue of **J** is greater than unity:

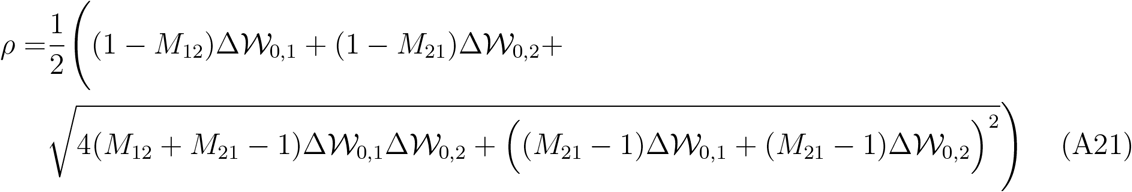

Term 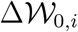 can be thought of as the rate of spread of *A* allele in *i^th^* patch. Then 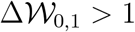 (because *A* is advantageous in the first patch), and 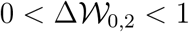 (because *A* is deleterious in the second patch). It is important to note that these inequalities hold regardless of the mode of selection. More formally:

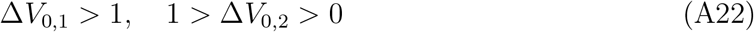

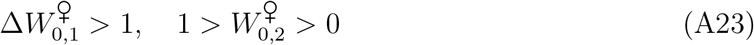

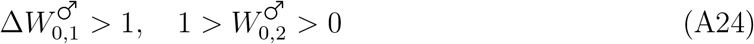

Substituting (A19) in (A21), and rearranging (see *Mathematica* notebook for details), we find that *ρ* > 1 when:

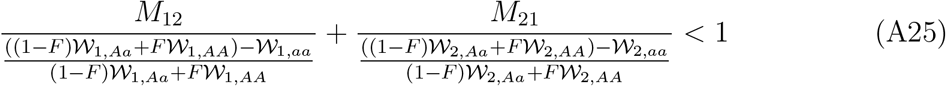

Terms in denominators of inequality A25 represent the relative fitness of the invading allele *A*. Letting *F* = 0 and parameterizing such that 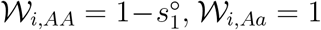, and 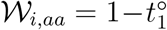, we retrieve Bulmer’s inequality 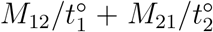. Parameterizing according to our fitness scheme (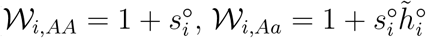, and 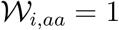) yields:

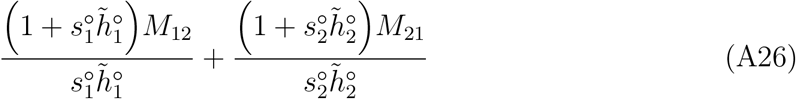

Conversely to the previous case, 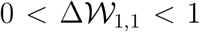 (as invading allele *a* is deleterious in the first patch) and 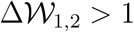 (given that *a* is beneficial in the second patch). Once the allele *A* has escaped extinction, it can either fix in both patches or be maintained for finite number of generations by divergent selection. Thus, the criterion for protected polymorphism is that both allele *A* and *a* can escape extinction. By linearizing the system of replicator equations about 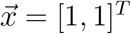 we obtain Jacobian:

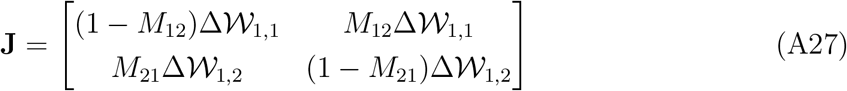

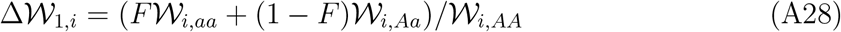

Therefore, allele *a* is allowed to invade whenever:

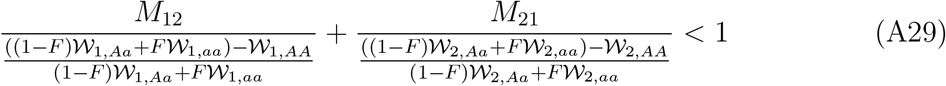

which reduces to 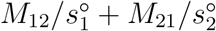 with fitness notation of Bulmer.

#### C.2 Pollen dispersal

The analysis is more complicated for the three selection scenarios when pollen disperses because one has to re-parameterize migration rate in addition to dominance coefficients. We use Jacobian of the form:

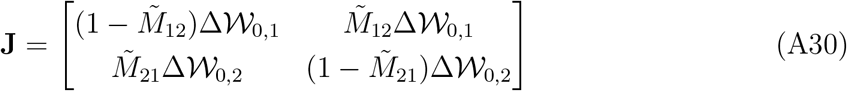

The leading eigenvalue of (A30) is:

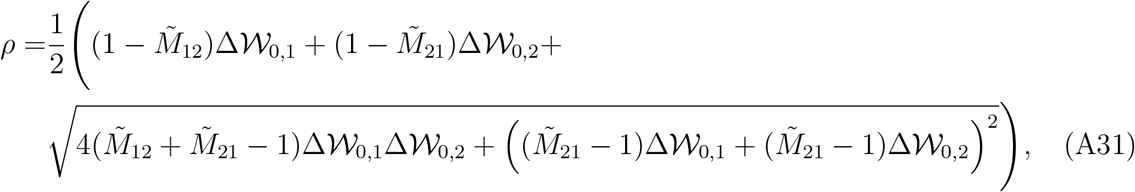

where 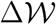 and 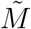 are parameterized as outlined in Section 2.2. Given that 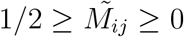 for all *i* and *j*, and

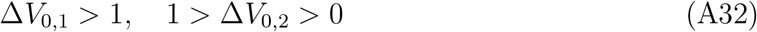

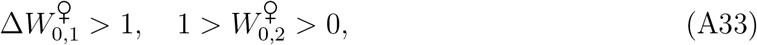

the same inequalities hold as in the case of seed dispersal. However, under male fecundity selection migration rates have to be:

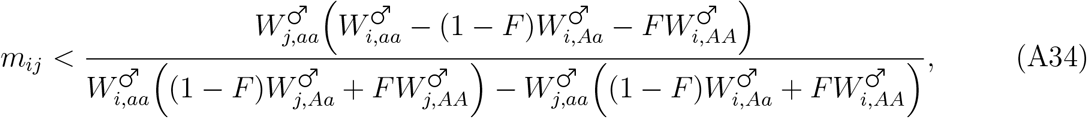

for both *m*_12_ and *M*_21_, or:

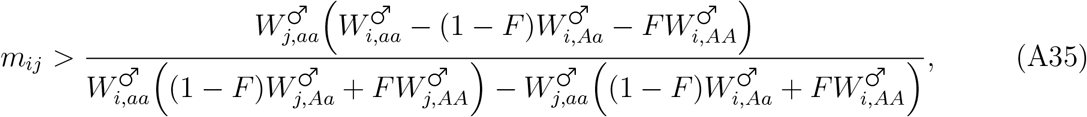

for both *m*_12_ and *m*_21_. The conditions (A34) and (A35) are identical to saying that *ω_i_* > 0 for all *i*, or *ω_i_* < 0 for all *i*. A possible intuitive explanation for these conditions is as follows. If migration from favored to disfavored patch is too high, the mutant alleles are transferred to a disfavored patch where they are purged, thus causing the mutant to go extinct. If, on the other hand, migration from disfavored to favored patch is high, then the spreading mutant is swamped by the influx of deleterious residents from the opposite patch.

### D Simulation method

Selection and migration are assumed to alter genotype frequencies deterministically. Genetic drift is implemented by randomly drawing genotypes from multinomial distribution right after reproduction. We relax the assumption that the population has to reach equilibrium in the inbreeding coefficient *F*. Each simulation run terminates in a successful or failed invasion and is composed of the following four steps:

1. Inject a single heterozygote containing allele *A* in a population that is fixed for *a* allele. When interested in *P_i_*, the mutant is injected in *i^th^* patch.
2. Update genotype frequencies by applying equations (2a)–(4b); This emulates selection on sexual components, pollen dispersal, and reproduction (including selfing).
3. Sample the genotypes from multinomial distribution to determine the genotype frequencies after the reproduction step: 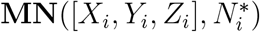, where 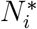 is the size of *i^th^* patch.
4. Update genotype frequencies due to seed dispersal by implementing (5a) and (5b).
5. Compute the number of each genotype after viability selection as 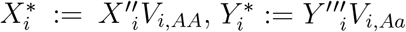, and 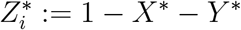.
6. Is the number of invading mutant alleles larger than 1000? If not, convert genotype numbers to genotype frequencies by dividing drawn genotype numbers with the total patch size and begin the new generation by going to step 2. If yes, count it as a successful invasion and terminate the simulation run.

If the mutant allele does not invade in 10,000 generations, we count the simulation run as a failed invasion. The establishment probability is obtained by running 10,000 simulations and computing the fraction of runs that ended in the successful invasion. All error bars in the figures correspond to the standard deviation of this metric. The chosen number for invasion threshold is well over 1/*s*_1_, given that we use *s*_1_ = 0.01. Both patches contain 10,000 individuals, and populations are always in |*N_s_*| ≫ 1 regime.

### E Simulations under various selection and migration modes

Our approximation gives a good fit for simulated data under female and male fecundity, although the fit is not excellent as in the case of viability selection (Figures A10–A13). In the case of male fecundity selection under complete selfing (*S* = 1), the male component does not contribute to the fitness, and the allele is expected to behave neutrally. One can see that this occurs because the probability that the invading allele reaches the threshold that we use to determine whether the invasion is successful is inversely proportional to the size of the threshold, 1/*N*_thres_, (grey dashed line in figures below). The figures below compare the analytics to simulated data, with upper row denoting the case when allele originates in the favored patch, and bottom row depicts the same dynamic but when allele appears in the disfavored patch. From left to right, columns show codominant, partially dominant, and partially recessive case, respectively.

**Figure A10:**
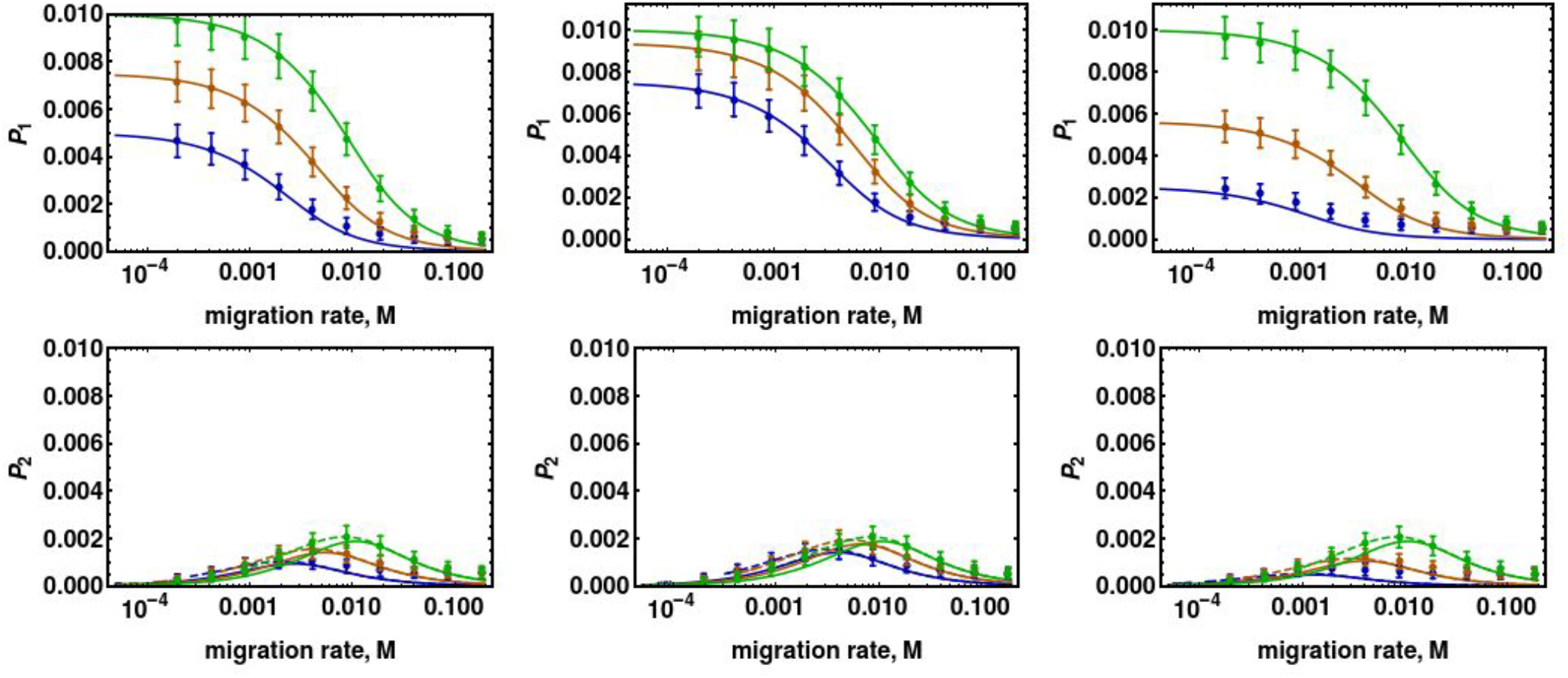
Comparison of analytical solution to simulations when selection acts on the female fecundity and only seed disperses. Left column: codominant case 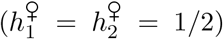; Middle column: dominant case 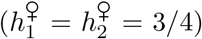; Right column: recessive case 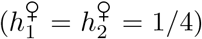. Upper and lower panels depict the establishment probability conditioning on allele emerging in favored and disfavored patch, respecitvely. Parameters: 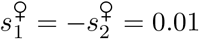, *M*_12_ = *M*_21_ = *M*.

**Figure A11:**
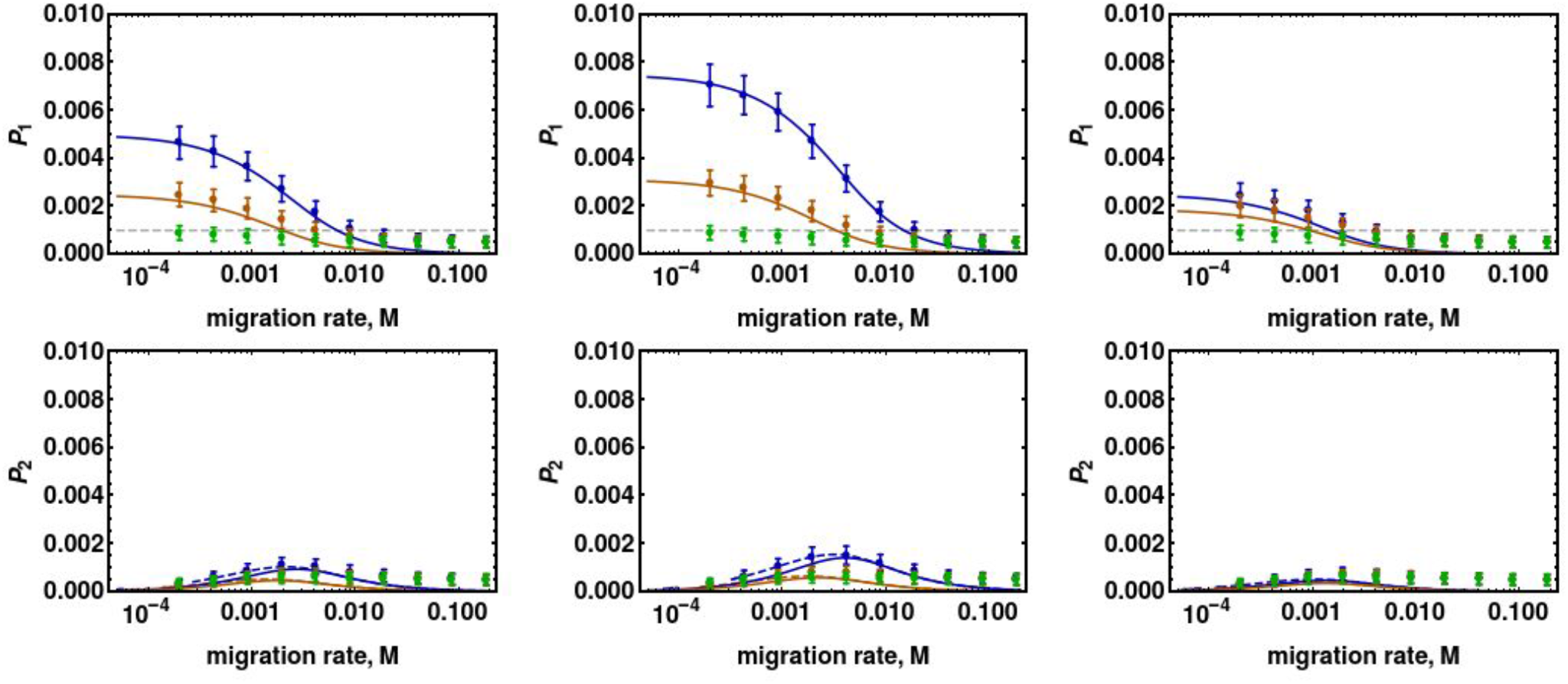
Comparison of analytical solution to simulations when selection acts on male fecundity and only seed disperses. Left column: codominant case 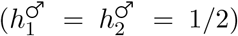; Middle column: dominant case 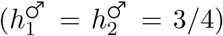; Right column: recessive case 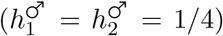. Upper and lower panels depict the establishment probability conditioning on allele emerging in favored and disfavored patch, respectively. Parameters: 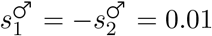, *M*_12_ = *M*_21_ = *M*.

**Figure A12:**
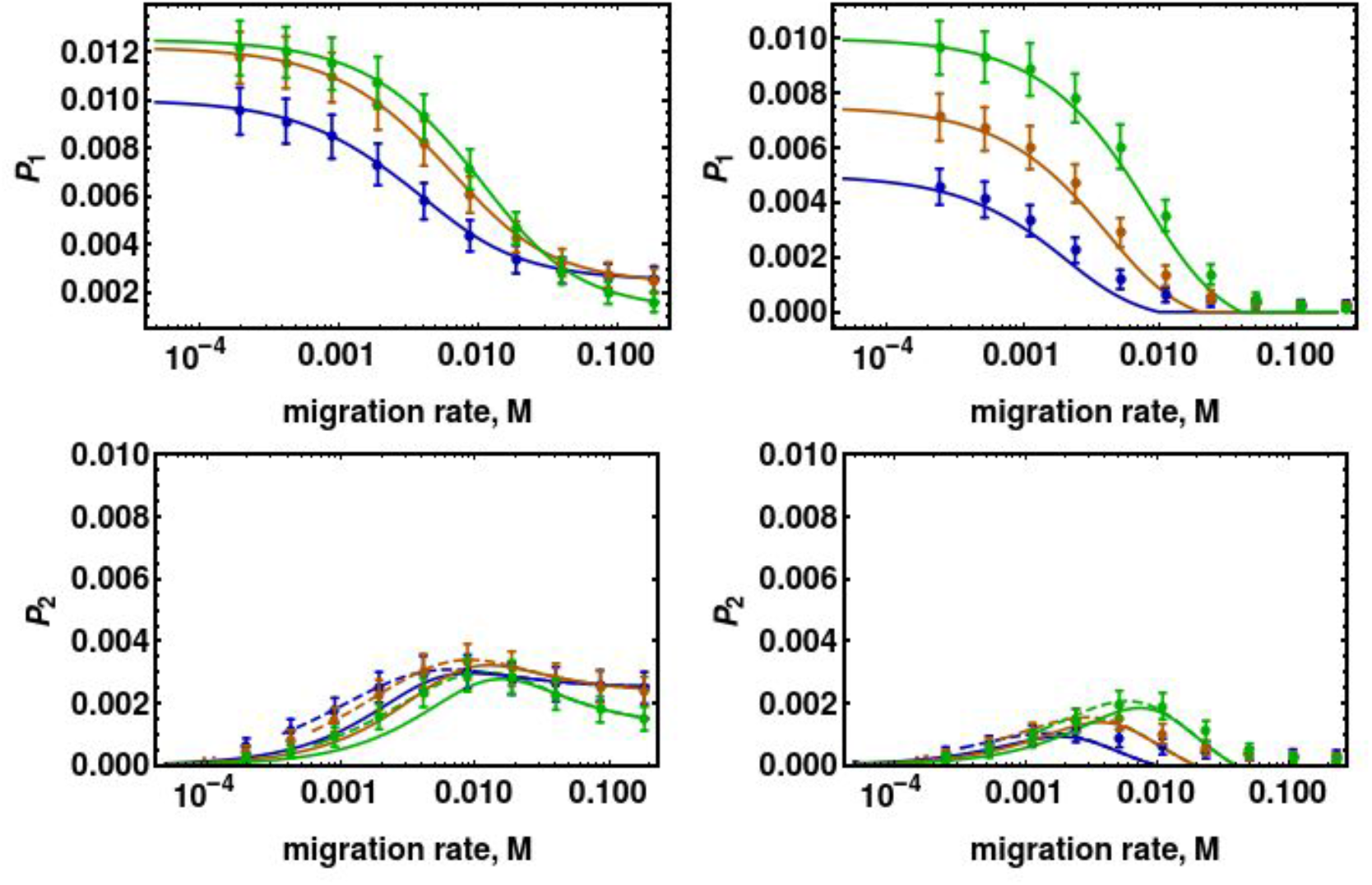
Comparison of analytical solution to simulations when selection acts on the female fecundity, only seed disperses, and selection or migration are asymmetrical. Left column: asymmetric selection 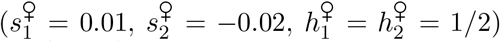; Right column: asymmetric migration (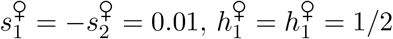, *M*_12_ = *M*, and *M*_21_ = 1.25*M*).

**Figure A13:**
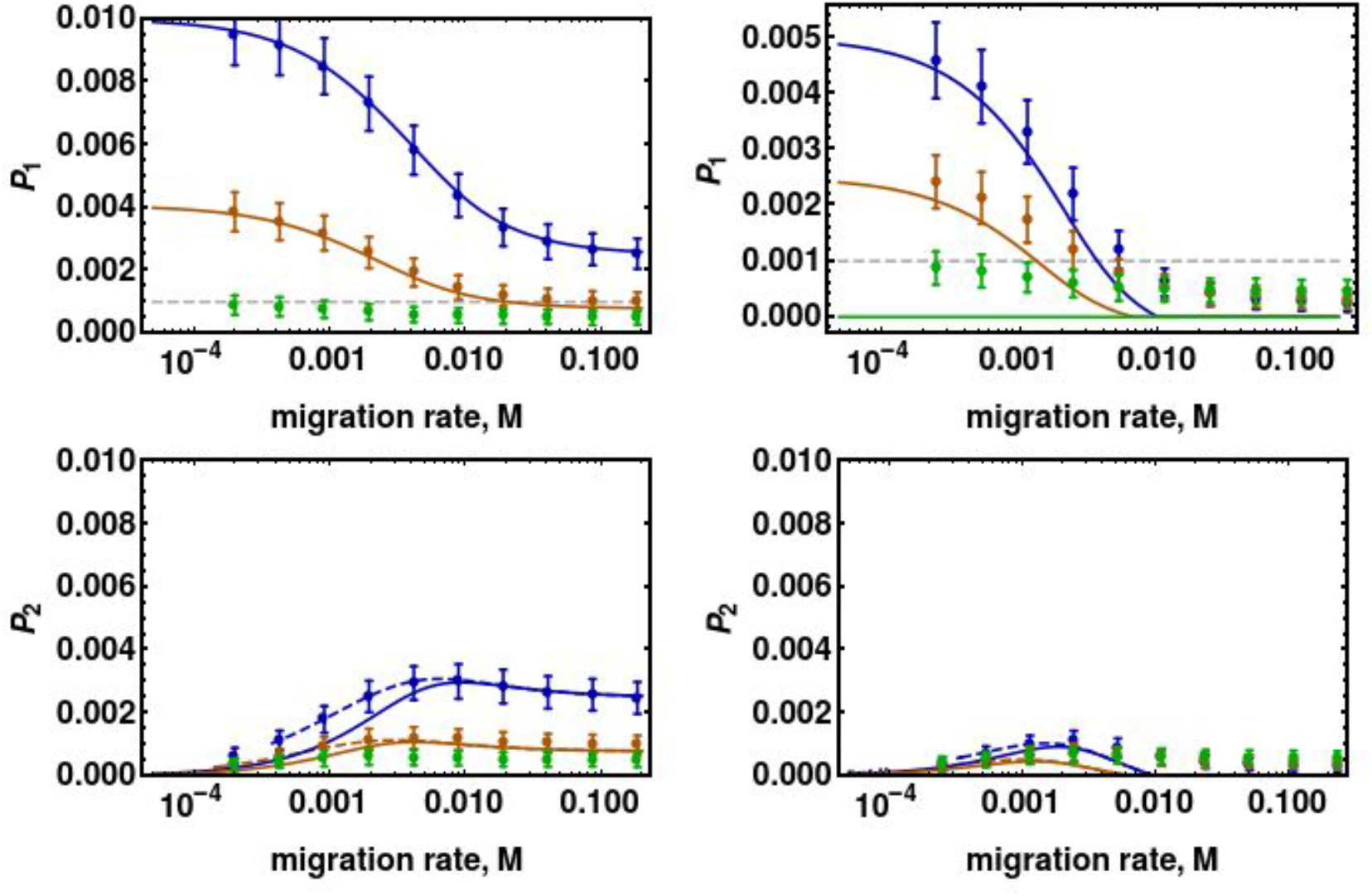
Comparison of analytical solution to simulations when selection acts on the male fecundity, only seed migrates, and selection or migration are asymmetrical. Left column: asymmetric selection 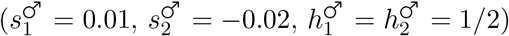; Right column: asymmetric migration (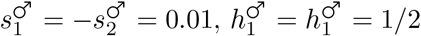, *M*_12_ = *M*, and *M*_21_ = 1.25*M*).

**Figure A14:**
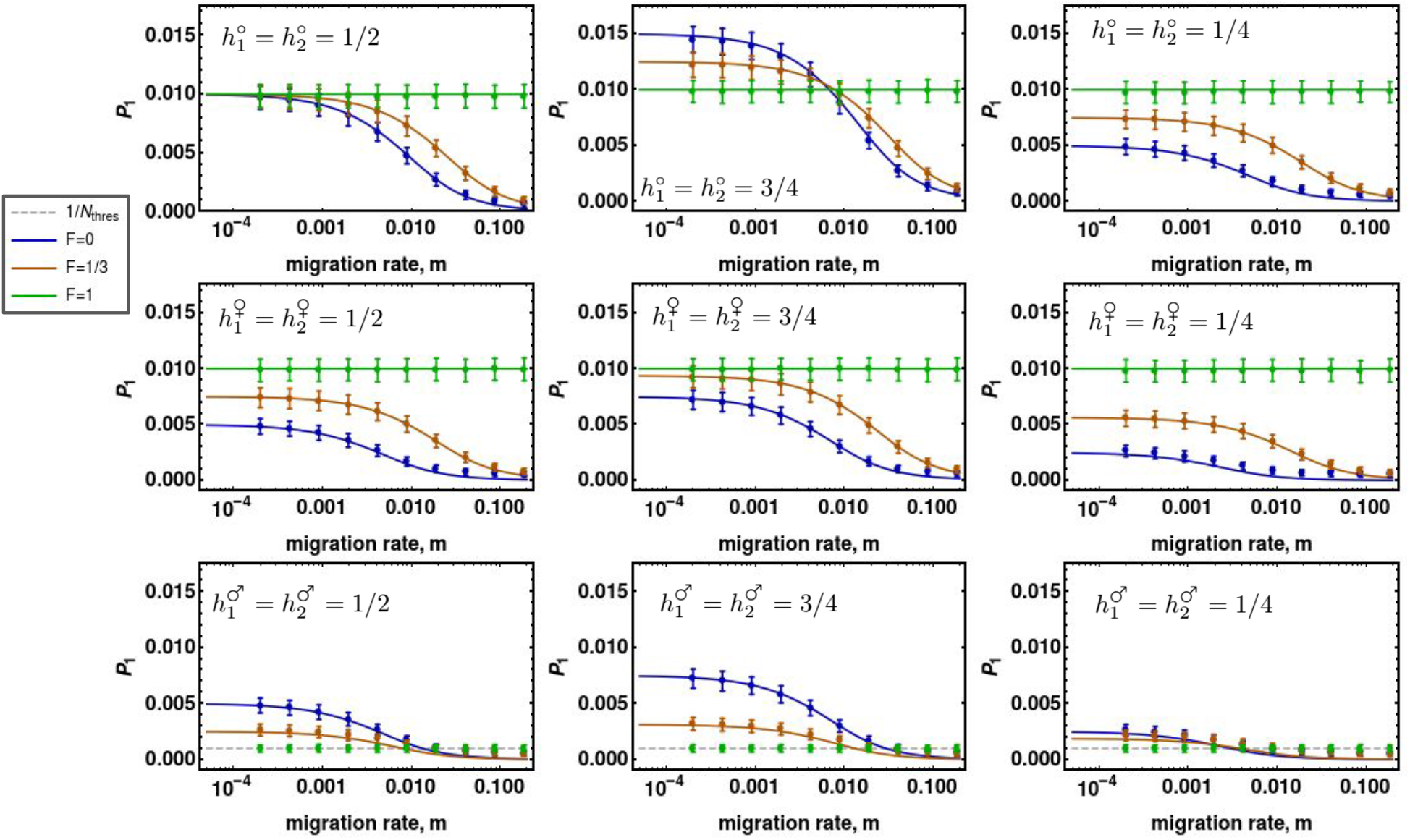
Comparison of analytical solution to simulations when selection acts on the viability and only pollen disperses. Allele starts in the favored patch. Top row: viability selection; Middle row: female fecundity selection; Bottom row: Male fecundity selection. Dominances reported in panels. Other parameters: corresponding seelection coefficients are always *s*_1_ = −*s*_2_ = 0.01, and *m*_12_ = *m*_21_ = *m*.

**Figure A15:**
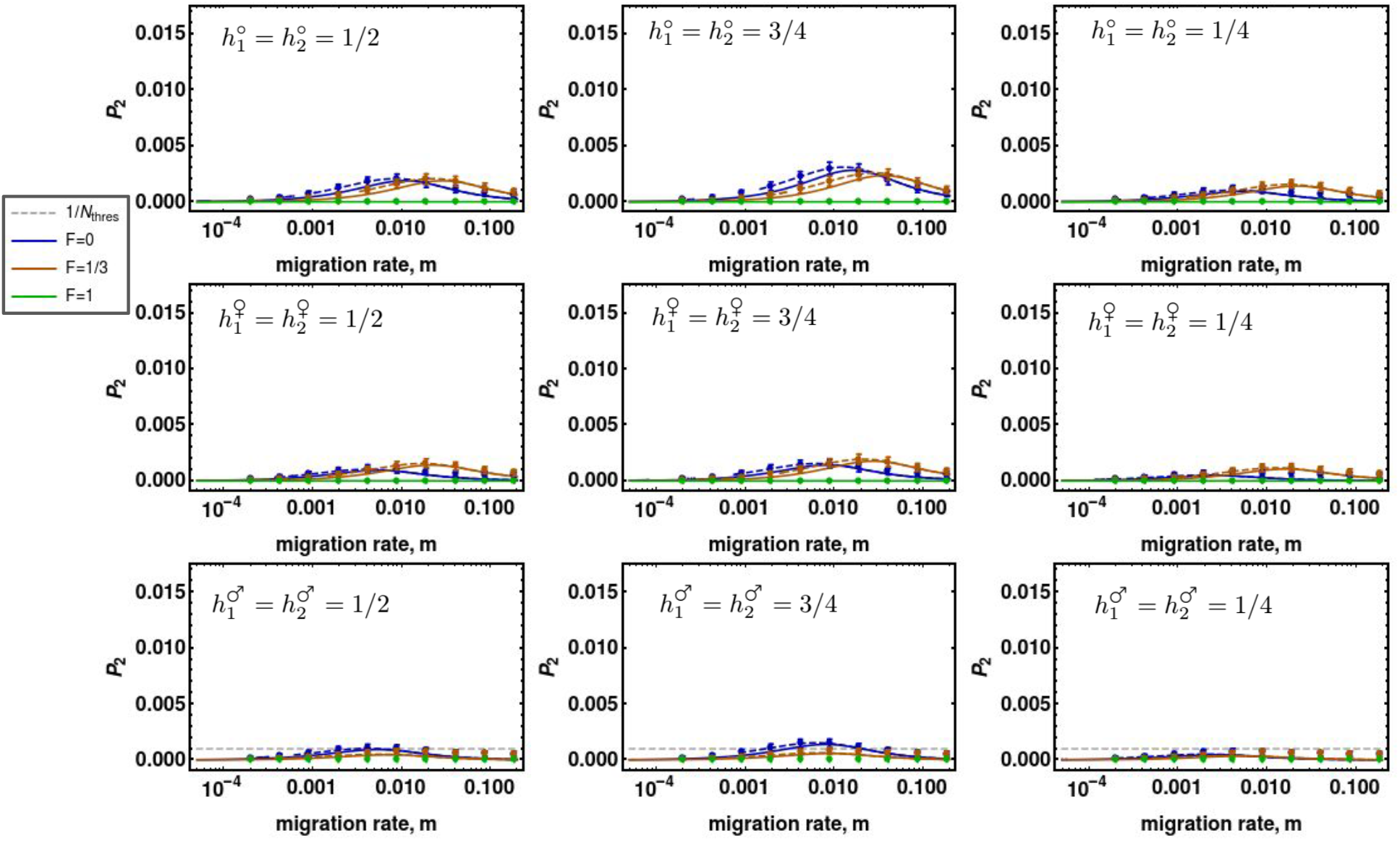
Same as above, but allele starts in the disfavored patch.

### F Procedure for computing *β* indicators

Suppose one wants to derive *β* indicator when selection operates on *j^th^* fitness component (where *j* ∈ {*V, F, M*}) in *k^th^* patch (where *k* ∈ {1, 2}). Selfing affects the establishment probability through three factors: the effective population size 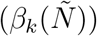, effective favored and disfavored dominance (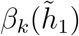 and 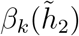), and the effective migration rate 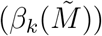. Starting from equations (A16) and (A17), expressions for *β* are obtained using the following steps in a sequential manner:

- 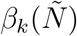: Parameterize migration according to eqns. 12a–12c, and then set *F* = 0 to exclude the effect via 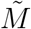. Let 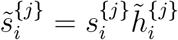 for *i* ∈ {1, 2}. Next, set *F* = 0 to exclude the effect on dominances. Take a derivative in *F* and evaluate at *F* = 0.
- 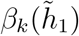: Parameterize migration according to eqns. 12a–12c, and then set *F* = 0 to exclude the effect via 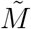. Set *F* = 0 to exclude the effect via 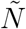. Let 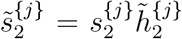 and then set *F* = 0 to exclude the effect via 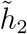. Let 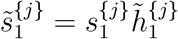. Take a derivative in *F* and evaluate at *F* = 0.
- 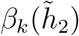: Parameterize migration according to eqns. 12a–12c, and then set *F* = 0 to exclude the effect via 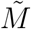. Set *F* = 0 to exclude the effect via 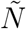. Let 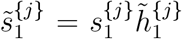 and then set *F* = 0 to exclude the effect via 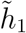. Let 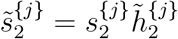. Take a derivative in *F* and evaluate at *F* = 0.
- 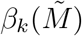: Set *F* = 0 to exclude the effect via 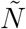. Let 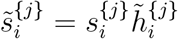 for *i* ∈ {1, 2}. Then set *F* = 0 to exclude the effect via 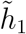, and 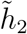. Next, parameterize migration according to eqns. 12a–12c. Take a derivative in *F* and evaluate at *F* = 0.

**Figure A16:**
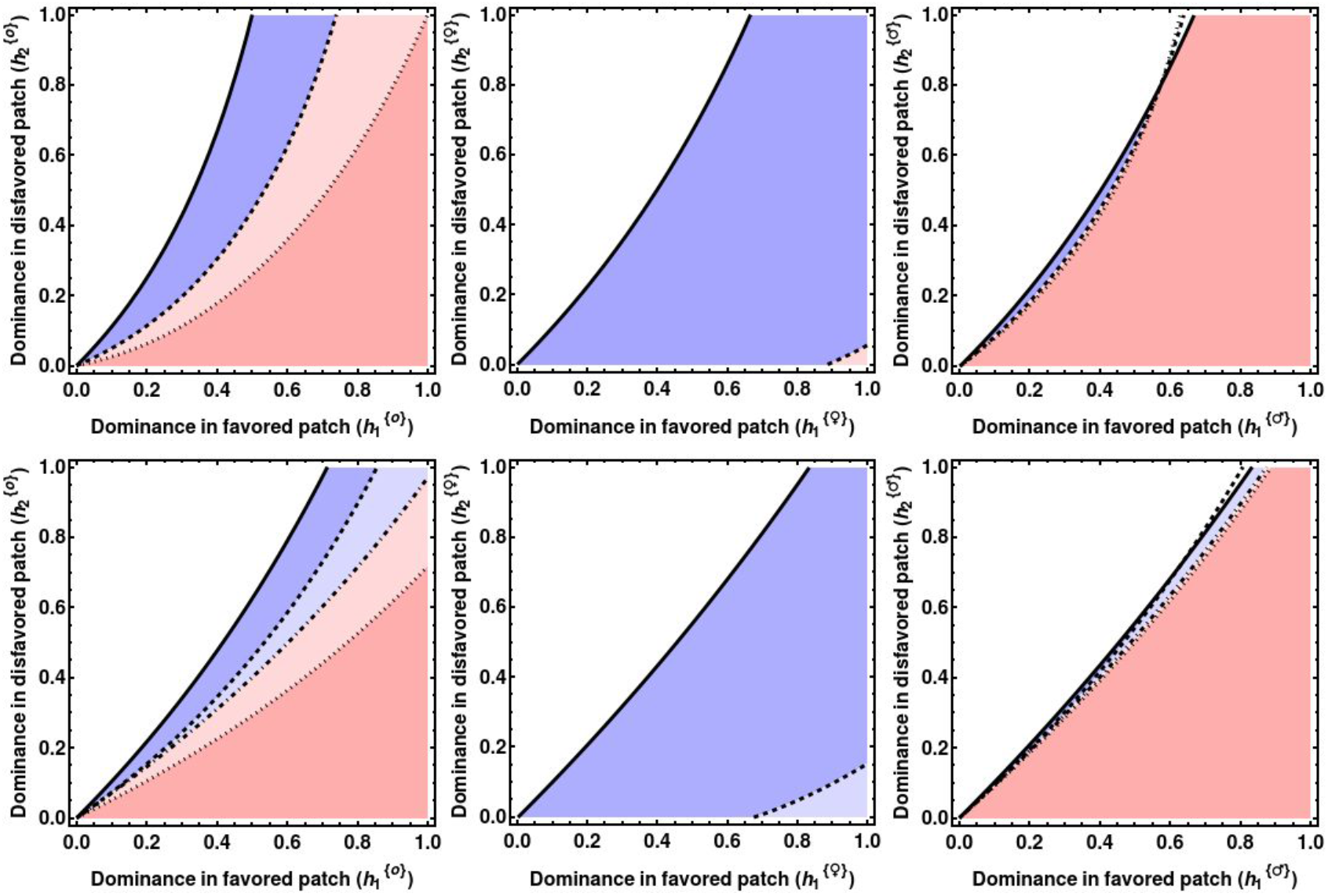
Zones of selfing’s effect on the probability of establishment across viability (left panel), female fecundity (middle panel), and male fecundity (right panel), conditioning on allele appearing in the disfavored patch. Only seed migrates. Parameters are identical to those used in Figure 4. Upper row: scenarios with seed migration; Bottom row: scenarios with pollen migration.

Parameters 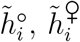, and 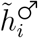 are given by equations (9a)–(9c). Whenever a *β_i_*(*x*) is greater than zero, a shift to selfing increases the probability that an allele becomes established, conditioning on starting in patch *i*. Evaluating the derivative in point other than *F* = 0 will change the results quantitatively, but not qualitatively.

### G The consequences of shift to selfing on local adaptation

Where fecundity selection is examined, the establishment probability equations were parameterized with 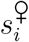, and 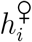, while selection on male fitness component was done by parameterizing with 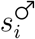, and 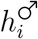. Other than that, parameters were quantitatively identical to the viability selection case. Migration rates were also kept constant across different scenarios, and only the migration type has changed.

**Figure A17:**
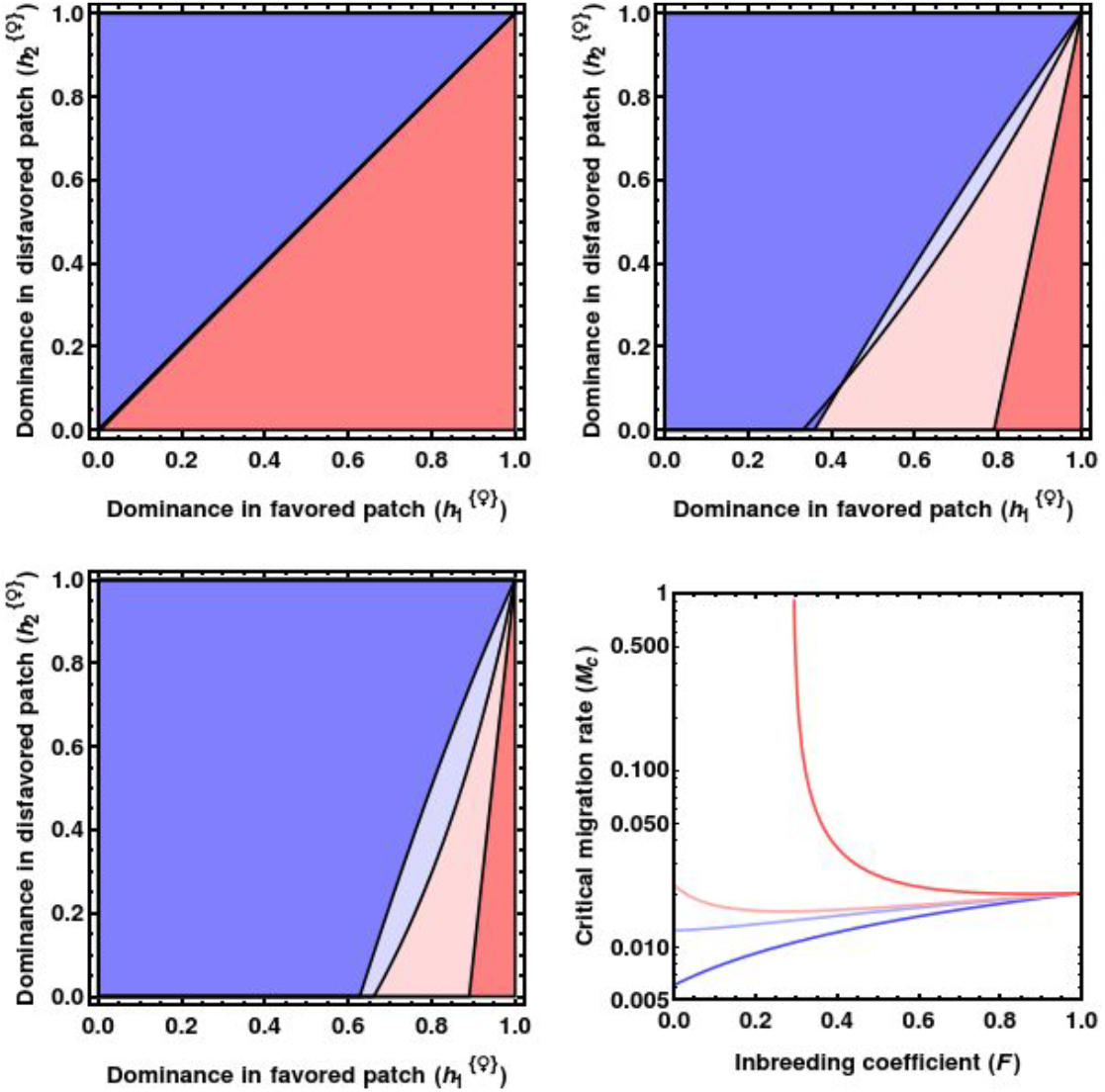
The consequences of a shift to selfing on establishment of local adaptation under female fecundity selection and seed dispersal. Color-coding and parameters as in the main text and parameters as in the main text.

**Figure A18:**
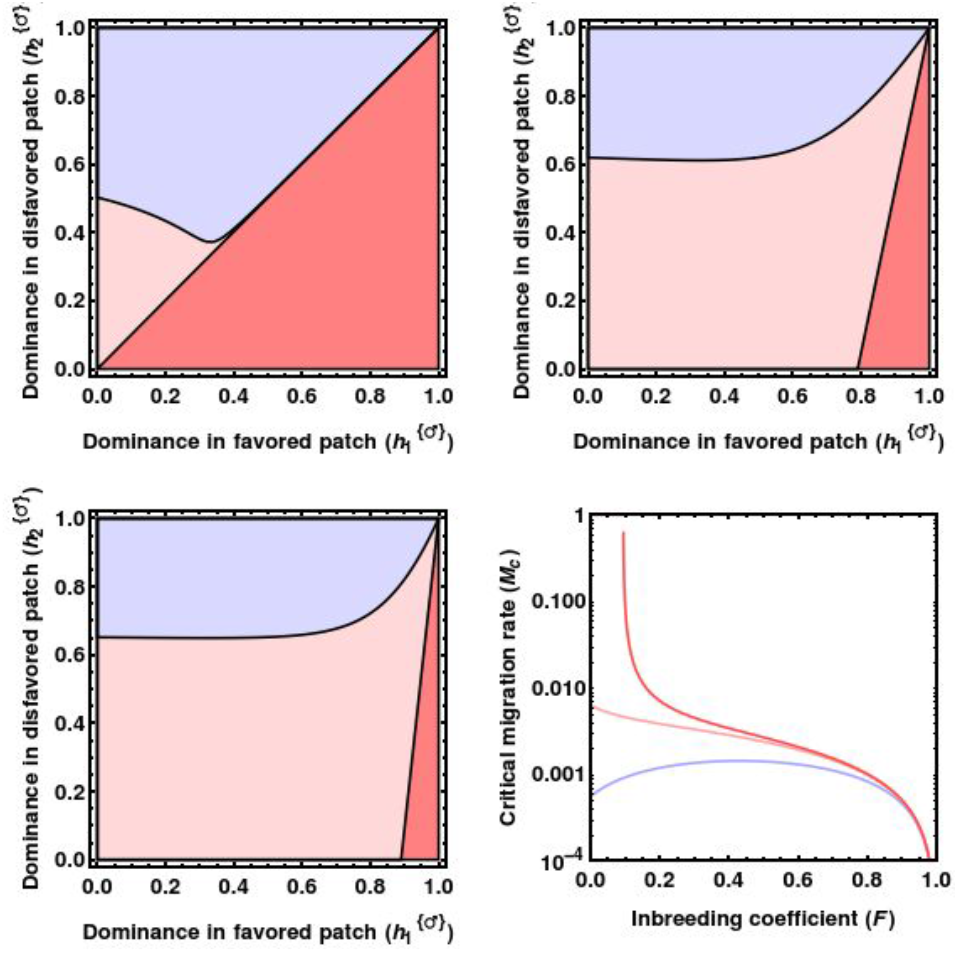
The consequences of a shift to selfing on establishment of local adaptation under male fecundity selection and seed dispersal. Color-coding and parameters as in the main text and parameters as in the main text.

**Figure A19:**
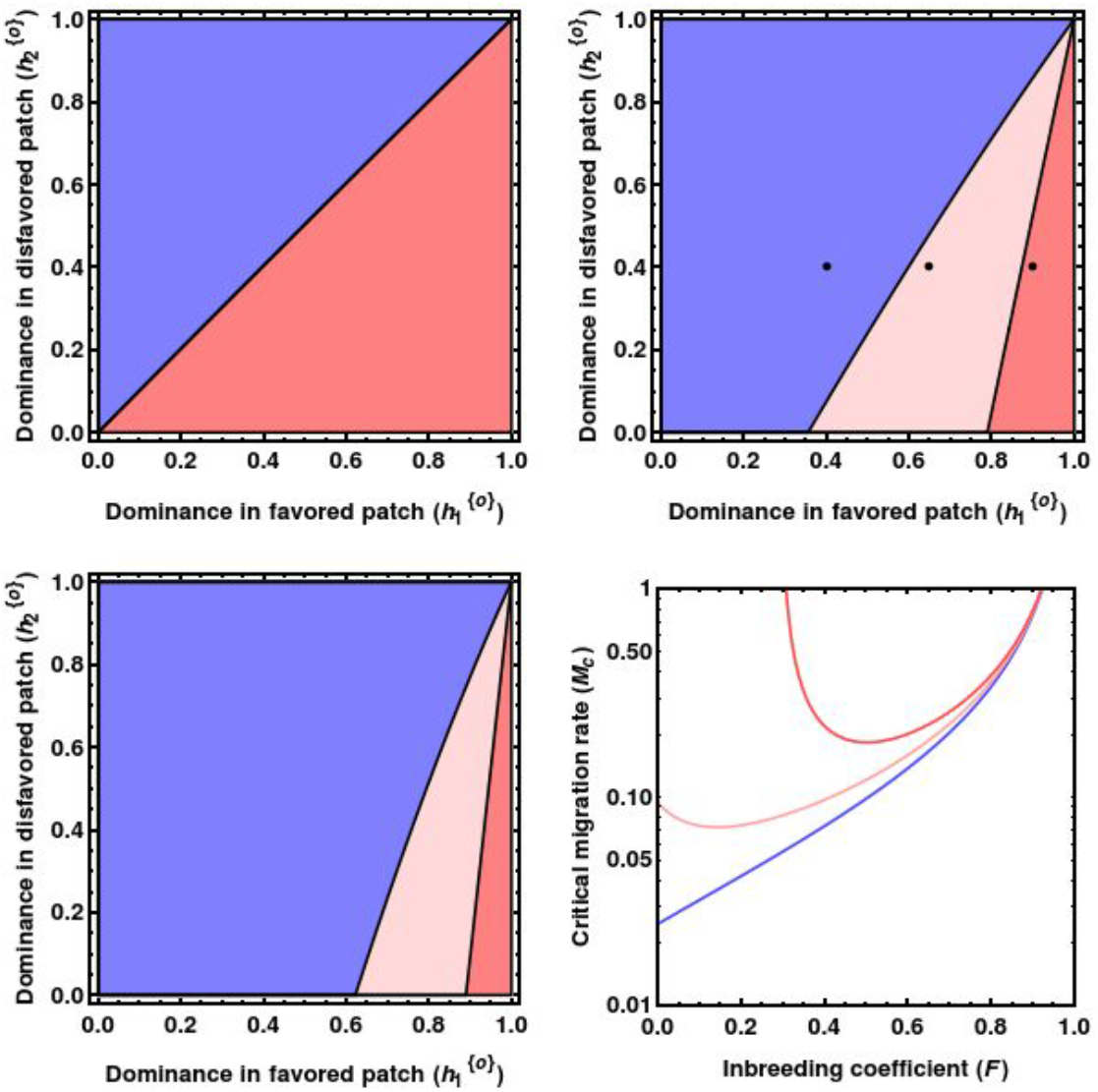
The consequences of a shift to selfing on establishment of local adaptation under viability selection and pollen dispersal.

### H Notation glossary

For the sake of clarity, Table 1 indicates the meaning of symbols used in the derivation of the deterministic model. As a general rule, parameters with tilde signify an *effective* parameter that accounts for the effect of selfing, selection mode, and migration type on the dynamic. In other words, the selfing population with a specific parameter behaves like an outcrossing population with an effective parameter. Upper-case symbols are reserved for processes affecting the diploid phase of a life cycle, whereas the lower-case represents the process of the haploid phase.

