## Supplementary material for "Establishment of local adaptation in partly self-fertilizing populations": Contains simulator source code and Mathematica notebook that can be used to reproduce all theoretical derivations and figures in the manuscript.: main.cpp

```
#include <stdio.h>
#include <stdlib.h>
#include <iostream>
#include <gsl/gsl_rng.h>
#include <gsl/gsl_randist.h>
#include <time.h>
#include <string.h>
#include <math.h>
#include <stdbool.h>

#define patch_nb 2
#define genotype_nb 3


int main(int argc, char *argv[])
{
    (void)argc;
    unsigned int i, j, k, u;
    unsigned int patch_size[patch_nb];

    double mean_w;

    double gtype_freq[patch_nb][genotype_nb], offspring_self[genotype_nb], offspring_outcross[genotype_nb];
    double w[patch_nb][genotype_nb], w_gphyte_female[patch_nb][genotype_nb], w_gphyte_male[patch_nb][genotype_nb];
    double wA_gamete_female[patch_nb], wa_gamete_female[patch_nb], wA_gamete_male[patch_nb], wa_gamete_male[patch_nb];
    double w_gphyte_female_mean[patch_nb], w_gphyte_male_mean[patch_nb], w_female_mean[patch_nb], w_male_mean[patch_nb];
    double allele_freq_female_gamete[patch_nb], allele_freq_male_gamete[patch_nb];
    double gametophyte_female[patch_nb], gametophyte_male[patch_nb];
    double transit_frame[patch_nb][genotype_nb];
    unsigned int offspring_full[genotype_nb];

    double w_A_F_, w_a_F_, w_AA_F_, w_Aa_F_;
    double w_A_m_, w_a_m_;

    double sigma;
    double mig_rate_pollen[patch_nb], mig_rate_seed[patch_nb];
    double allele_frame[patch_nb]; // define forward and backward migration rates

    double invader_nb;
    int mutant_init[2];

    bool exit_;
    double invasion_thresh;

    unsigned int invasion_count=0;
    unsigned int time_threshold=10000;

    unsigned int run_nb;

    // Reading variables
    sscanf (argv[1], "%lf", &w[0][0]);
    sscanf (argv[2], "%lf", &w[0][1]);
    sscanf (argv[3], "%lf", &w[0][2]);
    sscanf (argv[4], "%lf", &w[1][0]);
    sscanf (argv[5], "%lf", &w[1][1]);
    sscanf (argv[6], "%lf", &w[1][2]);

    sscanf (argv[7], "%lf", &w_gphyte_female[0][0]);
    sscanf (argv[8], "%lf", &w_gphyte_female[0][1]);
    sscanf (argv[9], "%lf", &w_gphyte_female[0][2]);
    sscanf (argv[10], "%lf", &w_gphyte_female[1][0]);
    sscanf (argv[11], "%lf", &w_gphyte_female[1][1]);
    sscanf (argv[12], "%lf", &w_gphyte_female[1][2]);

    sscanf (argv[13], "%lf", &w_gphyte_male[0][0]);
    sscanf (argv[14], "%lf", &w_gphyte_male[0][1]);
    sscanf (argv[15], "%lf", &w_gphyte_male[0][2]);
    sscanf (argv[16], "%lf", &w_gphyte_male[1][0]);
    sscanf (argv[17], "%lf", &w_gphyte_male[1][1]);
    sscanf (argv[18], "%lf", &w_gphyte_male[1][2]);

    sscanf (argv[19], "%lf", &wA_gamete_female[0]);
    sscanf (argv[20], "%lf", &wa_gamete_female[0]);
    sscanf (argv[21], "%lf", &wA_gamete_female[1]);
    sscanf (argv[22], "%lf", &wa_gamete_female[1]);

    sscanf (argv[23], "%lf", &wA_gamete_male[0]);
    sscanf (argv[24], "%lf", &wa_gamete_male[0]);
    sscanf (argv[25], "%lf", &wA_gamete_male[1]);
    sscanf (argv[26], "%lf", &wa_gamete_male[1]);

    sscanf (argv[27], "%lf", &mig_rate_pollen[0]);
    sscanf (argv[28], "%lf", &mig_rate_pollen[1]);

    sscanf (argv[29], "%lf", &mig_rate_seed[0]);
    sscanf (argv[30], "%lf", &mig_rate_seed[1]);

    sscanf (argv[31], "%lf", &sigma);

    sscanf (argv[32], "%u", &patch_size[0]);
    sscanf (argv[33], "%u", &patch_size[1]);

    sscanf (argv[34], "%u", &mutant_init[0]);
    sscanf (argv[35], "%u", &mutant_init[1]);

    sscanf (argv[36], "%u", &run_nb);

    invasion_thresh=1000;

    /* Seed the random number generator */
    gsl_rng *gBaseRand;
    int randSeed=10000;
    gBaseRand=gsl_rng_alloc(gsl_rng_mt19937);
    sscanf (argv[37], "%u", &randSeed);
    gsl_rng_set(gBaseRand,randSeed);

    unsigned int time_invasion=0;

    // Each iteration through the loop is a single simulation run
    for (u=0; u<run_nb; u++) {

        // Seed the population with number of mutants specified in input (i.e., mutant_init[patch])
        // We are tracking genotype frequencies, so convert mutant numbers to frequencies
        gtype_freq[0][0]=0;
        gtype_freq[0][1]=(double)mutant_init[0]/(double)patch_size[0];
        gtype_freq[0][2]=1-gtype_freq[0][1];

        gtype_freq[1][0]=0;
        gtype_freq[1][1]=(double)mutant_init[1]/(double)patch_size[1];
        gtype_freq[1][2]=1-gtype_freq[1][1];

        exit_=false;
        unsigned int t=0;

        // Set elements of array holding offspring numbers to zero
        for (i=0;i<genotype_nb;i++) {
            offspring_self[i]=0;
            offspring_outcross[i]=0;
        }

        while (exit_==false) {
                /* ================================== Gamete production dispersal ================================== */
                for (i = 0; i < patch_nb; i++) {
                    // ***** Stage 1: Differential production of gametophytes by adults *****
                    // Calculate the mean gametophyte fitness, male and female
                    w_male_mean[i] = (gtype_freq[i][0] * w_gphyte_male[i][0]) + (gtype_freq[i][1] * w_gphyte_male[i][1]) + (gtype_freq[i][2] * w_gphyte_male[i][2]);
                    w_female_mean[i] = (gtype_freq[i][0] * w_gphyte_female[i][0]) + (gtype_freq[i][1] * w_gphyte_female[i][1]) + (gtype_freq[i][2] * w_gphyte_female[i][2]);
                    // Frequency of A after gametophytic selection
                    gametophyte_male[i] = (gtype_freq[i][0] * w_gphyte_male[i][0] + (gtype_freq[i][1] * w_gphyte_male[i][1])/2) / w_male_mean[i];
                    gametophyte_female[i] = (gtype_freq[i][0] * w_gphyte_female[i][0] + (gtype_freq[i][1] * w_gphyte_female[i][1])/2) / w_female_mean[i];
                    // ***** Stage 2: Differential production of gametes by gametophytes *****
                    // Calculate the mean gamete fitness, male and female
                    w_gphyte_male_mean[i] = gametophyte_male[i] * wA_gamete_male[i] + (1-gametophyte_male[i]) * wa_gamete_male[i];
                    w_gphyte_female_mean[i] = gametophyte_female[i] * wA_gamete_female[i] + (1-gametophyte_female[i]) * wa_gamete_female[i];


                    // Frequency of A after gametic selection
                    allele_freq_male_gamete[i] = gametophyte_male[i] * (wA_gamete_male[i])/(w_gphyte_male_mean[i]);
                    allele_freq_female_gamete[i] = gametophyte_female[i] * (wA_gamete_female[i])/(w_gphyte_female_mean[i]);

                }

                // ***** Stage 3: Male gamete dispersal *****
                // New patch i freq of A = fraction mig. j-->i  +  fraction stay in ith patch
                allele_frame[0] = (mig_rate_pollen[0] * allele_freq_male_gamete[1]) + ((1 - mig_rate_pollen[0]) * allele_freq_male_gamete[0]);
                allele_frame[1] = (mig_rate_pollen[1] * allele_freq_male_gamete[0]) + ((1 - mig_rate_pollen[1]) * allele_freq_male_gamete[1]);

                // update allele frequencies
                allele_freq_male_gamete[0]=allele_frame[0];
                allele_freq_male_gamete[1]=allele_frame[1];


                /* ================================== Reproduction ================================== */
                // ***** Stage 4: Mating, (1-S) fraction of the population outcrosses and S fraction is selfing *****
                for (i = 0; i < patch_nb; i++) {

                    // Calculate the contribution parameters
                    w_A_F_ = (wA_gamete_female[i])/(w_gphyte_female_mean[i]);
                    w_a_F_ = (wa_gamete_female[i])/(w_gphyte_female_mean[i]);

                    w_A_m_ = wA_gamete_male[i]/(wA_gamete_male[i] + wa_gamete_male[i]);
                    w_a_m_ = wa_gamete_male[i]/(wA_gamete_male[i] + wa_gamete_male[i]);

                    w_AA_F_ = (w_gphyte_female[i][0])/(w_female_mean[i]);
                    w_Aa_F_ = (w_gphyte_female[i][1])/(w_female_mean[i]);

                    offspring_self[0] = w_A_F_ * ( gtype_freq[i][0] * w_AA_F_ + w_Aa_F_ * w_A_m_ * gtype_freq[i][1]/2.0 );
                    offspring_self[1] = gtype_freq[i][1]/2.0 * w_Aa_F_ * ( w_A_F_ * w_a_m_ + w_a_F_ * w_A_m_ );
                    offspring_self[2] = 1 - offspring_self[0] - offspring_self[1];

                    offspring_outcross[0] = (allele_freq_male_gamete[i]*allele_freq_female_gamete[i]);
                    offspring_outcross[1] = (allele_freq_male_gamete[i]*(1-allele_freq_female_gamete[i])+allele_freq_female_gamete[i]*(1-allele_freq_male_gamete[i]));
                    offspring_outcross[2] = ((1-allele_freq_male_gamete[i])*(1-allele_freq_female_gamete[i]));

                    for (k=0; k < genotype_nb; k++) gtype_freq[i][k] =(1-sigma)*offspring_outcross[k] + sigma*offspring_self[k];
                    gsl_ran_multinomial(gBaseRand, genotype_nb, patch_size[i], (const double*)gtype_freq[i], offspring_full);
                    for (k=0; k < genotype_nb; k++) gtype_freq[i][k] = (double)offspring_full[k]/(double)patch_size[i];

                }

                /* ================================== Offspring dispersal ================================== */
                // ***** Stage 5: Formed offspring disperses *****
                for (j = 0; j < genotype_nb; j++) {

                    //      new patch 0 = fraction mig. from patch 1  +  fraction stay in patch 0
                    transit_frame[0][j] = (mig_rate_seed[0] * gtype_freq[1][j] + (1 - mig_rate_seed[0]) * gtype_freq[0][j]);

                    //      new patch 1 = fraction mig. from patch 0  +  fraction stay in patch 1
                    transit_frame[1][j] = (mig_rate_seed[1] * gtype_freq[0][j] + (1 - mig_rate_seed[1]) * gtype_freq[1][j]);

                }

                for (i=0;i<patch_nb;i++) {
                    for (j=0;j<genotype_nb;j++) gtype_freq[i][j]=transit_frame[i][j];
                }


                /* ================================== Survival ================================== */
                // ***** Stage 6: Differential survival of offspring to adulthood *****
                for (i=0; i<patch_nb; i++) {
                    mean_w = w[i][0]*gtype_freq[i][0]+w[i][1]*gtype_freq[i][1]+w[i][2]*gtype_freq[i][2];            // w_bar = fAA*WAA + fAa*WAa + faa*Waa


                    for (j=0; j<genotype_nb; j++) gtype_freq[i][j]=(w[i][j]*gtype_freq[i][j])/(mean_w);             // survival_prob(..)=f..*(W../w_bar)
                }


                /* ================================== Extinction/Fixation conditions ============ */
                // if mutant number in the good patch exceeds 1/s, count as successful invasion
                // run the simulation until allele fixes or goes extinct --> record sojourn time
                invader_nb = patch_size[0]*(2*gtype_freq[0][0] + (gtype_freq[0][1]));

                if (((gtype_freq[0][0] + gtype_freq[0][1]/2.0)+(gtype_freq[1][0] + gtype_freq[1][1]/2.0)==0) || (t>time_threshold))
                {
                    exit_=true;
                } else {
                    if ( invader_nb>invasion_thresh )
                    {
                        exit_=true;
                        time_invasion += t;
                        invasion_count++;
                    }
                }
                t++;
        }
    }

    // Print: input pars, invasion probability, mean invasion time
    printf("%lf %lf %lf %lf %lf %lf %lf %lf %lf %lf %lf %lf %lf %lf %lf %lf %lf %lf %lf %lf %lf %lf %lf %lf %lf %lf %lf %lf %lf %lf %lf %u %u %u %u %u %u %lf %lf\n",
            w[0][0],
            w[0][1],
            w[0][2],
            w[1][0],
            w[1][1],
            w[1][2],

            w_gphyte_female[0][0],
            w_gphyte_female[0][1],
            w_gphyte_female[0][2],
            w_gphyte_female[1][0],
            w_gphyte_female[1][1],
            w_gphyte_female[1][2],

            w_gphyte_male[0][0],
            w_gphyte_male[0][1],
            w_gphyte_male[0][2],
            w_gphyte_male[1][0],
            w_gphyte_male[1][1],
            w_gphyte_male[1][2],

            wA_gamete_female[0],
            wa_gamete_female[0],
            wA_gamete_male[0],
            wa_gamete_male[0],

            wA_gamete_female[1],
            wa_gamete_female[1],
            wA_gamete_male[1],
            wa_gamete_male[1],

            mig_rate_pollen[0],
            mig_rate_pollen[1],
            mig_rate_seed[0],
            mig_rate_seed[1],

            sigma,
            patch_size[0],
            patch_size[1],
            mutant_init[0],
            mutant_init[1],

            run_nb,
            randSeed,
            (double)invasion_count/run_nb,
            (double)time_invasion/invasion_count);
return 0;
}
```
