## Supplementary material for "Establishment of local adaptation in partly self-fertilizing populations": Scripts for running the simulator and processing the raw data, together with the complete output necessary for plotting using Mathematica notebook.: compare_simulations_gamete.R

```
remove(list=ls())
setwd("~/Desktop/incoming_projects/mating_system/simulation/selfing_sim/processed_data_gamete_dispersal/")
library(dplyr)
library(ggplot2)
library(scales)

#================== Viability selection ==================
# Start in good patch
df <-read.table(file = './good_via_gamete.out',header = FALSE, stringsAsFactors = FALSE)
colnames(df) <- c("V1_AA","V1_Aa","V1_aa","V2_AA","V2_Aa","V2_aa",
                  "WF1_AA","WF1_Aa","WF1_aa","WF2_AA","WF2_Aa","WF2_aa",
                  "WM1_AA","WM1_Aa","WM1_aa","WM2_AA","WM2_Aa","WM2_aa",
                  "wF1_A","wF1_a","wM1_A","wM1_a",
                  "wF2_A","wF2_a","wM2_A","wM2_a",
                  "mP1","mP2","mS1","mS2","sigma",
                  "N1","N2","mu1","mu2","run_nb","rndSeed",
                  "P_est","P_time")
# When considering viability selection, we can only focus on subset of all columns
df <- df %>% select(V1_AA,V1_Aa,V1_aa,V2_AA,V2_Aa,V2_aa,sigma,mP1,mP2,mS1,mS2,P_est) %>% na.omit

# Parameters used when running the simulation; Change if required
codom_vector <- c(1.01, 1.005, 1, 0.99, 0.995, 1)
dom_vector <- c(1.01, 1.0075, 1, 0.99, 0.9925, 1)
rec_vector <- c(1.01, 1.0025, 1, 0.99, 0.9975, 1)
asym_sel <- c(1.0125, 1.01, 1, 0.99, 0.995, 1)

# Make separate data frames holding scenarios with different dominances
df_codom <- df[unlist(apply(df[,c(1:6)], 1, function(x) all(x==codom_vector))),]    # Simulations with codominance
df_dom <- df[unlist(apply(df[,c(1:6)], 1, function(x) all(x==dom_vector))),]        # Simulations with dominance
df_rec <- df[unlist(apply(df[,c(1:6)], 1, function(x) all(x==rec_vector))),]        # Simulations with recessivity
df_asym_sel <- df[unlist(apply(df[,c(1:6)], 1, function(x) all(x==asym_sel))),]     # Simulations with asymmetric selection
df_asym_mig <- df_codom[df_codom$mP2!=df_codom$mP1,]                                # Simulations with asymmetric migration
# Imporant note: Because asymmetric migration has been executed for a codominant case
#                when wanting to plot codominance and symmetric selection, one has to
#                subset the whole of results with codominance for those cases with symmetrical pollen dispersal
df_codom <- df_codom[df_codom$mP2==df_codom$mP1,]

# Plotting part
df_codom_plot <- df_codom %>% group_by(sigma,mP1) %>%  
  summarise(mean_Pest=mean(P_est), sd_Pest=sd(P_est), .groups='drop')
write.csv(df_codom_plot, './good_via_codom_processed.csv', row.names = FALSE)
ggplot(df_codom_plot, aes(x=mP1, y=mean_Pest, color=sigma)) + scale_x_continuous(trans = log10_trans()) + geom_point()

df_dom_plot <- df_dom %>% group_by(sigma,mP1) %>%  
  summarise(mean_Pest=mean(P_est), sd_Pest=sd(P_est), .groups='drop')
write.csv(df_dom_plot, './good_via_dom_processed.csv', row.names = FALSE)
ggplot(df_dom_plot, aes(x=mP1, y=mean_Pest, color=sigma)) + scale_x_continuous(trans = log10_trans()) + geom_point()

df_rec_plot <- df_rec %>% group_by(sigma,mP1) %>%  
  summarise(mean_Pest=mean(P_est), sd_Pest=sd(P_est), .groups='drop')
write.csv(df_rec_plot, './good_via_rec_processed.csv', row.names = FALSE)
ggplot(df_rec_plot, aes(x=mP1, y=mean_Pest, color=sigma)) + scale_x_continuous(trans = log10_trans()) + geom_point()

df_asym_sel_plot <- df_asym_sel %>% group_by(sigma,mP1) %>%  
  summarise(mean_Pest=mean(P_est), sd_Pest=sd(P_est), .groups='drop')
write.csv(df_asym_sel_plot, './good_via_asym_sel_processed.csv', row.names = FALSE)
ggplot(df_asym_sel_plot, aes(x=mP1, y=mean_Pest, color=sigma)) + scale_x_continuous(trans = log10_trans()) + geom_point()

df_asym_mig_plot <- df_asym_mig %>% group_by(sigma,mP1) %>%  
  summarise(mean_Pest=mean(P_est), sd_Pest=sd(P_est), .groups='drop')
write.csv(df_asym_mig_plot, './good_via_asym_mig_processed.csv', row.names = FALSE)
ggplot(df_asym_mig_plot, aes(x=mP1, y=mean_Pest, color=sigma)) + scale_x_continuous(trans = log10_trans()) + geom_point()


# Start in bad patch
df <-read.table(file = './bad_via_gamete.out',header = FALSE, stringsAsFactors = FALSE)
colnames(df) <- c("V1_AA","V1_Aa","V1_aa","V2_AA","V2_Aa","V2_aa",
                  "WF1_AA","WF1_Aa","WF1_aa","WF2_AA","WF2_Aa","WF2_aa",
                  "WM1_AA","WM1_Aa","WM1_aa","WM2_AA","WM2_Aa","WM2_aa",
                  "wF1_A","wF1_a","wM1_A","wM1_a",
                  "wF2_A","wF2_a","wM2_A","wM2_a",
                  "mP1","mP2","mS1","mS2","sigma",
                  "N1","N2","mu1","mu2","run_nb","rndSeed",
                  "P_est","P_time")
df <- df %>% select(V1_AA,V1_Aa,V1_aa,V2_AA,V2_Aa,V2_aa,sigma,mP1,mP2,mS1,mS2,P_est) %>% na.omit

codom_vector <- c(1.01, 1.005, 1, 0.99, 0.995, 1)
dom_vector <- c(1.01, 1.0075, 1, 0.99, 0.9925, 1)
rec_vector <- c(1.01, 1.0025, 1, 0.99, 0.9975, 1)
asym_sel <- c(1.0125, 1.01, 1, 0.99, 0.995, 1)
df_codom <- df[unlist(apply(df[,c(1:6)], 1, function(x) all(x==codom_vector))),]

df_dom <- df[unlist(apply(df[,c(1:6)], 1, function(x) all(x==dom_vector))),]
df_rec <- df[unlist(apply(df[,c(1:6)], 1, function(x) all(x==rec_vector))),]
df_asym_sel <- df[unlist(apply(df[,c(1:6)], 1, function(x) all(x==asym_sel))),]
df_asym_mig <- df_codom[df_codom$mP1!=df_codom$mP2,]
df_codom <- df_codom[df_codom$mP1==df_codom$mP2,]

df_codom_plot <- df_codom %>% group_by(sigma,mP1) %>%  
  summarise(mean_Pest=mean(P_est), sd_Pest=sd(P_est), .groups='drop')
write.csv(df_codom_plot, './bad_via_codom_processed.csv', row.names = FALSE)
ggplot(df_codom_plot, aes(x=mP1, y=mean_Pest, color=sigma)) + scale_x_continuous(trans = log10_trans()) + geom_point()

df_dom_plot <- df_dom %>% group_by(sigma,mP1) %>%  
  summarise(mean_Pest=mean(P_est), sd_Pest=sd(P_est), .groups='drop')
write.csv(df_dom_plot, './bad_via_dom_processed.csv', row.names = FALSE)
ggplot(df_dom_plot, aes(x=mP1, y=mean_Pest, color=sigma)) + scale_x_continuous(trans = log10_trans()) + geom_point()

df_rec_plot <- df_rec %>% group_by(sigma,mP1) %>%  
  summarise(mean_Pest=mean(P_est), sd_Pest=sd(P_est), .groups='drop')
write.csv(df_rec_plot, './bad_via_rec_processed.csv', row.names = FALSE)
ggplot(df_rec_plot, aes(x=mP1, y=mean_Pest, color=sigma)) + scale_x_continuous(trans = log10_trans()) + geom_point()

df_asym_sel_plot <- df_asym_sel %>% group_by(sigma,mP1) %>%  
  summarise(mean_Pest=mean(P_est), sd_Pest=sd(P_est), .groups='drop')
write.csv(df_asym_sel_plot, './bad_via_asym_sel_processed.csv', row.names = FALSE)
ggplot(df_asym_sel_plot, aes(x=mP1, y=mean_Pest, color=sigma)) + scale_x_continuous(trans = log10_trans()) + geom_point()

df_asym_mig_plot <- df_asym_mig %>% group_by(sigma,mP1) %>%  
  summarise(mean_Pest=mean(P_est), sd_Pest=sd(P_est), .groups='drop')
write.csv(df_asym_mig_plot, './bad_via_asym_mig_processed.csv', row.names = FALSE)
ggplot(df_asym_mig_plot, aes(x=mP1, y=mean_Pest, color=sigma)) + scale_x_continuous(trans = log10_trans()) + geom_point()


#================== Female fitness component selection ==================
# Start in good patch
df <-read.table(file = './good_female_gamete.out',header = FALSE, stringsAsFactors = FALSE)
colnames(df) <- c("V1_AA","V1_Aa","V1_aa","V2_AA","V2_Aa","V2_aa",
                  "WF1_AA","WF1_Aa","WF1_aa","WF2_AA","WF2_Aa","WF2_aa",
                  "WM1_AA","WM1_Aa","WM1_aa","WM2_AA","WM2_Aa","WM2_aa",
                  "wF1_A","wF1_a","wM1_A","wM1_a",
                  "wF2_A","wF2_a","wM2_A","wM2_a",
                  "mP1","mP2","mS1","mS2","sigma",
                  "N1","N2","mu1","mu2","run_nb","rndSeed",
                  "P_est","P_time")
df <- df %>% select(WF1_AA,WF1_Aa,WF1_aa,WF2_AA,WF2_Aa,WF2_aa,sigma,mP1,mP2,mS1,mS2,P_est) %>% na.omit

codom_vector <- c(1.01, 1.005, 1, 0.99, 0.995, 1)
dom_vector <- c(1.01, 1.0075, 1, 0.99, 0.9925, 1)
rec_vector <- c(1.01, 1.0025, 1, 0.99, 0.9975, 1)
asym_sel <- c(1.0125, 1.01, 1, 0.99, 0.995, 1)
df_codom <- df[unlist(apply(df[,c(1:6)], 1, function(x) all(x==codom_vector))),]

df_dom <- df[unlist(apply(df[,c(1:6)], 1, function(x) all(x==dom_vector))),]
df_rec <- df[unlist(apply(df[,c(1:6)], 1, function(x) all(x==rec_vector))),]
df_asym_sel <- df[unlist(apply(df[,c(1:6)], 1, function(x) all(x==asym_sel))),]
df_asym_mig <- df_codom[df_codom$mP2!=df_codom$mP1,]
df_codom <- df_codom[df_codom$mP2==df_codom$mP1,]

df_codom_plot <- df_codom %>% group_by(sigma,mP1) %>%  
  summarise(mean_Pest=mean(P_est), sd_Pest=sd(P_est), .groups='drop')
write.csv(df_codom_plot, './good_female_codom_processed.csv', row.names = FALSE)
ggplot(df_codom_plot, aes(x=mP1, y=mean_Pest, color=sigma)) + scale_x_continuous(trans = log10_trans()) + geom_point()

df_dom_plot <- df_dom %>% group_by(sigma,mP1) %>%  
  summarise(mean_Pest=mean(P_est), sd_Pest=sd(P_est), .groups='drop')
write.csv(df_dom_plot, './good_female_dom_processed.csv', row.names = FALSE)
ggplot(df_dom_plot, aes(x=mP1, y=mean_Pest, color=sigma)) + scale_x_continuous(trans = log10_trans()) + geom_point()

df_rec_plot <- df_rec %>% group_by(sigma,mP1) %>%  
  summarise(mean_Pest=mean(P_est), sd_Pest=sd(P_est), .groups='drop')
write.csv(df_rec_plot, './good_female_rec_processed.csv', row.names = FALSE)
ggplot(df_rec_plot, aes(x=mP1, y=mean_Pest, color=sigma)) + scale_x_continuous(trans = log10_trans()) + geom_point()

df_asym_sel_plot <- df_asym_sel %>% group_by(sigma,mP1) %>%  
  summarise(mean_Pest=mean(P_est), sd_Pest=sd(P_est), .groups='drop')
write.csv(df_asym_sel_plot, './good_female_asym_sel_processed.csv', row.names = FALSE)
ggplot(df_asym_sel_plot, aes(x=mP1, y=mean_Pest, color=sigma)) + scale_x_continuous(trans = log10_trans()) + geom_point()

df_asym_mig_plot <- df_asym_mig %>% group_by(sigma,mP1) %>%  
  summarise(mean_Pest=mean(P_est), sd_Pest=sd(P_est), .groups='drop')
write.csv(df_asym_mig_plot, './good_female_asym_mig_processed.csv', row.names = FALSE)
ggplot(df_asym_mig_plot, aes(x=mP1, y=mean_Pest, color=sigma)) + scale_x_continuous(trans = log10_trans()) + geom_point()


# Start in bad patch
df <-read.table(file = './bad_female_gamete.out',header = FALSE, stringsAsFactors = FALSE)
colnames(df) <- c("V1_AA","V1_Aa","V1_aa","V2_AA","V2_Aa","V2_aa",
                  "WF1_AA","WF1_Aa","WF1_aa","WF2_AA","WF2_Aa","WF2_aa",
                  "WM1_AA","WM1_Aa","WM1_aa","WM2_AA","WM2_Aa","WM2_aa",
                  "wF1_A","wF1_a","wM1_A","wM1_a",
                  "wF2_A","wF2_a","wM2_A","wM2_a",
                  "mP1","mP2","mS1","mS2","sigma",
                  "N1","N2","mu1","mu2","run_nb","rndSeed",
                  "P_est","P_time")
df <- df %>% select(WF1_AA,WF1_Aa,WF1_aa,WF2_AA,WF2_Aa,WF2_aa,sigma,mP1,mP2,mS1,mS2,P_est) %>% na.omit

codom_vector <- c(1.01, 1.005, 1, 0.99, 0.995, 1)
dom_vector <- c(1.01, 1.0075, 1, 0.99, 0.9925, 1)
rec_vector <- c(1.01, 1.0025, 1, 0.99, 0.9975, 1)
asym_sel <- c(1.0125, 1.01, 1, 0.99, 0.995, 1)
df_codom <- df[unlist(apply(df[,c(1:6)], 1, function(x) all(x==codom_vector))),]

df_dom <- df[unlist(apply(df[,c(1:6)], 1, function(x) all(x==dom_vector))),]
df_rec <- df[unlist(apply(df[,c(1:6)], 1, function(x) all(x==rec_vector))),]
df_asym_sel <- df[unlist(apply(df[,c(1:6)], 1, function(x) all(x==asym_sel))),]
df_asym_mig <- df_codom[df_codom$mP2!=df_codom$mP1,]
df_codom <- df_codom[df_codom$mP2==df_codom$mP1,]

df_codom_plot <- df_codom %>% group_by(sigma,mP1) %>%  
  summarise(mean_Pest=mean(P_est), sd_Pest=sd(P_est), .groups='drop')
write.csv(df_codom_plot, './bad_female_codom_processed.csv', row.names = FALSE)
ggplot(df_codom_plot, aes(x=mP1, y=mean_Pest, color=sigma)) + scale_x_continuous(trans = log10_trans()) + geom_point()

df_dom_plot <- df_dom %>% group_by(sigma,mP1) %>%  
  summarise(mean_Pest=mean(P_est), sd_Pest=sd(P_est), .groups='drop')
write.csv(df_dom_plot, './bad_female_dom_processed.csv', row.names = FALSE)
ggplot(df_dom_plot, aes(x=mP1, y=mean_Pest, color=sigma)) + scale_x_continuous(trans = log10_trans()) + geom_point()

df_rec_plot <- df_rec %>% group_by(sigma,mP1) %>%  
  summarise(mean_Pest=mean(P_est), sd_Pest=sd(P_est), .groups='drop')
write.csv(df_rec_plot, './bad_female_rec_processed.csv', row.names = FALSE)
ggplot(df_rec_plot, aes(x=mP1, y=mean_Pest, color=sigma)) + scale_x_continuous(trans = log10_trans()) + geom_point()

df_asym_sel_plot <- df_asym_sel %>% group_by(sigma,mP1) %>%  
  summarise(mean_Pest=mean(P_est), sd_Pest=sd(P_est), .groups='drop')
write.csv(df_asym_sel_plot, './bad_female_asym_sel_processed.csv', row.names = FALSE)
ggplot(df_asym_sel_plot, aes(x=mP1, y=mean_Pest, color=sigma)) + scale_x_continuous(trans = log10_trans()) + geom_point()

df_asym_mig_plot <- df_asym_mig %>% group_by(sigma,mP1) %>%  
  summarise(mean_Pest=mean(P_est), sd_Pest=sd(P_est), .groups='drop')
write.csv(df_asym_mig_plot, './bad_female_asym_mig_processed.csv', row.names = FALSE)
ggplot(df_asym_mig_plot, aes(x=mP1, y=mean_Pest, color=sigma)) + scale_x_continuous(trans = log10_trans()) + geom_point()


#================== Male fitness component selection ==================
# Start in good patch
df <-read.table(file = './good_male_gamete.out',header = FALSE, stringsAsFactors = FALSE)
colnames(df) <- c("V1_AA","V1_Aa","V1_aa","V2_AA","V2_Aa","V2_aa",
                  "WF1_AA","WF1_Aa","WF1_aa","WF2_AA","WF2_Aa","WF2_aa",
                  "WM1_AA","WM1_Aa","WM1_aa","WM2_AA","WM2_Aa","WM2_aa",
                  "wF1_A","wF1_a","wM1_A","wM1_a",
                  "wF2_A","wF2_a","wM2_A","wM2_a",
                  "mP1","mP2","mS1","mS2","sigma",
                  "N1","N2","mu1","mu2","run_nb","rndSeed",
                  "P_est","P_time")
df <- df %>% select(WM1_AA,WM1_Aa,WM1_aa,WM2_AA,WM2_Aa,WM2_aa,sigma,mP1,mP2,mS1,mS2,P_est) %>% na.omit

codom_vector <- c(1.01, 1.005, 1, 0.99, 0.995, 1)
dom_vector <- c(1.01, 1.0075, 1, 0.99, 0.9925, 1)
rec_vector <- c(1.01, 1.0025, 1, 0.99, 0.9975, 1)
asym_sel <- c(1.0125, 1.01, 1, 0.99, 0.995, 1)
df_codom <- df[unlist(apply(df[,c(1:6)], 1, function(x) all(x==codom_vector))),]

df_dom <- df[unlist(apply(df[,c(1:6)], 1, function(x) all(x==dom_vector))),]
df_rec <- df[unlist(apply(df[,c(1:6)], 1, function(x) all(x==rec_vector))),]
df_asym_sel <- df[unlist(apply(df[,c(1:6)], 1, function(x) all(x==asym_sel))),]
df_asym_mig <- df_codom[df_codom$mP2!=df_codom$mP1,]
df_codom <- df_codom[df_codom$mP2==df_codom$mP1,]

df_codom_plot <- df_codom %>% group_by(sigma,mP1) %>%  
  summarise(mean_Pest=mean(P_est), sd_Pest=sd(P_est), .groups='drop')
write.csv(df_codom_plot, './good_male_codom_processed.csv', row.names = FALSE)
ggplot(df_codom_plot, aes(x=mP1, y=mean_Pest, color=sigma)) + scale_x_continuous(trans = log10_trans()) + geom_point()

df_dom_plot <- df_dom %>% group_by(sigma,mP1) %>%  
  summarise(mean_Pest=mean(P_est), sd_Pest=sd(P_est), .groups='drop')
write.csv(df_dom_plot, './good_male_dom_processed.csv', row.names = FALSE)
ggplot(df_dom_plot, aes(x=mP1, y=mean_Pest, color=sigma)) + scale_x_continuous(trans = log10_trans()) + geom_point()

df_rec_plot <- df_rec %>% group_by(sigma,mP1) %>%  
  summarise(mean_Pest=mean(P_est), sd_Pest=sd(P_est), .groups='drop')
write.csv(df_rec_plot, './good_male_rec_processed.csv', row.names = FALSE)
ggplot(df_rec_plot, aes(x=mP1, y=mean_Pest, color=sigma)) + scale_x_continuous(trans = log10_trans()) + geom_point()

df_asym_sel_plot <- df_asym_sel %>% group_by(sigma,mP1) %>%  
  summarise(mean_Pest=mean(P_est), sd_Pest=sd(P_est), .groups='drop')
write.csv(df_asym_sel_plot, './good_male_asym_sel_processed.csv', row.names = FALSE)
ggplot(df_asym_sel_plot, aes(x=mP1, y=mean_Pest, color=sigma)) + scale_x_continuous(trans = log10_trans()) + geom_point()

df_asym_mig_plot <- df_asym_mig %>% group_by(sigma,mP1) %>%  
  summarise(mean_Pest=mean(P_est), sd_Pest=sd(P_est), .groups='drop')
write.csv(df_asym_mig_plot, './good_male_asym_mig_processed.csv', row.names = FALSE)
ggplot(df_asym_mig_plot, aes(x=mP1, y=mean_Pest, color=sigma)) + scale_x_continuous(trans = log10_trans()) + geom_point()


# Start in bad patch
df <-read.table(file = './bad_male_gamete.out',header = FALSE, stringsAsFactors = FALSE)
colnames(df) <- c("V1_AA","V1_Aa","V1_aa","V2_AA","V2_Aa","V2_aa",
                  "WF1_AA","WF1_Aa","WF1_aa","WF2_AA","WF2_Aa","WF2_aa",
                  "WM1_AA","WM1_Aa","WM1_aa","WM2_AA","WM2_Aa","WM2_aa",
                  "wF1_A","wF1_a","wM1_A","wM1_a",
                  "wF2_A","wF2_a","wM2_A","wM2_a",
                  "mP1","mP2","mS1","mS2","sigma",
                  "N1","N2","mu1","mu2","run_nb","rndSeed",
                  "P_est","P_time")
df <- df %>% select(WM1_AA,WM1_Aa,WM1_aa,WM2_AA,WM2_Aa,WM2_aa,sigma,mP1,mP2,mS1,mS2,P_est) %>% na.omit

codom_vector <- c(1.01, 1.005, 1, 0.99, 0.995, 1)
dom_vector <- c(1.01, 1.0075, 1, 0.99, 0.9925, 1)
rec_vector <- c(1.01, 1.0025, 1, 0.99, 0.9975, 1)
asym_sel <- c(1.0125, 1.01, 1, 0.99, 0.995, 1)
df_codom <- df[unlist(apply(df[,c(1:6)], 1, function(x) all(x==codom_vector))),]

df_dom <- df[unlist(apply(df[,c(1:6)], 1, function(x) all(x==dom_vector))),]
df_rec <- df[unlist(apply(df[,c(1:6)], 1, function(x) all(x==rec_vector))),]
df_asym_sel <- df[unlist(apply(df[,c(1:6)], 1, function(x) all(x==asym_sel))),]
df_asym_mig <- df_codom[df_codom$mP2!=df_codom$mP1,]
df_codom <- df_codom[df_codom$mP2==df_codom$mP1,]

df_codom_plot <- df_codom %>% group_by(sigma,mP1) %>%  
  summarise(mean_Pest=mean(P_est), sd_Pest=sd(P_est), .groups='drop')
write.csv(df_codom_plot, './bad_male_codom_processed.csv', row.names = FALSE)
ggplot(df_codom_plot, aes(x=mP1, y=mean_Pest, color=sigma)) + scale_x_continuous(trans = log10_trans()) + geom_point()

df_dom_plot <- df_dom %>% group_by(sigma,mP1) %>%  
  summarise(mean_Pest=mean(P_est), sd_Pest=sd(P_est), .groups='drop')
write.csv(df_dom_plot, './bad_male_dom_processed.csv', row.names = FALSE)
ggplot(df_dom_plot, aes(x=mP1, y=mean_Pest, color=sigma)) + scale_x_continuous(trans = log10_trans()) + geom_point()

df_rec_plot <- df_rec %>% group_by(sigma,mP1) %>%  
  summarise(mean_Pest=mean(P_est), sd_Pest=sd(P_est), .groups='drop')
write.csv(df_rec_plot, './bad_male_rec_processed.csv', row.names = FALSE)
ggplot(df_rec_plot, aes(x=mP1, y=mean_Pest, color=sigma)) + scale_x_continuous(trans = log10_trans()) + geom_point()

df_asym_sel_plot <- df_asym_sel %>% group_by(sigma,mP1) %>%  
  summarise(mean_Pest=mean(P_est), sd_Pest=sd(P_est), .groups='drop')
write.csv(df_asym_sel_plot, './bad_male_asym_sel_processed.csv', row.names = FALSE)
ggplot(df_asym_sel_plot, aes(x=mP1, y=mean_Pest, color=sigma)) + scale_x_continuous(trans = log10_trans()) + geom_point()

df_asym_mig_plot <- df_asym_mig %>% group_by(sigma,mP1) %>%  
  summarise(mean_Pest=mean(P_est), sd_Pest=sd(P_est), .groups='drop')
write.csv(df_asym_mig_plot, './bad_male_asym_mig_processed.csv', row.names = FALSE)
ggplot(df_asym_mig_plot, aes(x=mP1, y=mean_Pest, color=sigma)) + scale_x_continuous(trans = log10_trans()) + geom_point()
```
