## Supplementary material for "Establishment of local adaptation in partly self-fertilizing populations": Scripts for running the simulator and processing the raw data, together with the complete output necessary for plotting using Mathematica notebook.: mating_system_offspring.sh

```
#!/bin/bash

# +++++++++++++++++++++++++++++++++++++++++++++++++++++++++++
# ++++++++++++++++++++++++ Comments +++++++++++++++++++++++++
# +++++++++++++++++++++++++++++++++++++++++++++++++++++++++++

# Usage: selfing_sim [w1AA][w1Aa][w1aa][w2AA][w2Aa][w2aa][m12pollen][m21pollen][m12seed][m21seed][sigma][patch1_size][patch2_size][mut_init1][mut_init2][run_nb][rand_seed]
#
# When running on personal computer, add gsl to PATH:
LD_LIBRARY_PATH=/home/bogi/gsl/lib
export LD_LIBRARY_PATH

# If running on cluster, don't forget to load gsl module
#module load gsl/2.4 

# +++++++++++++++++++++++++++++++++++++++++++++++++++++++++++
# ++++++++++++++++++ The overall philosophy +++++++++++++++++
# +++++++++++++++++++++++++++++++++++++++++++++++++++++++++++

# 1. Generate as many PRN as there are simulation runs; Use device files as PRNG
# 2. Execute simulations via 'parallel' and pipe all PRNs; This yields data for one set of parameters (e.g., migration rate)
# 3. Set new input parameters and goto step 1

generate_rnd() {
	>$rnd_src                                                         
    	for ((k = 1 ; k <= sim_runs ; k++))                                         
    	do                                                                          
        	cat /dev/urandom | od -vAn -N4 -tu4 >> $rnd_src               
    	done
}


# +++++++++++++++++++++++++++++++++++++++++++++++++++++++++++
# ++++++++++++++++++++++++ Parameters +++++++++++++++++++++++
# +++++++++++++++++++++++++++++++++++++++++++++++++++++++++++

alpha=1.25		# Multiplies migration rate (patch 1 --> patch 2) when asymmetrical migration is studied
beta=1.0125		# Denotes selection coefficient (patch 1) when asymmetrical selection is studied
sim_runs=1000		# For each combination of parameters, run 1000 simulations
# Output is grouped by the patch where mutant was injected (good vs. bad), and selection mode (viability, female fecundity, and male fecundity, respectively)
# We will process the raw output using 'compare_simulations.R' and finally plot it from Mathematica notebook
good_start_via="./good_via_offspring.out"
bad_start_via="./bad_via_offspring.out"
good_start_female="./good_female_offspring.out"
bad_start_female="./bad_female_offspring.out"
good_start_male="./good_male_offspring.out"
bad_start_male="./bad_male_offspring.out"
rnd_src="rand_seed.out"
>$rnd_src
seed_pars=( 0.0002 0.000427592 0.000914176 0.00195447 0.00417859 0.00893367 0.0190999 0.0408348 0.0873032 0.186651 )		# Seed migration rates
selfing_pars=( 0.00 0.50 1.00 )													# Three selfing rates examined


# +++++++++++++++++++++++++++++++++++++++++++++++++++++++++++
# ++++++++++++++++++++++++ Bulk run +++++++++++++++++++++++++
# +++++++++++++++++++++++++++++++++++++++++++++++++++++++++++

# Execution order:
# 1) Symmetric selection, symmetric migration, codominant
# 2) Symmetric selection, symmetric migration, dominant
# 3) Symmetric selection, symmetric migration, recessive
# 4) Symmetric selection, asymmetric migration, codominant
# 5) Asymmetric selection, symmetric migration, codominant
# Selection on viability
>$good_start_via
>$bad_start_via
for i in "${selfing_pars[@]}"
do
	for j in "${seed_pars[@]}"
	do
	generate_rnd
	cat $rnd_src | parallel "./selfing_sim 1.01 1.005 1 0.99 0.995 1 1 1 1 1 1 1 1 1 1 1 1 1 1 1 1 1 1 1 1 1 0 0 $j $j $i 10000 10000 1 0 10000 {}" >> $good_start_via
	generate_rnd
	cat $rnd_src | parallel "./selfing_sim 1.01 1.0075 1 0.99 0.9925 1 1 1 1 1 1 1 1 1 1 1 1 1 1 1 1 1 1 1 1 1 0 0 $j $j $i 10000 10000 1 0 10000 {}" >> $good_start_via
	generate_rnd	
	cat $rnd_src | parallel "./selfing_sim 1.01 1.0025 1 0.99 0.9975 1 1 1 1 1 1 1 1 1 1 1 1 1 1 1 1 1 1 1 1 1 0 0 $j $j $i 10000 10000 1 0 10000 {}" >> $good_start_via
	generate_rnd	
	cat $rnd_src | parallel "./selfing_sim 1.01 1.005 1 0.99 0.995 1 1 1 1 1 1 1 1 1 1 1 1 1 1 1 1 1 1 1 1 1 0 0 $(echo $alpha*$j | bc -l) $j $i 10000 10000 1 0 10000 {}" >> $good_start_via
	generate_rnd
	cat $rnd_src | parallel "./selfing_sim $beta 1.01 1 0.99 0.995 1 1 1 1 1 1 1 1 1 1 1 1 1 1 1 1 1 1 1 1 1 0 0 $j $j $i 10000 10000 1 0 10000 {}" >> $good_start_via
	done
done

for i in "${selfing_pars[@]}"
do
	for j in "${seed_pars[@]}"
	do
	generate_rnd
	cat $rnd_src | parallel "./selfing_sim 1.01 1.005 1 0.99 0.995 1 1 1 1 1 1 1 1 1 1 1 1 1 1 1 1 1 1 1 1 1 0 0 $j $j $i 10000 10000 0 1 10000 {}" >> $bad_start_via
	generate_rnd
	cat $rnd_src | parallel "./selfing_sim 1.01 1.0075 1 0.99 0.9925 1 1 1 1 1 1 1 1 1 1 1 1 1 1 1 1 1 1 1 1 1 0 0 $j $j $i 10000 10000 0 1 10000 {}" >> $bad_start_via
	generate_rnd	
	cat $rnd_src | parallel "./selfing_sim 1.01 1.0025 1 0.99 0.9975 1 1 1 1 1 1 1 1 1 1 1 1 1 1 1 1 1 1 1 1 1 0 0 $j $j $i 10000 10000 0 1 10000 {}" >> $bad_start_via
	generate_rnd	
	cat $rnd_src | parallel "./selfing_sim 1.01 1.005 1 0.99 0.995 1 1 1 1 1 1 1 1 1 1 1 1 1 1 1 1 1 1 1 1 1 0 0 $(echo $alpha*$j | bc -l) $j $i 10000 10000 0 1 10000 {}" >> $bad_start_via
	generate_rnd
	cat $rnd_src | parallel "./selfing_sim $beta 1.01 1 0.99 0.995 1 1 1 1 1 1 1 1 1 1 1 1 1 1 1 1 1 1 1 1 1 0 0 $j $j $i 10000 10000 0 1 10000 {}" >> $bad_start_via
	done
done


# Selection on female component
>$good_start_female
>$bad_start_female
for i in "${selfing_pars[@]}"
do
	for j in "${seed_pars[@]}"
	do
	generate_rnd
	cat $rnd_src | parallel "./selfing_sim 1 1 1 1 1 1 1.01 1.005 1 0.99 0.995 1 1 1 1 1 1 1 1 1 1 1 1 1 1 1 0 0 $j $j $i 10000 10000 1 0 10000 {}" >> $good_start_female
	generate_rnd	
	cat $rnd_src | parallel "./selfing_sim 1 1 1 1 1 1 1.01 1.0075 1 0.99 0.9925 1 1 1 1 1 1 1 1 1 1 1 1 1 1 1 0 0 $j $j $i 10000 10000 1 0 10000 {}" >> $good_start_female
	generate_rnd
	cat $rnd_src | parallel "./selfing_sim 1 1 1 1 1 1 1.01 1.0025 1 0.99 0.9975 1 1 1 1 1 1 1 1 1 1 1 1 1 1 1 0 0 $j $j $i 10000 10000 1 0 10000 {}" >> $good_start_female
	generate_rnd
	cat $rnd_src | parallel "./selfing_sim 1 1 1 1 1 1 1.01 1.005 1 0.99 0.995 1 1 1 1 1 1 1 1 1 1 1 1 1 1 1 0 0 $(echo $alpha*$j | bc -l) $j $i 10000 10000 1 0 10000 {}" >> $good_start_female
	generate_rnd
	cat $rnd_src | parallel "./selfing_sim 1 1 1 1 1 1 $beta 1.01 1 0.99 0.995 1 1 1 1 1 1 1 1 1 1 1 1 1 1 1 0 0 $j $j $i 10000 10000 1 0 10000 {}" >> $good_start_female
	done
done

for i in "${selfing_pars[@]}"
do
	for j in "${seed_pars[@]}"
	do 
	generate_rnd
	cat $rnd_src | parallel "./selfing_sim 1 1 1 1 1 1 1.01 1.005 1 0.99 0.995 1 1 1 1 1 1 1 1 1 1 1 1 1 1 1 0 0 $j $j $i 10000 10000 0 1 10000 {}" >> $bad_start_female
	generate_rnd
	cat $rnd_src | parallel "./selfing_sim 1 1 1 1 1 1 1.01 1.0075 1 0.99 0.9925 1 1 1 1 1 1 1 1 1 1 1 1 1 1 1 0 0 $j $j $i 10000 10000 0 1 10000 {}" >> $bad_start_female
	generate_rnd
	cat $rnd_src | parallel "./selfing_sim 1 1 1 1 1 1 1.01 1.0025 1 0.99 0.9975 1 1 1 1 1 1 1 1 1 1 1 1 1 1 1 0 0 $j $j $i 10000 10000 0 1 10000 {}" >> $bad_start_female
	generate_rnd
	cat $rnd_src | parallel "./selfing_sim 1 1 1 1 1 1 1.01 1.005 1 0.99 0.995 1 1 1 1 1 1 1 1 1 1 1 1 1 1 1 0 0 $(echo $alpha*$j | bc -l) $j $i 10000 10000 0 1 10000 {}" >> $bad_start_female
	generate_rnd
	cat $rnd_src | parallel "./selfing_sim 1 1 1 1 1 1 $beta 1.01 1 0.99 0.995 1 1 1 1 1 1 1 1 1 1 1 1 1 1 1 0 0 $j $j $i 10000 10000 0 1 10000 {}" >> $bad_start_female
	done
done


# Selection on male component
>$good_start_male
>$bad_start_male
for i in "${selfing_pars[@]}"
do
	for j in "${seed_pars[@]}"
	do
	generate_rnd
	cat $rnd_src | parallel "./selfing_sim 1 1 1 1 1 1 1 1 1 1 1 1 1.01 1.005 1 0.99 0.995 1 1 1 1 1 1 1 1 1 0 0 $j $j $i 10000 10000 1 0 10000 {}" >> $good_start_male
	generate_rnd
	cat $rnd_src | parallel "./selfing_sim 1 1 1 1 1 1 1 1 1 1 1 1 1.01 1.0075 1 0.99 0.9925 1 1 1 1 1 1 1 1 1 0 0 $j $j $i 10000 10000 1 0 10000 {}" >> $good_start_male
	generate_rnd
	cat $rnd_src | parallel "./selfing_sim 1 1 1 1 1 1 1 1 1 1 1 1 1.01 1.0025 1 0.99 0.9975 1 1 1 1 1 1 1 1 1 0 0 $j $j $i 10000 10000 1 0 10000 {}" >> $good_start_male
	generate_rnd
	cat $rnd_src | parallel "./selfing_sim 1 1 1 1 1 1 1 1 1 1 1 1 1.01 1.005 1 0.99 0.995 1 1 1 1 1 1 1 1 1 0 0 $(echo $alpha*$j | bc -l) $j $i 10000 10000 1 0 10000 {}" >> $good_start_male
	generate_rnd
	cat $rnd_src | parallel "./selfing_sim 1 1 1 1 1 1 1 1 1 1 1 1 $beta 1.01 1 0.99 0.995 1 1 1 1 1 1 1 1 1 0 0 $j $j $i 10000 10000 1 0 10000 {}" >> $good_start_male
	done
done

for i in "${selfing_pars[@]}"
do
	for j in "${seed_pars[@]}"
	do
	generate_rnd
	cat $rnd_src | parallel "./selfing_sim 1 1 1 1 1 1 1 1 1 1 1 1 1.01 1.005 1 0.99 0.995 1 1 1 1 1 1 1 1 1 0 0 $j $j $i 10000 10000 0 1 10000 {}" >> $bad_start_male
	generate_rnd
	cat $rnd_src | parallel "./selfing_sim 1 1 1 1 1 1 1 1 1 1 1 1 1.01 1.0075 1 0.99 0.9925 1 1 1 1 1 1 1 1 1 0 0 $j $j $i 10000 10000 0 1 10000 {}" >> $bad_start_male
	generate_rnd
	cat $rnd_src | parallel "./selfing_sim 1 1 1 1 1 1 1 1 1 1 1 1 1.01 1.0025 1 0.99 0.9975 1 1 1 1 1 1 1 1 1 0 0 $j $j $i 10000 10000 0 1 10000 {}" >> $bad_start_male
	generate_rnd
	cat $rnd_src | parallel "./selfing_sim 1 1 1 1 1 1 1 1 1 1 1 1 1.01 1.005 1 0.99 0.995 1 1 1 1 1 1 1 1 1 0 0 $(echo $alpha*$j | bc -l) $j $i 10000 10000 0 1 10000 {}" >> $bad_start_male
	generate_rnd
	cat $rnd_src | parallel "./selfing_sim 1 1 1 1 1 1 1 1 1 1 1 1 $beta 1.01 1 0.99 0.995 1 1 1 1 1 1 1 1 1 0 0 $j $j $i 10000 10000 0 1 10000 {}" >> $bad_start_male
	done
done
```
