## Supplementary material for "Establishment of local adaptation in partly self-fertilizing populations": Scripts for running the simulator and processing the raw data, together with the complete output necessary for plotting using Mathematica notebook.: compare_simulations_self_varied.R

```
remove(list=ls())
setwd("~/Desktop/incoming_projects/mating_system/simulation/selfing_sim/processed_data_selfing_varied/")
library(dplyr)
library(ggplot2)
library(scales)

sPar=0.01;

# Note that whenever sigma is loaded it is converted to population inbreeding coefficient
#================== Viability selection ==================
# Start in good patch
df <-read.table(file = './good_via_F.out',header = FALSE, stringsAsFactors = FALSE)
colnames(df) <- c("V1_AA","V1_Aa","V1_aa","V2_AA","V2_Aa","V2_aa",
                  "WF1_AA","WF1_Aa","WF1_aa","WF2_AA","WF2_Aa","WF2_aa",
                  "WM1_AA","WM1_Aa","WM1_aa","WM2_AA","WM2_Aa","WM2_aa",
                  "wF1_A","wF1_a","wM1_A","wM1_a",
                  "wF2_A","wF2_a","wM2_A","wM2_a",
                  "mP1","mP2","mS1","mS2","sigma",
                  "N1","N2","mu1","mu2","run_nb","rndSeed",
                  "P_est","P_time")
df <- df %>% select(V1_AA,V1_Aa,V1_aa,V2_AA,V2_Aa,V2_aa,sigma,mP1,mP2,mS1,mS2,P_est) %>% na.omit
df$sigma <- df$sigma/(2-df$sigma);

df_plot <- df %>% group_by(sigma, V1_Aa, mS1) %>%  
  summarise(mean_Pest=mean(P_est), sd_Pest=sd(P_est), .groups='drop')
df_plot$V1_Aa <- round((df_plot$V1_Aa-1)/sPar, digits = 4)
write.csv(df_plot, './good_via_selfing_varied_processed.csv', row.names = FALSE)
ggplot(df_plot, aes(x=sigma, y=mean_Pest, col=as.factor(mS1))) + scale_y_continuous(trans='log10')  + geom_point()


# Start in bad patch
df <-read.table(file = './bad_via_F.out',header = FALSE, stringsAsFactors = FALSE)
colnames(df) <- c("V1_AA","V1_Aa","V1_aa","V2_AA","V2_Aa","V2_aa",
                  "WF1_AA","WF1_Aa","WF1_aa","WF2_AA","WF2_Aa","WF2_aa",
                  "WM1_AA","WM1_Aa","WM1_aa","WM2_AA","WM2_Aa","WM2_aa",
                  "wF1_A","wF1_a","wM1_A","wM1_a",
                  "wF2_A","wF2_a","wM2_A","wM2_a",
                  "mP1","mP2","mS1","mS2","sigma",
                  "N1","N2","mu1","mu2","run_nb","rndSeed",
                  "P_est","P_time")
df <- df %>% select(V1_AA,V1_Aa,V1_aa,V2_AA,V2_Aa,V2_aa,sigma,mP1,mP2,mS1,mS2,P_est) %>% na.omit
df$sigma <- df$sigma/(2-df$sigma);

df_plot <- df %>% group_by(sigma, V1_Aa, mS1) %>%  
  summarise(mean_Pest=mean(P_est), sd_Pest=sd(P_est), .groups='drop')
df_plot$V1_Aa <- round((df_plot$V1_Aa-1)/sPar, digits = 4)
write.csv(df_plot, './bad_via_selfing_varied_processed.csv', row.names = FALSE)
ggplot(df_plot, aes(x=sigma, y=mean_Pest, col=as.factor(mS1))) + scale_y_continuous(trans='log10') + geom_point()
```
